# *Enterobacterales* plasmid sharing amongst human bloodstream infections, livestock, wastewater, and waterway niches in Oxfordshire, UK

**DOI:** 10.1101/2022.05.06.490774

**Authors:** William Matlock, Samuel Lipworth, Kevin K. Chau, Manal Abu Oun, Leanne Barker, James Kavanagh, Monique Andersson, Sarah Oakley, Marcus Morgan, Derrick W. Crook, Daniel S. Read, Muna Anjum, Liam P. Shaw, Nicole Stoesser, REHAB Consortium

## Abstract

Plasmids enable the dissemination of antimicrobial resistance (AMR) in common *Enterobacterales* pathogens, representing a major public health challenge. However, the extent of plasmid sharing and evolution between *Enterobacterales* causing human infections and other niches remains unclear, including the emergence of resistance plasmids. Dense, unselected sampling is highly relevant to developing our understanding of plasmid epidemiology and designing appropriate interventions to limit the emergence and dissemination of plasmid-associated AMR. We established a geographically and temporally restricted collection of human bloodstream infection (BSI)-associated, livestock-associated (cattle, pig, poultry, and sheep faeces, farm soils) and wastewater treatment work (WwTW)-associated (influent, effluent, waterways upstream/downstream of effluent outlets) *Enterobacterales*. Isolates were collected between 2008-2020 from sites <60km apart in Oxfordshire, UK. Pangenome analysis of plasmid clusters revealed shared “backbones”, with phylogenies suggesting an intertwined ecology where well-conserved plasmid backbones carry diverse accessory functions, including AMR genes. Many plasmid “backbones” were seen across species and niches, raising the possibility that plasmid movement between these followed by rapid accessory gene change could be relatively common. Overall, the signature of identical plasmid sharing is likely to be a highly transient one, implying that plasmid movement might be occurring at greater rates than previously estimated, raising a challenge for future genomic One Health studies.

**Funding:** This study was funded by the Antimicrobial Resistance Cross-council Initiative supported by the seven research councils and the NIHR, UK.

## Introduction

*Enterobacterales* are found both in human niches (e.g., hospital patients(1,2) and wastewater(3)) and non-human niches (e.g., livestock-associated(4,5) and waterways(6)). In recent decades, widespread carriage of antimicrobial resistance (AMR) genes has complicated the treatment of *Enterobacterales* infections(7,8). The dissemination of AMR genes between *Enterobacterales* occurs in a ‘Russian-doll’-style hierarchy of nested, mobilisable genetic structures(9): genes not only move between bacterial hosts on mobilisable or conjugative plasmids but can also be transferred within and between plasmids and chromosomes by smaller mobile genetic elements (MGEs) such as insertion sequences(10,11). Despite gene gain/loss events, many plasmids have been shown to have a persistent structure encoding replication and transfer machinery(12,13).

Many plasmids can transfer between species and are seen across different niches(14) but the extent to which they are shared between human and non-human niches remains poorly understood. Previous studies investigating this topic have often been limited in size given the genetic diversity in these niches(15,16), and/or restricted to single species(17) or drug-resistant isolates(18), or are systematic studies, pooling geographically/temporally disparate samples (19,20). Further, fragmented genome assemblies in many cases make recovering complete plasmids, and other MGEs, impossible(21).

Instances of cross-niche transfer of plasmids are well-described, but the frequency of such events is poorly characterised. There are multiple instances where AMR genes have emerged from non-human niches and subsequently become major clinical problems in human *Enterobacterales* infections, highlighting the relevance of inter-niche transfer in AMR gene dissemination (e.g., *bla_CTX-M_*, *mcr-1*(22) and *bla*_NDM-1_(23)). In general, environmental bacteria are believed to be the original source of AMR genes that eventually become prevalent in clinical settings after transfer into clinical pathogens. However, we know little about natural rates of inter-niche transfer beyond these high-profile examples. It remains unclear how plasmids evolve within natural populations, meaning we understand little about the wider context in which AMR genes emerge and disseminate.

To explore *Enterobacterales* plasmid diversity and sharing across niches in a geographically and temporally restricted context, we studied hybrid assemblies (i.e., using both long and short reads) of large *Enterobacterales* isolate collections in Oxfordshire, UK, from (i) human bloodstream infections (BSI; 2008-2018), (ii) livestock-associated sources (faeces from cattle, pigs, poultry, sheep; surrounding environmental soils; [all 2017 except poultry 2019-2020], and (iii) wastewater treatment work (WwTW)-associated sources (influent, effluent, waterways upstream/downstream of effluent outlets; Oxfordshire, 2017).

## Results

Our dataset of *n*=3,697 plasmids from *n*=1,458 isolates (Fig. 1a, Table 1) contained bacteria from human bloodstream infections (BSI; *n*=1,880 plasmids from *n*=738 isolates), livestock-associated sources (cattle, pig, poultry, and sheep faeces, soils surrounding livestock farms; *n*=1,155 plasmids from *n*=512 isolates), and from wastewater treatment works (WwTW)-associated sources (influent, effluent, waterways upstream/downstream of effluent outlets; *n*=662 plasmids from *n*=208 isolates). All sampling sites were <60km apart (Fig. 1b) and timeframes overlapped (2008-2020; Fig. 1c). Isolates had a median 2 plasmids (IQR=1-4, range=0-16). Major *Enterobacterales* genera represented included: *n*=1,044 *Escherichia*, *n*=211 *Klebsiella*, *n*=125 *Citrobacter*, and *n*=63 *Enterobacter*.

**Fig. 1.**
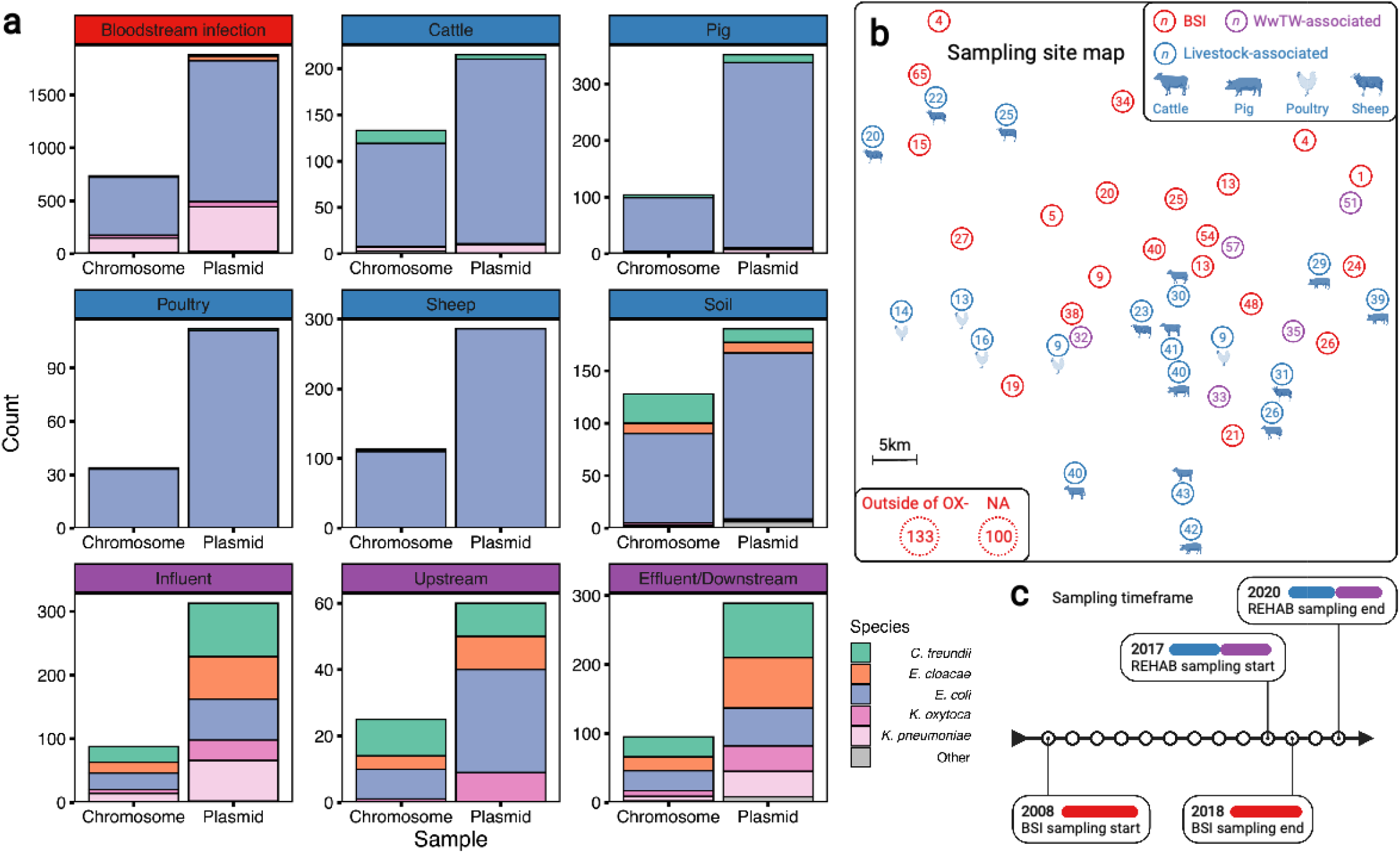
A diverse sample of geographically and temporally restricted *Enterobacterales*. **(a)** Number of chromosomes and plasmids by niche, stratified by isolate genus. **(b)** Map of approximate, relative distances between sampling sites, coloured by niche (human bloodstream infection [BSI], livestock-associated (cattle, pig, poultry, and sheep faeces, soils nearby livestock sites), and wastewater treatment work (WwTW)-associated sources (influent, effluent, waterways upstream/downstream of effluent outlets). Number in circles indicates how many of the *n*=1,458 isolates are from that location. **(c)** Sampling timeframe for BSI and REHAB (non-BSI) isolates.

**Table 1.** Isolate niche breakdown.

| **Table 1. Isolate niche breakdown.** |  |  |  |
| --- | --- | --- | --- |
| **Niche** | **Sample type(s)** | **No. isolates** | **No. plasmids** |
| Bloodstream infections (BSI) | Community, nosocomial, other healthcare associated infections | 738 | 1,880 |
| Livestock-associated | Cattle faeces | 133 | 215 |
|  | Sheep faeces | 113 | 286 |
|  | Pig faeces | 104 | 352 |
|  | Poultry faeces | 34 | 112 |
|  | Soil surrounding livestock farms | 128 | 190 |
| Wastewater treatment work (WwTW)-associated | Influent | 88 | 313 |
|  | Upstream waterways | 25 | 60 |
|  | Effluent/downstream waterways | 95 | 289 |
| **Total** |  | **1,458** | **3,697** |

Sampling niche was strongly associated with isolate genus (Fisher’s test, *p-*value<0.001; Table 2). *Klebsiella* isolates were disproportionately derived from BSI versus other niches (76% [161/212] versus 51% [738/1,458]). *Citrobacter* and *Enterobacter* were disproportionately derived from WwTW-associated versus other niches (52% [65/125] and 67% [42/63] versus 14% [208/1,458]). Chromosomal Mash trees (see Supplementary Methods) for the two most common species in the dataset, *E. coli* (72% [1,044/1,458]; see Fig. S1) and *K. pneumoniae* (11% [163/1,458]; Fig. S2) demonstrated intermixing of human and non-human isolates within clades, consistent with species-lineages not being structured by niche.

**Table 2.** Isolate genus breakdown.

| **Table 2. Isolate genus breakdown.** |  |  |  |  |  |  |
| --- | --- | --- | --- | --- | --- | --- |
| **Niche** | **Isolate genus** |  |  |  |  | **Total** |
|  | *Citrobacter* | *Enterobacter* | *Escherichia* | *Klebsiella* | *Other* |  |
| Bloodstream infections (BSI) | 6 | 11 | 547 | 161 | 13 | **738** |
| Livestock-associated | 54 | 10 | 433 | 14 | 1 | **512** |
| Wastewater treatment<br>work (WwTW)-associated | 65 | 42 | 64 | 37 | 0 | **208** |
| **Total** | **125** | **63** | **1,044** | **212** | **14** | **1,458** |

We contextualised our plasmids within known plasmid diversity using ‘plasmid taxonomic units’ (PTUs; using COPLA, see Supplementary Methods), designed to be equivalent to a plasmid ‘species’. We found 32% (1,193/3,697) of plasmids were unclassified, highlighting the substantial plasmid diversity within this geographically restricted dataset. In total, we found *n*=67 known PTUs, containing a median 9 plasmids (IQR=4-30, range=1-556), with the largest PTU-F_E_ (556/2,504), corresponding to F-type *Escherichia* plasmids.

### Near-identical plasmid sharing observed between human and livestock-associated *Enterobacterales*

We screened for near-identical plasmids shared across isolates by grouping those with a low Mash distance (*d*<0.0001) and highly similar lengths (longest plasmid ≤1% longer than shorter plasmids; see Supplementary Methods). We found *n*=225 near-identical groups of ≥2 members, recruiting 19% (712/3,697) plasmids. Bootstrapping accumulation curves for near-identical plasmid groups and singletons per the number of isolates (ACs; see Supplementary Methods), we revealed a highly ‘open’ accumulation (Heap’s parameter γ=0.97, Fig. S3) suggesting further isolate sampling would detect more unique plasmids approximately linearly. Restricted to BSI/livestock-associated isolates alone, we found similar curves for both niches (BSI γ=0.98, livestock-associated γ=0.94), suggesting they had similar levels of plasmid diversity.

Near-identical pairs of plasmids were most common, representing 71% (159/225) of groups (group size IQR=2-3, range=2-32). Plasmid members of near-identical groups represented multiple bacterial host STs (25% [56/225]), species (4% [9/225]), and genera (4% [9/225]), consistent with plasmids capable of inter-lineage/species/genus transfer. Further, 8% (17/225) of near-identical groups contained plasmids found across human BSIs and ≥1 other sampling niche (livestock-associated/WwTW-associated), suggesting inter-niche transfer (i.e., ‘cross-niche groups’; Fig. 2a). Within cross-niche groups, *n*=3/17 contained plasmids from multiple bacterial species (Fig. 2b), and most consisted of conjugative plasmids (*n*=5/17 conjugative, *n*=9/17 mobilisable, *n*=3/17 non-mobilisable; Fig. 2c). AMR genes were carried by plasmids in *n*=6/17 cross-niche groups (Fig. 2d), with *n*=5/6 of these groups containing ≥1 beta-lactamase protein encoding gene.

**Fig. 2.**
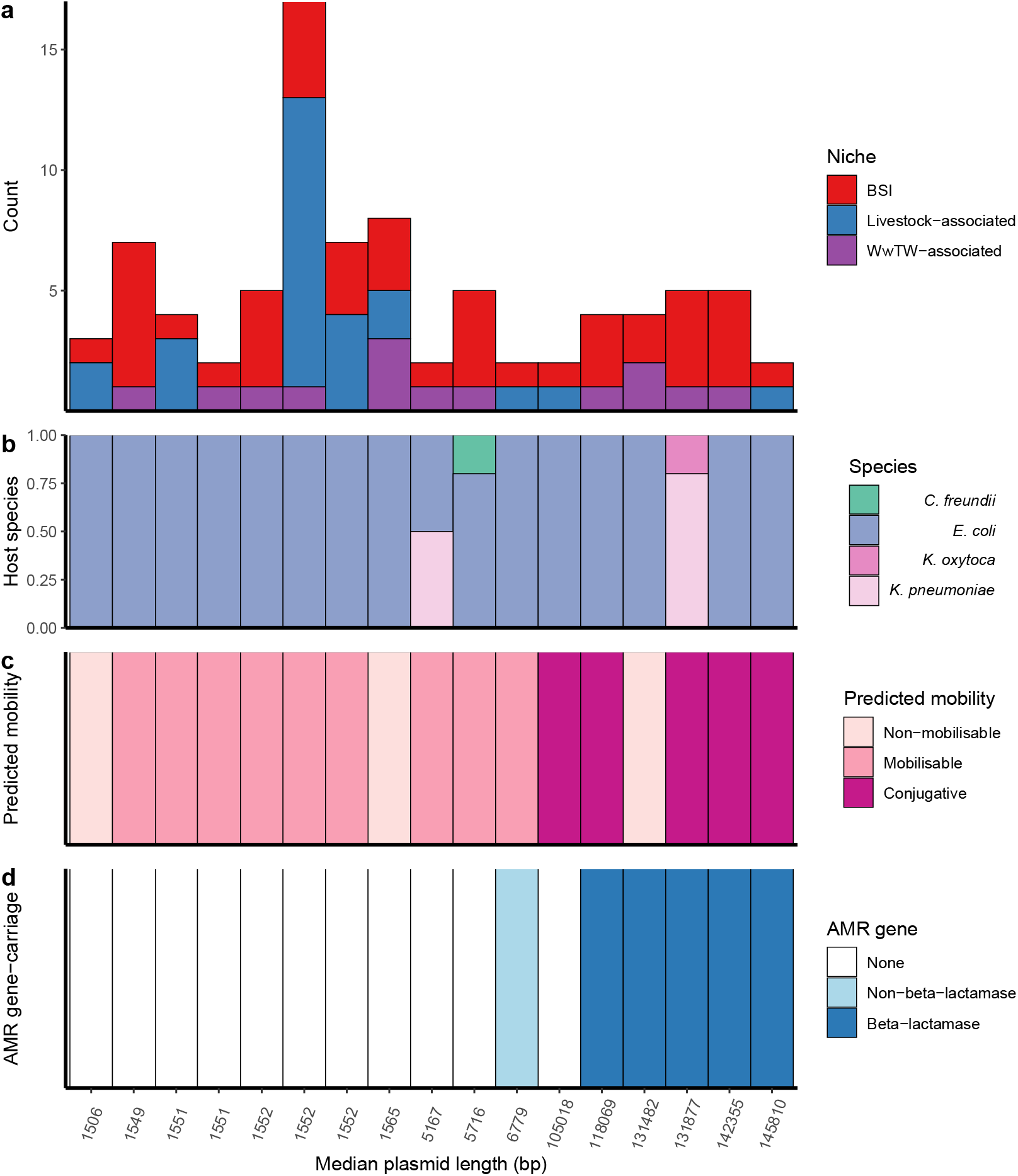
Cross-niche, near-identical plasmids. **(a)** Size of cross-niche, near-identical plasmid groups, coloured by niche (total *n*=84 plasmids). Median length (bp) of plasmids within groups increases from left to right. (**b)** Proportion of plasmid host species by group. **(c)** Predicted mobility of plasmid. **(d)** AMR gene carriage in plasmid.

Sharing between BSI and livestock-associated isolates was supported by 8/17 cross-niche groups (*n*=45 plasmids). Of these, *n*=2/8 contained non-mobilisable Col-type plasmids (one group contained BSI/pig/poultry/influent plasmids, and one group contained BSI/poultry plasmids); *n*=4/8 contained mobilisable Col-type plasmids (two groups contained BSI/pig plasmids, one group contained BSI/sheep plasmids, and one group contained BSI/cattle/pig/poultry/sheep/influent plasmids), of which one group contained BSI/pig plasmids carrying the AMR genes *aph(3’’)-Ib*, *aph(6)-Id*, *dfrA14*, and *sul2* (see Supplementary Methods). The remaining 2/8 groups contained conjugative FIB-type BSI/sheep plasmids. One group contained plasmids, carrying the AMR genes *aph(3’’)-Ib*, *aph(6)-Id*, *bla_TEM-1_*, *dfrA5*, *sul2*, and the other group contained plasmids carrying the MDR efflux pump protein *robA*.

The beta-lactamase *bla*_TEM-1_ was the most common AMR gene detected (8% of total AMR gene annotations [424/5402]; see Supplementary Methods). In terms of sequence length (bp), plasmids made up 3.1% of the overall dataset but 13.8% of the *bla*_TEM-1_ -carrying proportion. Of the plasmid clusters, 16% (39/247) carried *bla*_TEM-1_, and of these 9 clusters were seen in human BSI and at least one other niche. Plasmid clusters either variably or always carrying *bla*_TEM-1_ were strongly associated with BSI (p<0.01, Chi-squared test X^2^=8.19, 33/161 of BSI clusters containing *bla*_TEM-1_ vs. 5/86 for non-BSI clusters) and carried a higher number of other AMR genes (p<0.01, Wilcoxon text of *bla*_TEM-1_-plasmid clusters vs. others; see Fig. S4).

### Plasmid clustering reveals a diverse but intertwined population structure across niches

Near-identical plasmids shared across niches are a likely signature of recent transfer events, but we also wanted to examine the wider plasmid population structure. We therefore agnostically clustered all plasmids based on alignment-free sequence similarity (clusters were groups of *n*≥3 plasmids; see Supplementary Methods and Figs. S5-6). We defined *n*=247 plasmid clusters with median 5 members (IQR=3-10, range=3-123) recruiting 71% (2,627/3,697) of the plasmids. The remainder were either singletons (i.e., single, unconnected plasmids; 19% [718/3,697]) or doubletons (i.e., pairs of connected plasmids; 10% [352/3,697]). By bootstrapping *b*=1,000 ACs for plasmid clusters, doubletons, and singletons found against number of isolates sampled (Fig. S7; see Supplementary Methods), we estimated that the rarefaction curve had a Heap’s parameter γ=0.75, suggesting further isolate sampling would likely detect more plasmid diversity and clusters.

Of the plasmid clusters, *n*=69/247 (28%) plasmid clusters had ≥10 members, representing 50% (1,832/3,697) of all plasmids (Fig. 3a). 122/247 (49%) clusters contained BSI plasmids and plasmids from ≥1 other niche. This included 73/247 (30%) of clusters with both BSI and livestock-associated plasmids, representing *n*=38 unique plasmid replicon haplotypes (i.e., combinations of replication proteins) of which only 24% (9/38) were Col-type plasmids, which are often well-conserved and carry few genes(24). 72/247 (29%) of clusters contained both BSI, and influent/effluent/downstream plasmids, reflecting a route of *Enterobacterales* dissemination into waterways. In contrast, only 18/247 (7%) of clusters contained both BSI and upstream waterway plasmids, of which most (13/18 [72%]) also contained influent/effluent/downstream plasmids.

**Fig. 3.**
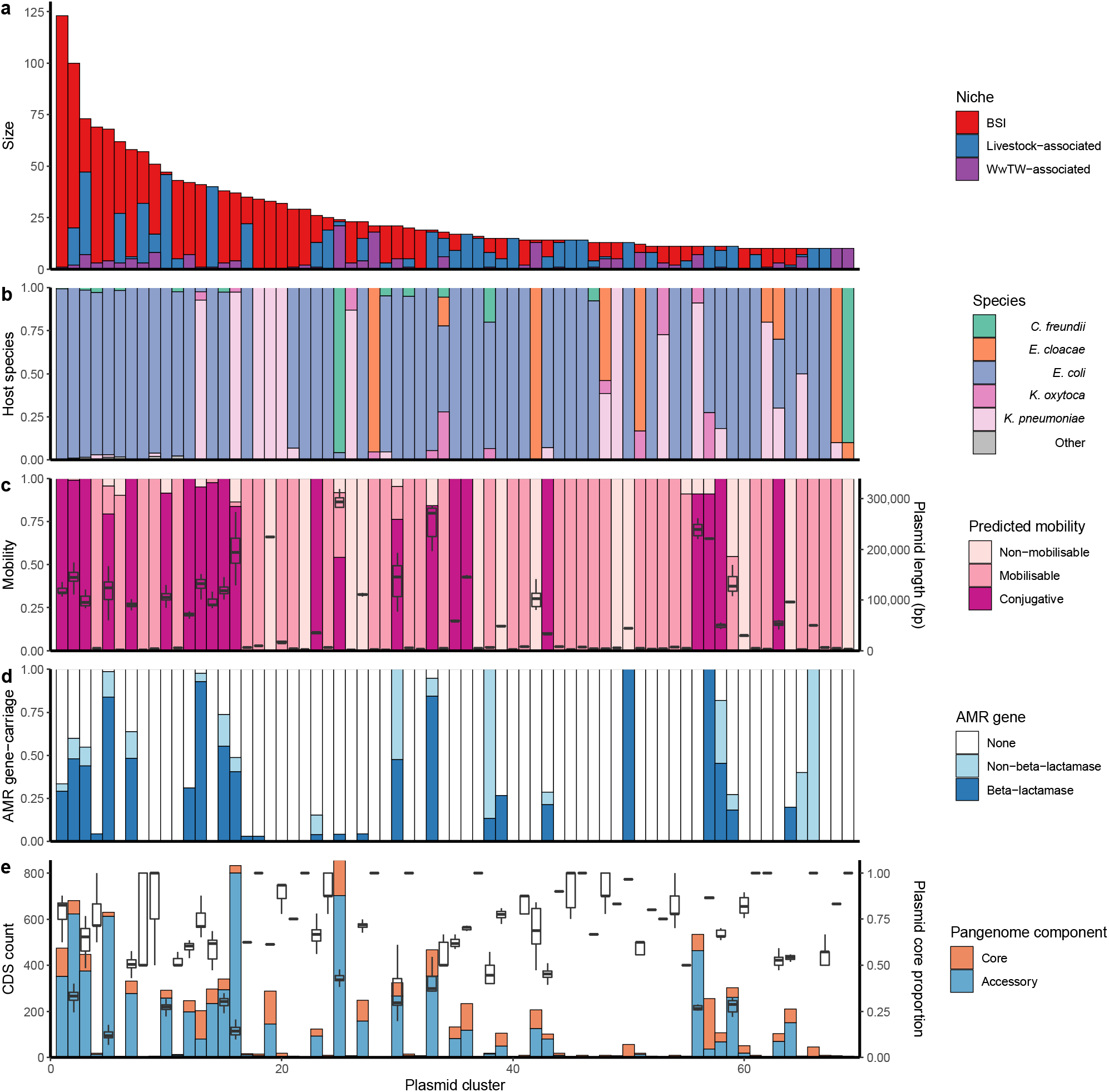
Genetically similar plasmids share between niches. **(a)** Size of plasmid clusters with at least 10 members, coloured by niche. Size of clusters decreases from left to right. (**b)** Proportion of plasmid host species by cluster. **(c)** Plasmid mobility class and size: Left hand axis shows proportion of plasmids with a predicted mobility class by cluster. Right hand axis shows plasmid length boxplots by cluster. **(d)** Proportions of AMR gene carriage by cluster. **(e)** Plasmid core and accessory genomes: Left hand axis shows the count of core and accessory coding sequences (CDS) by cluster. Right hand axis shows plasmid core gene proportion (i.e., plasmid core CDS/total plasmid CDS) boxplots by cluster.

Overall, plasmid clusters scored high homogeneity (*h*) but low completeness (*c*) with respect to biological and ecological characteristics (non-putative PTUs [*h*=0.99, *c*=0.66]; replicon haplotype [*h*=0.92, *c*=0.69]; bacterial host ST [*h*=0.84, *c*=0.14] in Fig. 3b; predicted mobility [*h*=0.93, *c*=0.20] in Fig. 3c). This indicated that clustered plasmids often had similar characteristics, but the same characteristics were often observed in multiple clusters. The imperfect homogeneity is to be anticipated as replicon haplotypes and mobilities can vary within plasmid families, and plasmid families can have diverse host ranges(14).

Plasmids carrying AMR genes were found in 21% (52/247) of the plasmid clusters (i.e., ‘antimicrobial resistance gene (ARG)-carrying clusters’), representing *n*=550 plasmids (Fig. 3d). Of the ARG-carrying clusters, 92% (48/52) contained at least one beta-lactamase-carrying plasmid (*n*=437 plasmids in total). AMR genes were present in a median proportion 67% of ARG-carrying cluster members (IQR=28-100%, range=3-100%). This highlights that AMR genes are not necessarily widespread on genetically similar plasmids and can be potentially acquired multiple different times through the activity of smaller MGEs (e.g., transposons) or recombination. For example, cluster 12 was a group of *n*=42 conjugative, PTU-F_E_ plasmids found in BSI, wastewater, and waterways. Of these, 31% (13/42) carried the AMR gene *bla_TEM-1_*, and in a range of genetic contexts: *n*=9/13 *bla_TEM-1_* genes were found within Tn*3* and *n*=4/13 were carried without a transposase, of which *n*=2/4 were found with the additional AMR genes *aph(6)-Id*, *aph(3’’)-Ib*, and *sul2*. AMR genes were disproportionately carried by F-type plasmids (61% [337/550] ARG-carrying cluster plasmids versus 34% [891/2627] of the total clustered plasmids), underlining the known role of F-type plasmids in AMR gene dissemination(13).

### An intertwined ecology of plasmids across human and livestock-associated niches

Plasmids can change their genetic content, particularly when subject to new selective pressures(25,26). Many plasmids have a structure with a ‘backbone’ of conserved core genes and a ‘cargo’ of variable accessory genes(12,13,27). We wanted to explore evidence for cross-niche plasmids with minimal mutational evolution in a shared backbone (compatible with ~years of evolutionary separation) but variable accessory gene repertoires.

We first conducted a pangenome-style analysis (see Supplementary Methods) on the *n*=69/247 plasmid clusters with ≥10 members. For each cluster, we determined “core” (genes found in ≥95% of plasmids) and “accessory” gene repertoires (found in <95% of plasmids). Within clusters, we found median 9 core genes (IQR=4-53, range=0-219), and median 9 accessory genes (IQR=3-145, range=0-801) (Fig. 3e). Core genes comprised a median proportion 42.2% of the total pangenome sizes (IQR=20.9-66.7%). At an individual plasmid level, core genes shared by a cluster comprised a median proportion 62.5% of each plasmid’s gene repertoire (IQR=37.4-83.3%; Fig. 3e). Putatively conjugative plasmids carried a significantly higher proportion of accessory genes in their repertoires than mobilisable/non-mobilisable plasmids (Kruskal-Wallis test [*H*(2)=193.01, *p*-value<0.001] followed by Dunn’s test).

Using multiple sequence alignments of the core genes within each cluster, we produced maximum likelihood phylogenies (see Supp. File 1 and Supplementary Methods). For this step, we only considered the *n*=62/69 clusters where each plasmid had ≥1 core gene. With the *n*=27/62 clusters that contained both BSI and livestock-associated plasmids, we measured the phylogenetic signal for plasmid sampling niche using Fritz and Purvis’ *D* (see Table S1 and Supplementary Methods). The analysis indicated that the evolutionary history of plasmid clusters is neither strictly segregated by sampling niche nor completely intermixed, but something intermediate.

Alongside the core gene phylogenies, we generated gene repertoire heatmaps (example cluster 2 in Fig. 4a-b; all clusters and heatmaps in Supp. File 1). By visualising the genes in a consensus synteny order (see Supplementary Methods), the putative backbone within each plasmid cluster is shown alongside its accessory gene and transposase repertoire. This highlights how plasmids might gain/lose accessory functions within a persistent backbone. Log-transformed linear regression revealed a significant relationship between Jaccard distance of accessory genes presence against core gene cophenetic distance (*y*=0.080log(*x*)+0.978, *R*^2^=0.47, *F*(1,52988)=4.75e4, *p*-value <0.001; see Fig. S8 and Supplementary Methods).

**Fig. 4.**
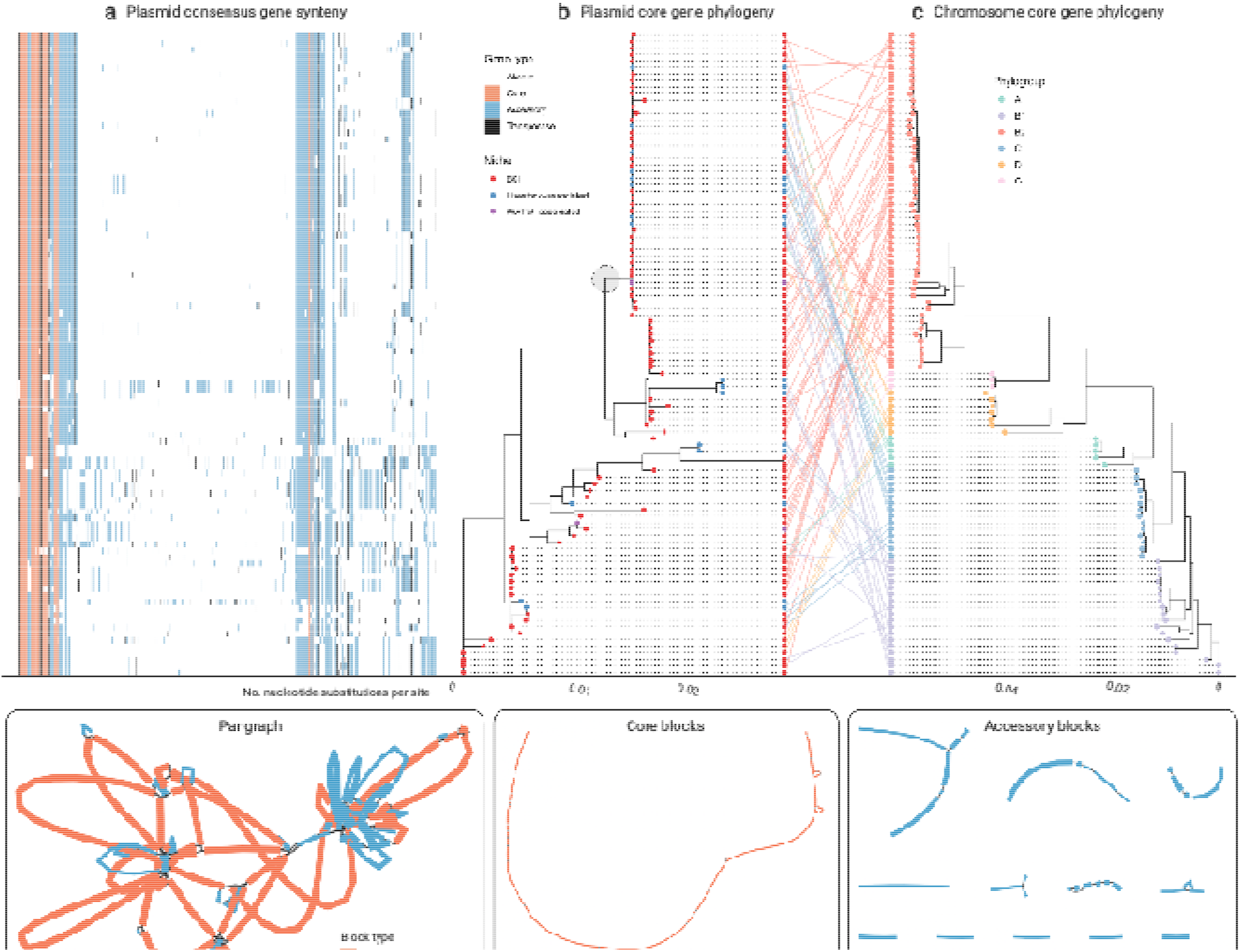
Cluster 2 plasmid and host evolution. **(a)** Consensus gene ordering for plasmid cluster 2, coloured by gene type (total *n*=99 plasmids; *n*=1 *S. enterica* isolate omitted). Genes are coloured by core, accessory, or transposase. **(b)** Plasmid core gene phylogeny with tips coloured by sampling niche. The grey circle highlights the clade of *n*=44 plasmids which were further analysed. **(c)** Plasmid host chromosome core gene phylogeny with tips coloured by sampling niche. Plasmid and host phylogeny tips are connected in a ‘tanglegram’ which connects pairs of plasmids and chromosomes from the same isolate. **(d)** Visualisation of the pangraph for *n*=44 plasmids in the grey-circled clade in (b). Blocks are coloured by presence in plasmids. **(e)** Core blocks (found in at least 95% of the *n*=44 plasmids). **(f)** Accessory blocks (found in less than 95% of the *n*=44 plasmids).

### Plasmid evolution between human and livestock-associated niches is not structured by bacterial host

Alongside vertical inheritance, conjugative and mobilisable plasmids are capable of inter-host transfer, crossing between bacterial lineages, species, up to phyla(14). Phylogenetic analysis can determine whether plasmid evolution between BSI and livestock-associated niches is driven by host clonal expansion or other means, as well as allow us to explore the early emergence of AMR gene carrying plasmids.

As a detailed example, we evaluated the largest plasmid cluster containing both human and livestock-associated plasmids (cluster 2, *n*=100 members). All plasmids carried at least one F-type replicon and were all putatively conjugative, with 75% (75/100) and 25% (25/100) assigned PTU-F_E_ and a putative PTU, respectively. Further, 48% (48/100) plasmids carried *bla_TEM-1_*, and 51% (51/100) carried >1 AMR gene. All host chromosomes were *E. coli* except OX-BSI-481_2 (*S. enterica* ST 2998; hereon omitted from the analysis). The *n*=99 *E. coli* isolates represented six phylogroups: A (5/99), B1 (18/99), B2 (52/99), C (14/99), D (7/99), and G (3/99; see Supplementary Methods).

Figure 4b-c shows the plasmid core gene phylogeny (*T*_plasmid_) and the *E. coli* host core gene phylogeny (*T*_chromosome_). The *E. coli* phylogeny was structured by six clades corresponding to the six phylogroups (see Supplementary Methods). We found low congruence between the plasmid core-gene phylogeny and the chromosomal core-gene phylogeny as seen in the central ‘tanglegram’ (i.e., lines connecting pairs of plasmid and chromosome tips from the same isolate). Additionally, we calculated a Robinson-Foulds distance *RF*(*T*_plasmid_, *T*_chromosome_)=162, reflecting a high number of structural differences between the phylogenies (see Supplementary Methods). There was some evidence of plasmid structuring by niche (Fritz and Purvis’ *D*=0.24; see Supplementary Methods).

Within the plasmid phylogeny, there was a clade of *n*=44 plasmids (support 100%; circled in grey in Figure 4b) containing both BSI and livestock-associated plasmids, which were within median 4 core gene SNPs of each other (IQR=2-8, range=0-59). Estimating plasmid evolution at an approximate rate of one SNP per year (see Supplementary Methods) would give a median time to most recent common ancestor of the backbone at approximately 4 years prior to sampling, consistent with recent movement between human and livestock-associated niches. This plasmid clade was mainly present in phylogroup B2 (20/44), but also A (3/44), B1 (9/44), C (8/44), and D (4/44), suggesting plasmid movement. Further, 77% (34/44) of plasmids within the clade carried *bla_TEM-1_* (BSI: 25/34, Livestock-associated: 8/34, WwTW-associated: 1/34), and 82% (36/44) carried ≥1 AMR gene, highlighting the role of plasmids in cross-niche dissemination of AMR.

To examine the evolution of entire plasmid sequences within the clade, we represented all *n*=44 plasmids as a ‘pangraph’ (Figure 4d; see Supplementary Methods). Briefly, pangraph converts input sequences into a consensus graph, where each sequence is a path along a set of homologous sequence alignments i.e., ‘blocks’, which in series form ‘pancontigs’. Filtering for ‘core blocks’ (i.e., those found in ≥95% plasmids), we found 4 pancontigs (40 blocks total), with the longest 98,269bp (total length 125,369bp), indicating a putative plasmid backbone (Fig. 4e). Then, filtering for ‘accessory blocks’ (i.e., those found in <95% plasmids), we found 18 pancontigs (39 blocks total), with median length 2,380bp (total length 63,753bp), forming the accessory gene repertoire (Fig. 4f). This points to a persistent plasmid backbone structure with loss/gain events at particular ‘hotspots’ as well as rearrangements.

## Discussion

Sharing of plasmids between different niches is normally focused on those carrying AMR genes that are of particular current clinical concern, such as extended-spectrum beta-lactamase (ESBL) or carbapenemase genes, meaning we lack information on the vast ‘denominator’ of background plasmid sharing, and on the dissemination of other AMR genes which are now widespread in clinical isolates and from which important insights might be gained. By analysing a dataset of *n*=3,697 systematically collected *Enterobacterales* plasmids sampled from human BSI, livestock- and WwTW-associated sources in a geographically and temporally restricted context, we found evidence supporting significant plasmid dissemination across niches, putting those which carry AMR genes of current major clinical concern into context. We found 225 instances of shared, near-identical plasmid groups, 25% of which were found across multiple bacterial STs, 4% across multiple bacterial species, and 8% in both human BSI and ≥1 non-BSI niche. Beyond this near-identical sharing, we analysed ‘clusters’ of plasmids and found that 73/247 clusters contained plasmids seen in both human BSIs and other contexts. Approximately a fifth (52/247) of plasmid clusters contained plasmids carrying AMR genes (*n*=550 plasmids). Our results suggest the need for broad, unselected, and detailed sampling frames to fully understand plasmid diversity and evolution, and to evaluate the “One Health” risk of AMR associated with plasmid-sharing across niches.

Whilst many plasmid clusters were strongly structured by host phylogeny and isolate source, some plasmids from human BSIs were highly genetically related to those in other niches, including livestock. However, not all of these carried AMR genes. Our results highlight the potential routes for transfer that exist through similar plasmids. However, recovering these instances of putative sharing is a sampling challenge. Accumulation curve analyses suggested increasing the size of our dataset would have led to further near-identical matches at an approximately linear rate, meaning even a dataset of this size captures only a small fraction of the true extent of plasmid sharing between human clinical and other non-human/clinical niches. This presents a challenge for designing appropriately powered studies. Had we only sampled *n*=100 livestock-associated isolates (i.e., around 20% of our actual sample), there was only a 39% chance that we would have detected ≥5 matches with BSI plasmids (Fig. S9).

Understanding the evolutionary history, distribution, and epidemiology of well-known genes in environmental plasmids may offer insights into the future trajectories of more recently emerged genes. For example, the first plasmid-encoded beta-lactamase to be described was *bla*_TEM-1_, identified in 1965 in an *E. coli* isolate in Greece(28) and now widely prevalent in *Enterobacterales*(29). *bla*_TEM-1_ has a narrow spectrum of activity and is now less clinically concerning than newer genes which mediate broad-spectrum resistance, but in our dataset *bla*_TEM-1_ was strongly associated with plasmid clusters seen in BSI and with the carriage of other AMR genes. *bla*_TEM-1_ may continue to play an important role in the spread of AMR-carrying plasmids which can transfer recently emerged genes, and similarities in its association with plasmids and other smaller transposable mobile genetic elements may reflect the future trajectory of other AMR genes of more recent clinical concern such as ESBLs and carbapenemases.

Given that plasmids observed in BSI isolates represent small proportion of human *Enterobacterales* diversity, many more sharing events may occur in the human gut(30) which we only sampled incompletely using wastewater influent as a proxy. The human colon contains around 10^14^ bacteria(31), with large ranges of *Enterobacteriaceae* abundance. Further, even small numbers of across-niche sharing events, such as transfer events of important AMR genes from species-to-species or niche-to-niche, may have significant clinical implications, as has been seen with several important AMR genes globally. Future studies need to carefully consider the limitations of sampling frames in detecting any genetic overlap, given both substantial diversity and the effects of niches and geography(11,16).

By examining plasmid relatedness compared to bacterial host relatedness, we demonstrated that cross-niche plasmid spread is not driven by clonal lineages. Using a pangenome-style analysis, we showed that plasmids can share sets of near-identical core genes alongside diverse accessory gene repertoires. While plasmids with more distantly related core genes tended to have dissimilar accessory gene content, plasmids with more closely related core genes shared a wide range of accessory gene content. This would be consistent with a hypothesis of persistent ‘backbone’ structures gaining and losing accessory functions as they move between hosts and niches. We suggest that this mode of transfer might be worth considering. Evolutionary models for plasmids which can accommodate well-conserved backbone evolution alongside accessory structural changes and gain/loss events are urgently needed. Estimating plasmid evolutionary rates remains a challenge, with little known about appropriate values for mutation rates in plasmids, and even less for non-mutational processes such as gene gain/loss.

Our study had several limitations. Our non-BSI isolates were not as temporally varied as the BSI isolates, meaning we could not fully explore temporal evolution. Isolate-based methodologies are limited in evaluating the true diversity of the niches sampled; composite approaches including metagenomics might shed additional insight in future studies. Further, the exact source of an isolate is poorly defined for wastewater/waterway isolates as they act as a confluence of multiple sources, although they represent important niches in their own right. We only analysed plasmids from complete genomes i.e., where the chromosome and all plasmids were circularised, meaning we disregarded ~23% and ~33% of BSI and non-BSI assemblies, respectively. The exclusive use of complete assemblies was to ensure full plasmid sequences could be examined in their full genomic context. We only focused on plasmids as horizontally transmissible elements here; detailed study of other smaller mobile genetic elements across-niches would represent interesting future work. We have also investigated a limited subset of *Enterobacterales*: plasmid sharing likely extends to other bacterial hosts not investigated here. Lastly, our isolate culture methods for livestock-associated samples may not have been as sensitive for the identification of *Klebsiella* spp. as for other *Enterobacterales* such as *Escherichia*, as we did not use enrichment and selective culture on Simmons citrate agar with inositol(32).

In conclusion, this study presents to our knowledge the largest evaluation of systematically collected *Enterobacterales* plasmids across human and non-human niches within a geographically and temporally restricted context. Plasmids can clearly disseminate between niches, although this dynamic likely varies by cluster; the overall number of near-identical plasmid groups identified across niches consistent with recent transfer events was 8% (17/225) and influenced by sample size. We demonstrate a likely intertwined ecology of plasmids across human and non-human niches, where different plasmid clusters are variably but incompletely structured and putative ‘backbone’ plasmid structures can rapidly gain and lose accessory genes following cross-niche spread. Future “One Health” studies require dense and unselected sampling, and complete/near-complete plasmid reconstruction, to appropriately understand plasmid epidemiology across niches.

## Materials and Methods

### Livestock-associated isolates

*n*=247 *Enterobacterales* isolates from farm-proximate soils and poultry faeces (*n*=19 farms; *n*=5 cattle, *n*=4 pig, *n*=5 poultry, *n*=5 sheep) were collected and sequenced for this study in 2017-2020. DNA extraction and sequencing was performed as in Shaw *et al*., 2021(11). Genomes were hybrid assemblies reconstructed using Unicycler(33) (v. 0.4.4; default hybrid assembly parameters except min_component_size 500 and --min_dead_end_size 500). Only complete assemblies (plasmids and chromosomes) were considered (*n*=162/247).

### BSI isolates

Sequenced Human BSI Enterobacterales isolates from patients presenting to *n*=4 hospitals within Oxfordshire, UK, September 2008-December 2018, as described in Lipworth *et al*., 2021(34) were also included. Although all patients were sampled in Oxfordshire, a total of *n*=505/738 patients resided in Oxfordshire, *n*=133/738 in surrounding counties, and *n*=100/738 had location information omitted. Only complete assemblies (*n*=738/953 total assembled) were considered.

### Other livestock-associated and WwTW-associated isolates

*Enterobacterales* isolates from faeces from the *n*=14 non-poultry farms and wastewater influent, effluent, and waterways upstream/downstream of effluent outlets surrounding *n*=5 WwTWs, across 3 seasonal timepoints in 2017 (as in (11)) were included. Only complete assemblies (*n*=558/827 total assembled) were considered.

### Statistical analysis and bioinformatics

Chromosome sequence types (STs) were determined with mlst (v. 2.19.0; see Supplementary Methods). To generate accumulation curves (ACs), new plasmid diversity was recorded for each isolate sampled randomly, without replacement. A bootstrapped average of *b*=1,000 ACs was used to estimate a Heap’s parameter (γ) by fitting a linear regression to log-log transformed data see (Supplementary Methods). We adopted three approaches to plasmid classification, using COPLA to classify plasmids into broad plasmid taxonomic units (PTUs), and also grouping and clustering plasmids into smaller clusters using alignment-free distances (see Supplementary Methods). Within plasmid clusters, we identified core genes with Panaroo (v. 1.2.9), aligned them with mafft (v7.407) and produced trees with IQ-tree (v. 2.0.6). Plots were primally produced using the R library ggplot2, with additional graphics in BioRender. More information can be found in the Supplementary Methods.

## Data availability

Study metadata is provided in Table S2. Accessions for poultry and environmental soil isolate reads are given in Table S3, and assemblies will shortly be made available on NCBI. Accessions for existing BSI and REHAB reads and assemblies can be found in Lipworth *et al*., 2021(34) (BioProject PRJNA604975) and Shaw *et al*., 2021(11) (BioProject PRJNA605147) respectively.

## Code availability

Analysis scripts can be found in the GitHub repository https://github.com/wtmatlock/oxfordshire-overlap.

## REHAB Consortium

Manal AbuOun^2^, Muna F. Anjum^2^, Mark J. Bailey^3^, Brett H^8^, Mike J. Bowes^3^, Kevin K. Chau^1^, Derrick W. Crook^1,6,7^, Nicola de Maio^1^, Nicholas Duggett^2^, Daniel J. Wilson^1,9^, Daniel Gilson^2^, H. Soon Gweon^3,4^, Alasdair Hubbard^10^, Sarah J. Hoosdally^1^, William Matlock^1^, James Kavanagh^1^, Hannah Jones^2^, Timothy E. A. Peto^1,6,7^, Daniel S. Read^3^, Robert Sebra^5^, Liam P. Shaw^1^, Anna E. Sheppard^1,6^, Richard P. Smith^2^, Emma Stubberfield^2^, Nicole Stoesser^1,6,7^, Jeremy Swann^1^, A. Sarah Walker^1,6,7^, Neil Woodford^11^

^1^ Nuffield Department of Medicine, University of Oxford, Oxford, UK

^2^ Animal and Plant Health Agency, Weybridge, Addlestone, UK

^3^ UK Centre for Ecology & Hydrology, Wallingford, UK

^4^ University of Reading, Reading, UK

^5^ Icahn Institute of Data Science and Genomic Technology, Mt Sinai, NY, USA

^6^ NIHR HPRU in healthcare-associated infection and antimicrobial resistance, University of Oxford, Oxford, UK

^7^ NIHR Oxford Biomedical Research Centre, University of Oxford, Oxford, UK

^8^Thames Water Utilities, Clearwater Court, Vastern Road, Reading, UK

^9^Wellcome Trust Centre for Human Genetics, University of Oxford, Roosevelt Drive, Oxford, UK

^10^Department of Tropical Disease Biology, Liverpool School of Tropical Medicine, Liverpool, UK

^11^Antimicrobial Resistance and Healthcare Associated Infections (AMRHAI) Reference Unit, National Infection Service, Public Health England, London, United Kingdom

