## Supplementary Methods for "*Enterobacterales* plasmid sharing amongst human bloodstream infections, livestock, wastewater, and waterway niches in Oxfordshire, UK"

**Taxonomic assignment**

Chromosome sequence types (STs) were determined with mlst^1^ (v. 2.19.0; PubMLST database^2^). For the *n*=11/1,458 chromosomes which could not be typed with mlst, species were determined with the PubMLST ‘species ID’ web-tool^3^, for which all had a support=100, except for *L. nimipressuralis* (support=83). Of these, *n*=5/11 were from BSI, *n*=4/11 from livestock, and *n*=2/11 from effluent/downstream of WwTWs. From the BSI isolates, we also included n=2 *Aeromonas* spp., a non-*Enterobacterales* genus from the wider *Gammaproteobacteria* class.

**Chromosome trees**

Trees for *E. coli* and *K. pnemoniae* chromosomes were produced using Mashtree^4^ on ‘accurate’ mode (--mindepth 0 --numcpus 12).

**PTU classification**

Plasmids were assigned a plasmid taxonomic unit (PTU) using COPLA^5^ (default parameters except -t circular, -k Bacteria, -p Pseudomonadota, -c Gammaproteobacteria, and

-o Enterobacterales)^6^. COPLA compares query plasmids to a database of PTU reference plasmids, assigning a PTU when both (i) the ANI>0.7 along 50% of the length of the smallest plasmid in the comparison, and (ii) a graph-neighbouring condition to existing PTU clusters is satisfied. The COPLA reference database contains over 10,000 curated, non-redundant plasmids retrieved from the 84th NCBI RefSeq database in 2017^7^. We contextualised our plasmids within known plasmid diversity using COPLA to determine each plasmid’s ‘plasmid taxonomic unit’ (PTU; see Supplementary Methods), which is designed to be equivalent to a ‘species’ concept for plasmids^5^. Briefly, COPLA classifies query plasmids based on average nucleotide identity (ANI) against a non-redundant reference plasmid database where most plasmids have been assigned to a reference PTU^40^. Within our sample, 64% (2,369/3,697) plasmids were assigned a PTU and 4% (135/3,697) a putative PTU (i.e., the query plasmid was clustered with 3 unclassified reference plasmids). This is consistent with a previous COPLA analysis of 1,000 *Enterobacterales* plasmids which found that 63% were classified into a PTU ^5^. The remaining 32% (1,193/3,697) of plasmids were unclassified (i.e., connected set with less than 4 plasmids) highlighting the previously unsampled plasmid diversity within our dataset. In total, we found *n*=67 known PTUs, containing a median 9 plasmids (IQR=4-30, range=1-556), where the largest assigned PTU (556/2,504) was PTU-F_E_, corresponding to F-type *Escherichia* plasmids^8,9^. The proportion of unclassified plasmids was higher in environmental/livestock samples (33%; 385/1,155) versus BSI samples (26%; 485/1,880), emphasising the underrepresentation of non-human plasmids in reference plasmid databases.

**Plasmid annotation**

All plasmids were annotated with Prokka^1080^ (v. 1.14.5) with default parameters. For replicon typing, Abricate^11^ (v. 1.0.0) was used with the PlasmidFinder^12^, ISfinder^13^, and BacMet^14^ databases with default parameters and output filtered for 80% minimum coverage. For annotating AMR genes, NCBI Antimicrobial Resistance Gene Finder (AMRFinderPlus)^15^ (v. 3.10.18) was used with default parameters. For putative plasmid mobilities, we used MOB-typer from MOB-suite^16^ (v. 3.03) with default parameters, which predicts mobility based on annotations of MOB-typer predicts mobility based on of annotations of relaxase (*mob*), mating pair formation (MPF) complex, and *oriT* genes. Briefly, a plasmid is putatively labelled conjugative if it has both relaxase and MPF, mobilisable if it has either relaxase or *oriT* but no MPF, and non-mobilisable if it has no relaxase and *oriT*.

**Near-identical plasmid screening**

Groups of near-identical plasmids were detected as connected components in a plasmid-plasmid network with Mash distance^17^ (v. 2.3; default parameters except sketch size -s 1000000) weighted edges, at a threshold *d*<0.0001. Briefly, Mash distance estimates an evolutionary distance on a reduced-length MinHash sketch of the sequences. Since Mash is a probabilistic estimate of evolutionary distance, we confirmed the probability of seeing any of our pairwise Mash distances in the near-identical groups by chance was 0. For whole genomes, Mash distance has a strong positive correlation with ANI ^18^. We also required the shortest plasmid to be within 1% length (bp) of the longest plasmid, to account for assembly errors. Network analysis was performed using the igraph^19^ library (v. 1.2.7) in R.

**Accumulation and rarefaction curves**

To generate an accumulation curve, isolates were sampled without replacement in a random order. For each isolate, the new plasmid diversity was recorded. For Fig. S4, we recorded the number of new near-identical plasmid groups and singletons. For Fig. S9, we recorded the number of near-identical matches with BSI plasmids from only environmental/livestock isolates. For Fig. S7, we recorded the number of new clusters, doubletons, and singletons. A bootstrapped average of *b*=1,000 accumulation curves was plotted for the rarefaction curve. The bootstraps were also used to estimate Heap’s parameter (γ) by fitting a linear regression to log-log transformed data using standard R libraries. For γ<0, it is possible to sample the entire diversity, and for 1>γ>0, the diversity will increase with every additional sample^20^.

**Plasmid similarity**

Plasmid Jaccard index (*JI*) was calculated using Mash^17^ (v. X; default parameters except sketch size -s 1000000). The Jaccard index (*JI*), given by

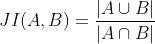

where *A*, *B* are the sets of *k*-mers of plasmids *a*, *b*, respectively. This measures extent of *k*-mer sharing between plasmids, range=0-1, where 1 indicates an identical *k*-mer repertoire. Since the sketch size was larger than the plasmid lengths (except for one plasmid in the dataset, OX-ENV-67_2, which was larger than 1Mbp at 1,310,597bp and was not clustered; the next smallest was OX-WTW-80_2 at 394,284bp), the calculated Jaccard indices were almost always exact.

**Plasmid network and clustering**

The determination of the plasmid-plasmid network, threshold, and clusters could be achieved with several alternative methodologies. Plasmid networks have previously been constructed by full sequence alignments^21^, annotated genes^22^, and alignment-free Mash distances^9,23,24^. We chose to use the Jaccard index of entire plasmid *21*-mer distributions to capture coding sequences, their immediate contexts^25,26^, and intergenic regions^27,28^, all of which have known importance to bacterial evolution. Further, our contained previously unsampled diversity as seen by the PTU analysis, and because reference-based classifications such as MOB and replicon typing schemes are known to be incongruent^29^ or unreliable: 16% (602/3,697) of our plasmids had an unidentifiable replicon type, which is not uncommon^8^. The evolutionary histories of plasmids can incorporate multiple gain, loss, and rearrangement events in addition to mutations^30^, and as such, traditional measures of genetic relatedness (e.g., single nucleotide variant [SNV] thresholds) used for genomic epidemiology of whole genomes are likely less appropriate here. These similarities formed the edge-weights in a plasmid-plasmid network, which was subsequently thresholded to sparsify the network and allow the detection of clusters.

Network thresholding to some extent depends subjectively on the dataset, with trade-offs between successfully revealing the underlying structure of plasmid relationships without excessively separating relatives. We chose a data-driven threshold as adopted by Ledda *et al.*, 2018^22^ for their plasmid network, which examined the evolution of connected components within the network. This ensured the threshold was chosen where the regime of connected component evolution approximately stabilises, minimising excessive network breakup. The threshold was chosen at *JI*=0.5, meaning that edges between plasmids with *JI*<0.5 were deleted from the network. From this threshold onwards, both the number of connected components and the number of singletons steadily increased at a similar rate (Fig. S5). This regime indicates an approximately stable non-singleton structure from *JI*=0.5 onwards.

We defined plasmid clusters as groups of *n*≥3 plasmids with high within-cluster-similarities and low between-cluster-similarities. Plasmid clusters were detected using the Louvain algorithm which optimises the network modularity by iterative expectation-maximisation^31^. This aims to maximise the density of edges within clusters against edges between clusters. Though non-deterministic, the Louvain algorithm showed low variation in cluster distribution over 50 runs, consistent with reproducible segregation of plasmids in clusters (range of clusters detected: 245-247; Fig. S6). The algorithm was implemented using the python-louvain (v. 0.16) Python module. Although the algorithm is non-deterministic, multiple runs demonstrated minimal variation at our chosen network threshold. Overall, these approaches add to the growing literature describing suitable methodologies for clustering plasmids.

Near-identical plasmid groups were also included in the wider cluster analysis, as many were cross-compartmental and found across bacterial hosts (see earlier, Fig. 2). Of the n=194/225 groups which were clustered, 100% (194/194) had all members fall within the same plasmid cluster, with *n*=30/247 clusters containing multiple near-identical plasmid groups. Only 6% (14/247) of plasmid clusters comprised exclusively near-identical plasmid groups, suggesting that near-identical groups of plasmids often have nearby genetically related plasmids. Examining the entire PTU distribution within clusters, most contained at least one unclassified plasmid (51%; 127/247) or plasmid assigned a putative PTU (9%; 23/247). However, many clusters exclusively contained just one known PTU (42%; 105/247).

**Cluster homogeneity and completeness**

Homogeneity (*h*) and completeness (*c*) are dual conditional entropy-based measures, independent of cluster and metadata label distributions^32^. A clustering satisfies homogeneity (*h* =1) if all cluster members have the same metadata label-type. Consider a network with $N$ nodes, partitioned by a set of metadata labels, $M=\{m_{i}|i=1,\ldots,n\}$, and a set of communities, $C=\{c_{j}|j=1,\ldots,m\}$. Let $A=\{a_{ij}\}$ represent the $ij^{\text{th}}$ entry in the contingency table of partitions. Hence, $a_{ij}$ counts the number of nodes with label $m_{i}$ in community $c_{j}$. We then say

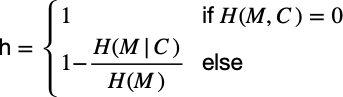

where

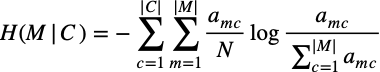

and

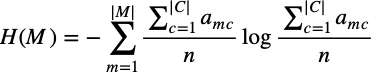

are the conditional entropy of the metadata given the clusters and the entropy of the clusters, respectively $H(M|C)=0$ when the cluster partition coincides with the metadata partition, and no new information is added. A cluster partition satisfies completeness (*c* =1) if all instances of a metadata label-type are assigned the same cluster. Completeness is defined dually by

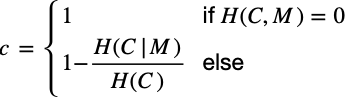

The measures were calculated using the clver library (v. 0.1.1) in R.

**Cluster pangenome analysis**

Cluster pangenomes were generated using Panaroo^33^ (v. 1.2.9) with parameters default except --clean-mode sensitive, --aligner mafft, -a core, and --core_threshold 0.95. For core gene alignments, the threshold was set at minimum 95% presence amongst clustered plasmids, whereby they were aligned using MAFFT^34^ (v. 7.407) with default parameters. An identical approach was taken for the host chromosome phylogeny in Figure 4. The median length of plasmids within a cluster was positively correlated with number of core genes (R=0.85, t=13.4, p-value<2.2e-16) and total pangenome size (R=0.87, t=14.6, p-value<2.2e-16).

**Plasmid core gene phylogenies**

Maximum likelihood core-gene phylogenies were generated using IQ-Tree^35^ (v. 2.0.6) with parameters -m GTR+F+I+G4 -keep-ident -T 2 -B 1000. The substitution model used was general time reversible (GTR) using empirical base frequencies form the alignment (F), allowing for invariable sites (I) and variable rates of substitution (G4). We used *n*=1000 ultrafast bootstraps (B 1000; see Minh *et al.*, 2013^36^) to visually inspect larger clades for support. Briefly, 95% support approximates a 95% probability that the clade is genuine. Only the *n*=62/69 clusters (excluding 6, 8, 26,2 9,3 2, 40, and 65) where every plasmid carried at least 1 core gene were analysed. Phylogenies were primarily plotted using the R library ggtree^37^.

**Fritz and Purvis’ *D***

Fritz and Purvis’ *D* measures phylogenetic signal for binary traits^38^. First, we calculate the character state changes required to observe our phylogeny (*d_obs_*). To account for phylogeny size and prevalence, *d_obs_* is standardised under the two null models (i) tip labels are random permuted (*d_r_*), and (ii) tip labels are distributed under the expectation of a Brownian motion model of evolution (*d_b_*). Then, we define

*D* = (*d_obs_* – $\bar{d_{b}}$)/( $\bar{d_{r}}$– $\bar{d_{b}}$).

Hence, for *D*$\approx$1, *d_obs_* follows *d_r_* more closely, and for *D*$\approx$0, *d_obs_* follows *d_b_* more closely. We calculated *d_obs_* *n*=10,000 times and averaged the result, as well as calculate *p*-values for significant deviation from *d_r_* or *d_b_*. *D* was implemented using the R library caper^39^. Fritz and Purvis’ *D* is normally used for cross-species analysis so is not benchmarked for plasmids. Results for phylogenies with less than 25 tips should be viewed more conservatively due to reduced statistical power in these instances.

We considered the binary ‘trait’ of human or livestock-associated isolate and estimated *D* with *n*=10,000 permutations. We found 42% (11/26) clusters had *D*>0.5 (see Table S1). However, only 23% (6/26) of phylogenies were significantly different (*p*-value<0.05) from the conserved null model, compared to 50% (13/26) significantly different from the random null model.

**Consensus gene synteny heatmaps**

For each cluster, we first generated a list of every possible pair of genes in the pangenome. Then for each plasmid, we counted the distance between these pairs, modulo the number of genes in the plasmid. If a gene was absent in a plasmid, NA was used. We then calculated the median of these values across all plasmids in the cluster. We then built a dendrogram from a hierarchal clustering of the median distances. The order of the tip labels in the dendrogram were then used as the ‘consensus gene synteny’.

**Accessory gene distances**

Plasmid accessory gene distances were calculated using pairwise Jaccard distances on gene presence-absences matrices. For plotting the cluster-wise plasmid core gene cophenetic distance against accessory gene presence-absence Jaccard distance, only the *n*=26/62 clusters with at least 50 accessory genes were plotted. The log-transformed linear regression of Jaccard distance of accessory genes presence against core gene cophenetic distance was fitted in R with standard libraries.

**Chromosome core gene phylogeny**

An identical approach was taken to the plasmid phylogenies. *E. coli* phylogroups were typed using EzClermont^40^ (v. 0.7.0) with default parameters. Robinson-Foulds distance was calculated using the R library phangorn^41^.

**Plasmid mutation rate**

Mutation rates per base pair in microbes typically arise from DNA replication and tend to be below *m*=10^-9^ per site per generation^42^ or perhaps as low as 10^-10^ per site per generation^43,44^. For a plasmid of size *L*, one therefore expects *L* x *m* mutations per plasmid per generation. For example, if the plasmid has *L*=10^5^ then in each generation 1 in 10,000 plasmids will gain a mutation. The generation time of *E. coli* per day in the human gut has been estimated to be between 6 and 20 generations per day^45^. For large plasmids that exist at a copy number of ~1, the plasmid generation time is the cell generation time. More generally, for a plasmid copy number *p* the number of replications of the plasmid expected for a given number of cell generations *g* will be *p* x *g* (assuming that plasmid copies are simply and linearly related to the realised number of replications per cell). A crude estimate for the expected mutation rate per time period for a plasmid is therefore given by *L* x *m* x *p* x *g.* For a plasmid of *L*=100 kbp and *p*=1, assuming *m*=[0.1-1]x10^-9^ per site per generation and *g*=[6-20]x365 per year, one would expect it to accumulate ~0.5 mutations a year (between ~0.02-0.7 depending on assumptions). One obtains the same result for *L*=10 kbp and *p*=10. There is a strong inverse correlation between plasmid size and copy number. This suggests that a suitable upper bound for the expected number of mutations for a typical plasmid per year (under neutral evolution) is of the order of magnitude of 1 SNP a year. This rough ‘SNPs and years’ rule-of-thumb appears consistent with known empirical results. For example: 100kbp I1-type *Shigella* plasmids isolated between 2007-2010 in Vietnam were separated by at most 2 SNPs^46^; 30kbp X4-type plasmids carrying *mcr-1* isolated between 2016-2018 in China were separated by most 4 SNPs^47^ (analysis not shown); 63.5 kbp pOXA48-like plasmids (*n*=202) in *Klebsiella* *pneumoniae* collected across Europe between 2013 and 2014 as part of EUSCAPE were overwhelmingly within 2 SNPs of each other (176/202)^48^; the same was true of 45.4kbp IncX3 plasmids (*n*=135) from the EUSCAPE dataset (all were within 6 SNPs of each other; see Figure 4 of that paper); and also of 113.4 kbp pKpQIL-like plasmids (*n*=91) from the EUSCAPE dataset – although a minority of these plasmids were separated by up to 20 SNPs, which seems suggestive of either ancestry before the two-year sampling frame or recombination.

**Pangraph analysis**

We used pangraph^49^ (v. 0.5.0) to build a pangraph of the clade within plasmid cluster 2, using the --circular flag and otherwise default parameters. We removed duplicated blocks from the pangraph. We used pangraph export (--edge-minimum length 0, default parameters) to export the graph to GFA format and then visualised this using Bandage^50^.

**Data visualisation**

Plots were primally produced using the R library ggplot2^51^, with additional graphics in BioRender^52^.

1.	Torsten Seeman. mlst. Preprint at (2017).

11.	Torsten Seeman. Abricate. Preprint at (2015).

19.	Csardi, G. & Nepusz, T. The igraph software package for complex network research. *InterJournal Complex Systems* **Complex Sy**, (2006).

32.	Rosenberg, A. & Hirschberg, J. V-Measure: A conditional entropy-based external cluster evaluation measure. in *EMNLP-CoNLL 2007 - Proceedings of the 2007 Joint Conference on Empirical Methods in Natural Language Processing and Computational Natural Language Learning* (2007).

39.	Orme, D. The caper package : comparative analysis of phylogenetics and evolution in R. *R package version 0.5, 2* (2013).

49.	Nicholas Noll, M. M. R. A. N. PanGraph: scalable bacterial pan-genome graph construction. *bioRxiv* (2022).

50.	Wick, R. R., Schultz, M. B., Zobel, J. & Holt, K. E. Bandage: Interactive visualization of de novo genome assemblies. *Bioinformatics* **31**, (2015).

51.	Gómez-Rubio, V. ggplot2 - Elegant Graphics for Data Analysis (2nd Edition) . *Journal of Statistical Software* **77**, (2017).

52.	Munday, E. BioRender. *N/a* (2021).
