## Supplementary Figures for "*Enterobacterales* plasmid sharing amongst human bloodstream infections, livestock, wastewater, and waterway niches in Oxfordshire, UK"

**Fig. S1.** Mash tree for *n*=1,044 *E. coli* chromosomes. Tree tips are coloured by sampling compartment, scale is Mash distance.

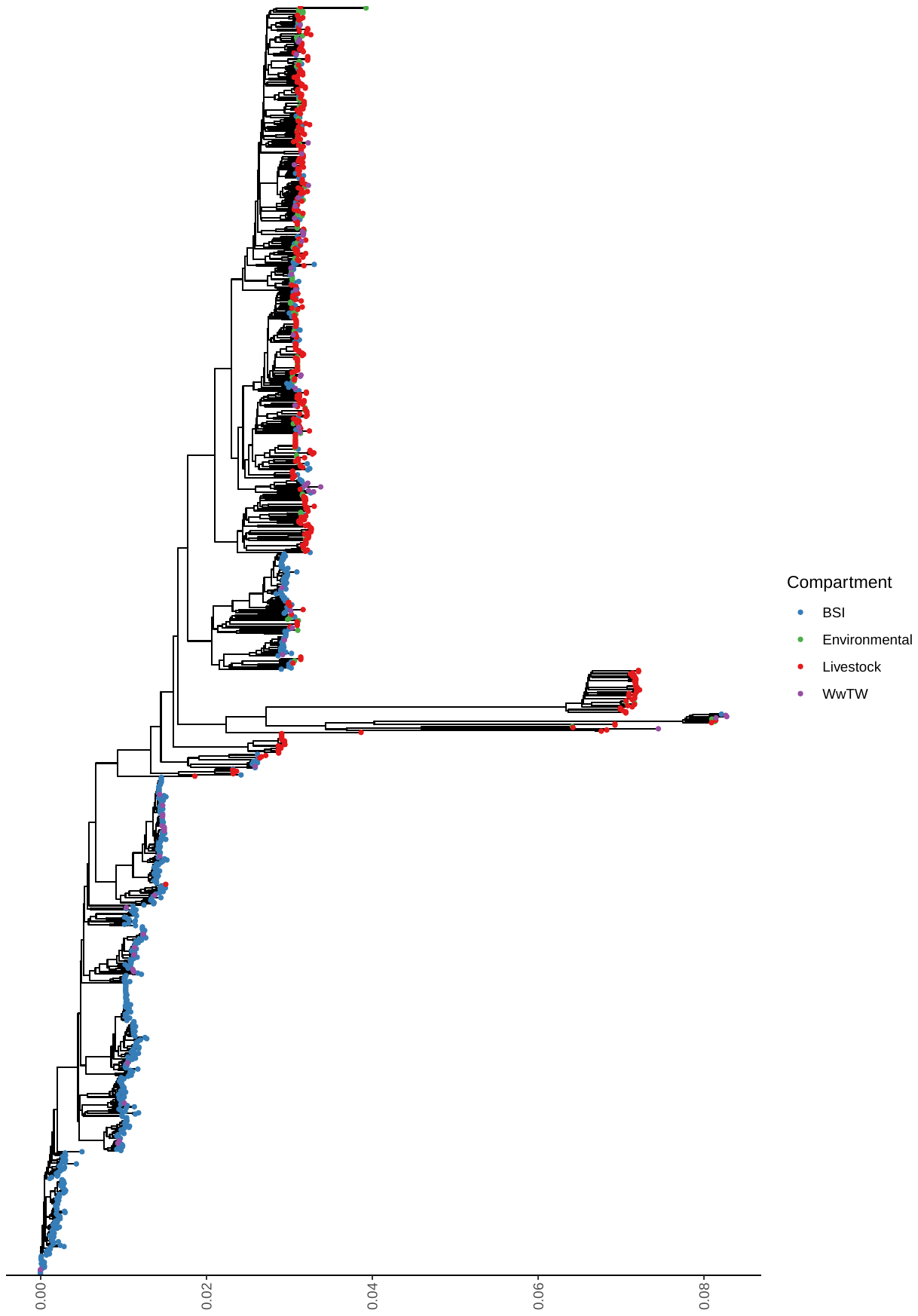

**Fig. S2.** Mash tree for *n*=163 *K. pneumoniae* chromosomes. Tree tips are coloured by sampling compartment, scale is Mash distance.

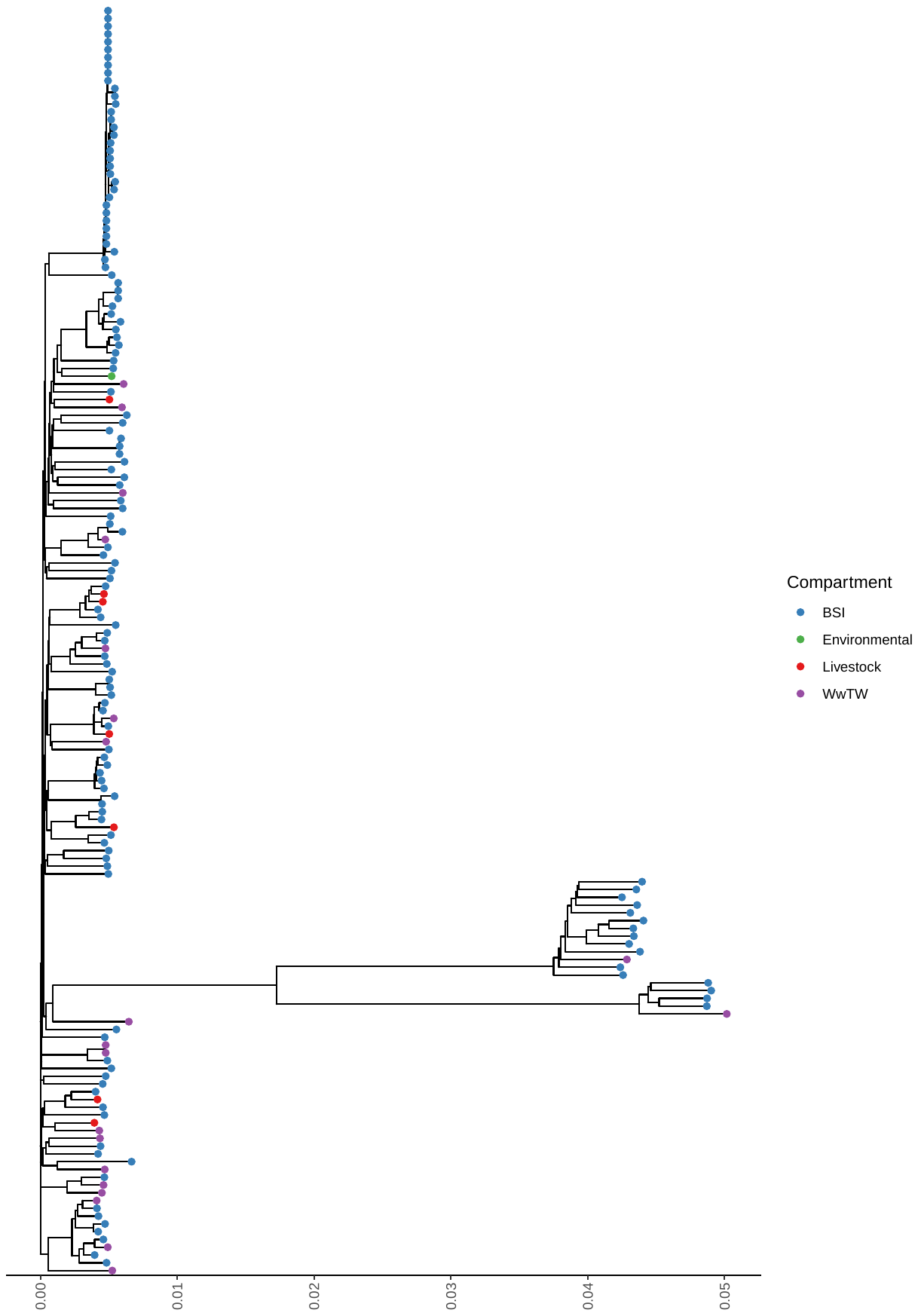

**Fig. S3.** Accumulation curves of near-identical plasmid groups and singletons against isolate sample size. Black lines represent *b*=1000 bootstrap simulations, the red line represents their average.

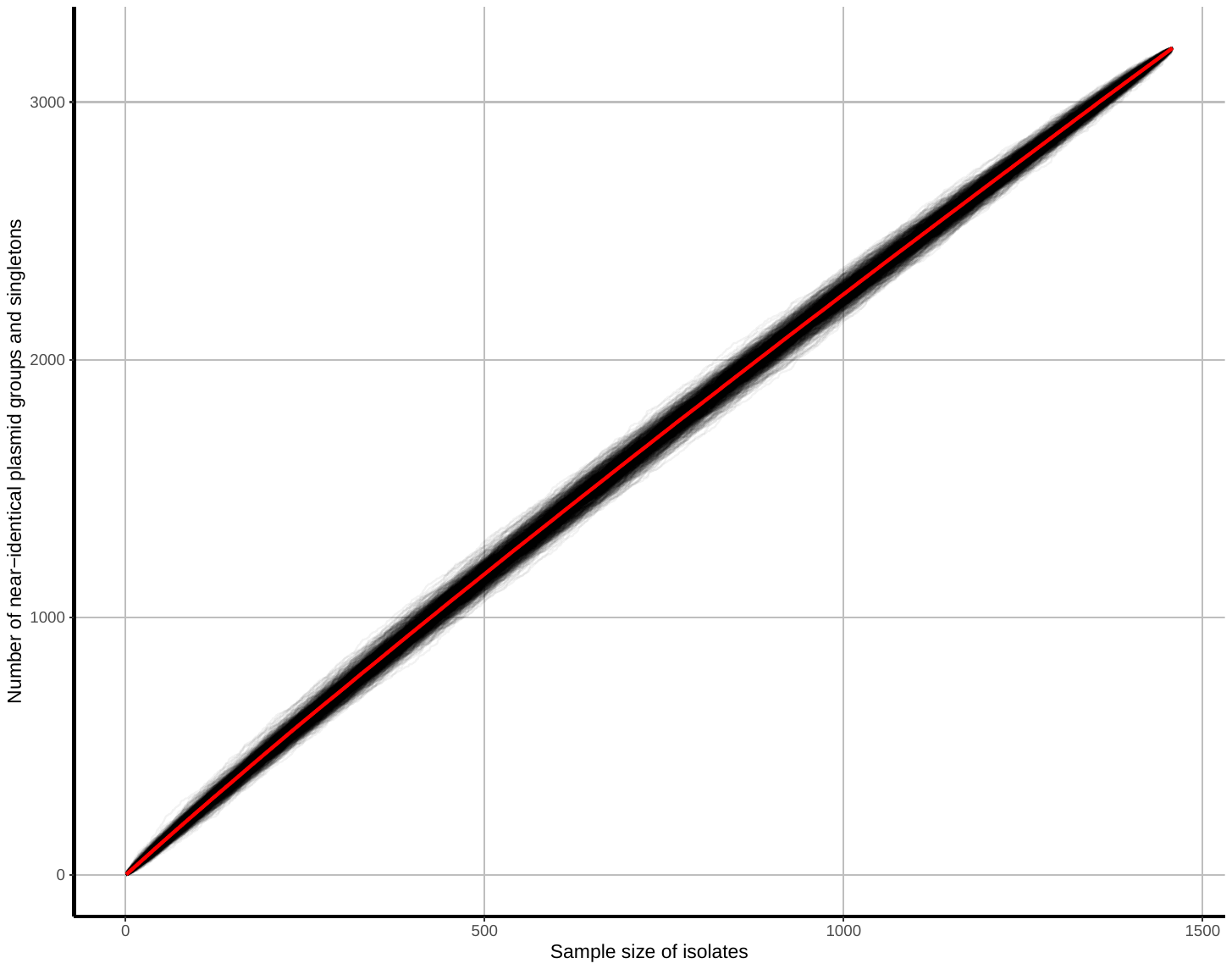

**Fig. S4. Plasmid clusters containing *bla*_TEM-1_ carry more AMR genes.** Each point is one plasmid cluster. *n*=247 clusters are shown, with panels facetted by the number of niches the plasmid cluster represented. *p*-values are from the Wilcoxon test.

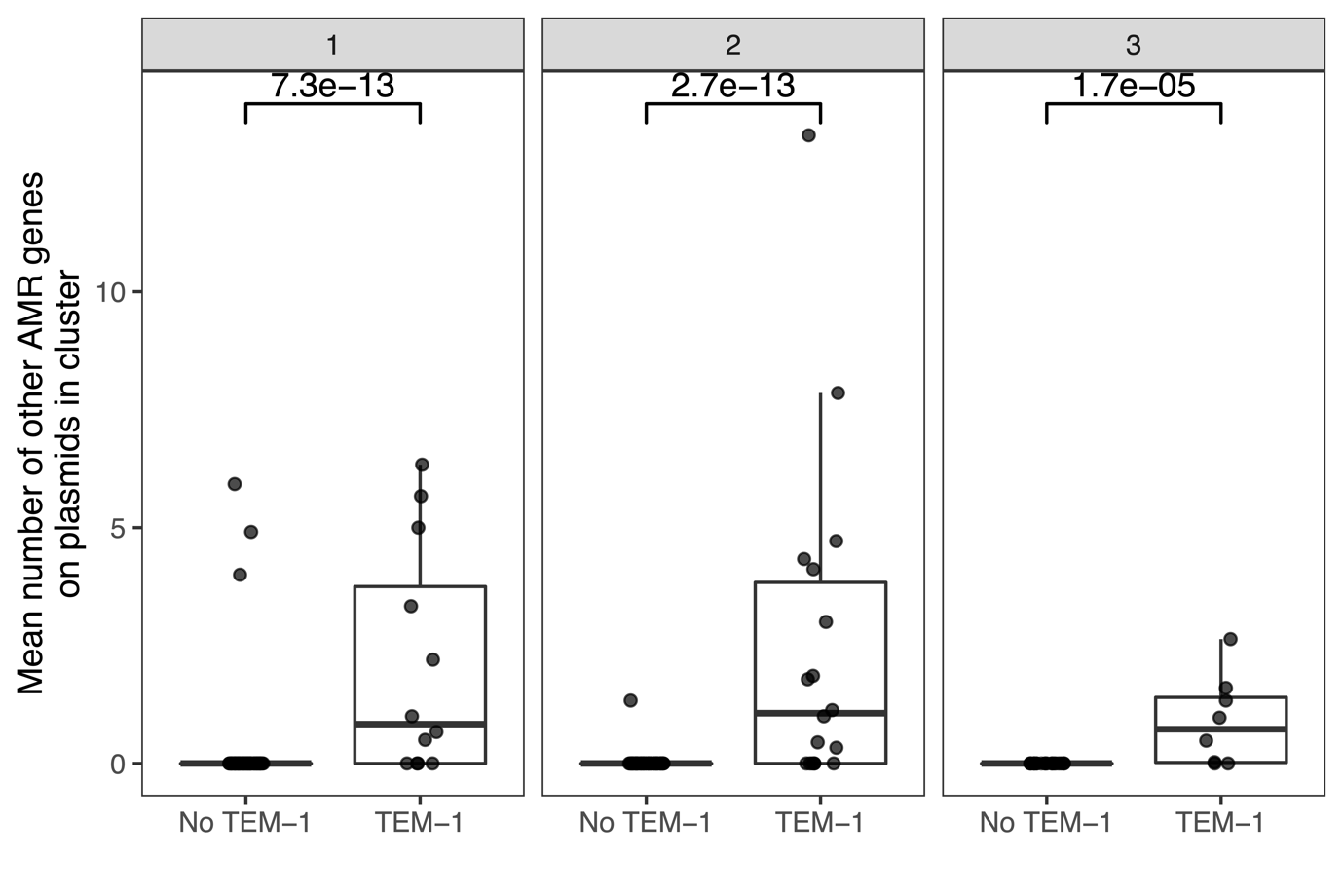

**Fig. S5.** Network evolution of largest connected component, number of connected components, and number of singletons, as edges are removed at increasing JI thresholds. The vertical red line represents the chosen threshold of *JI*=0.5.

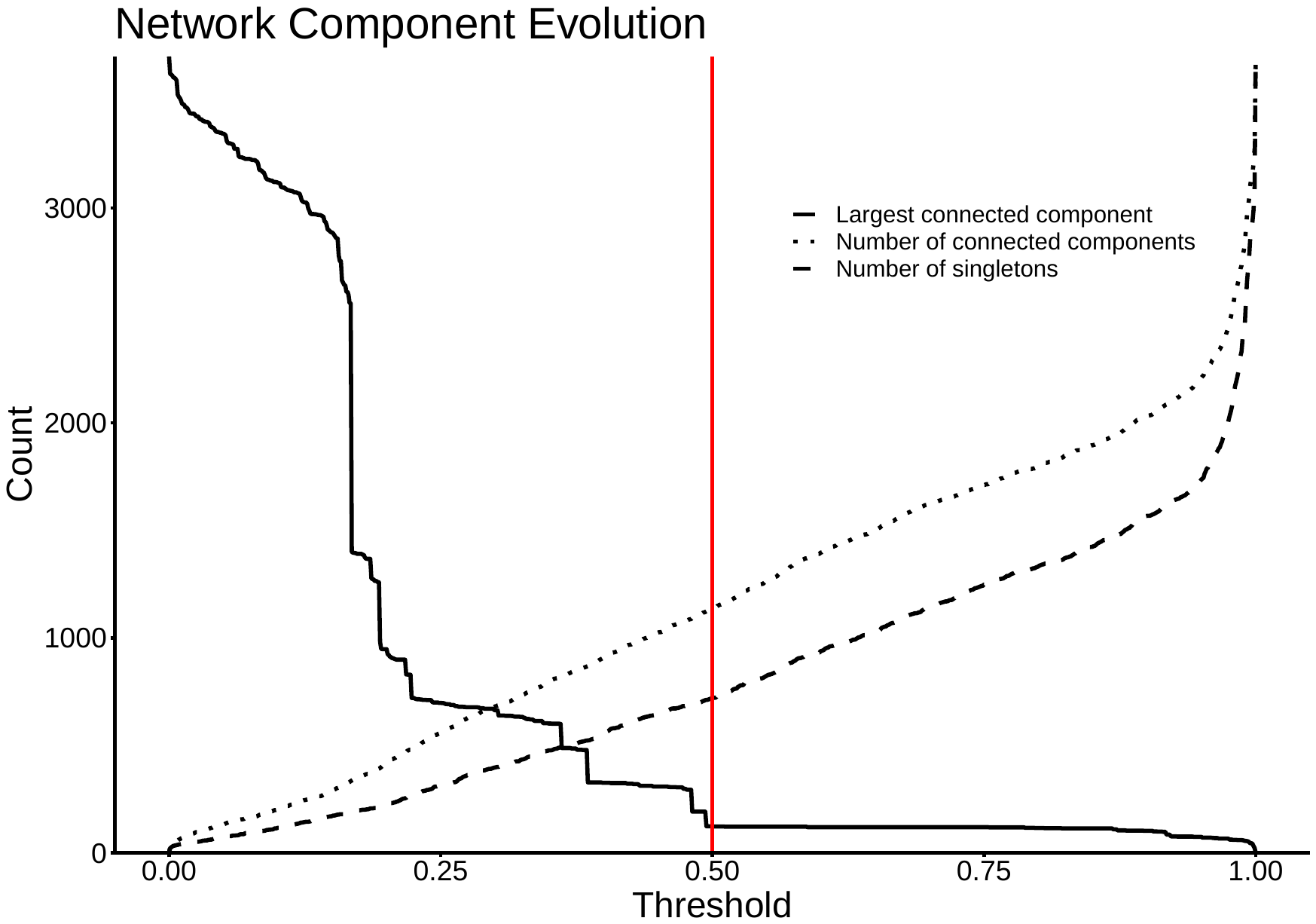

**Fig. S6.** Number of clusters detected within the plasmid network at increasing *JI* thresholds. Interval bars represent the IQR in cluster number at a given threshold over 50 runs of the Louvain algorithm.

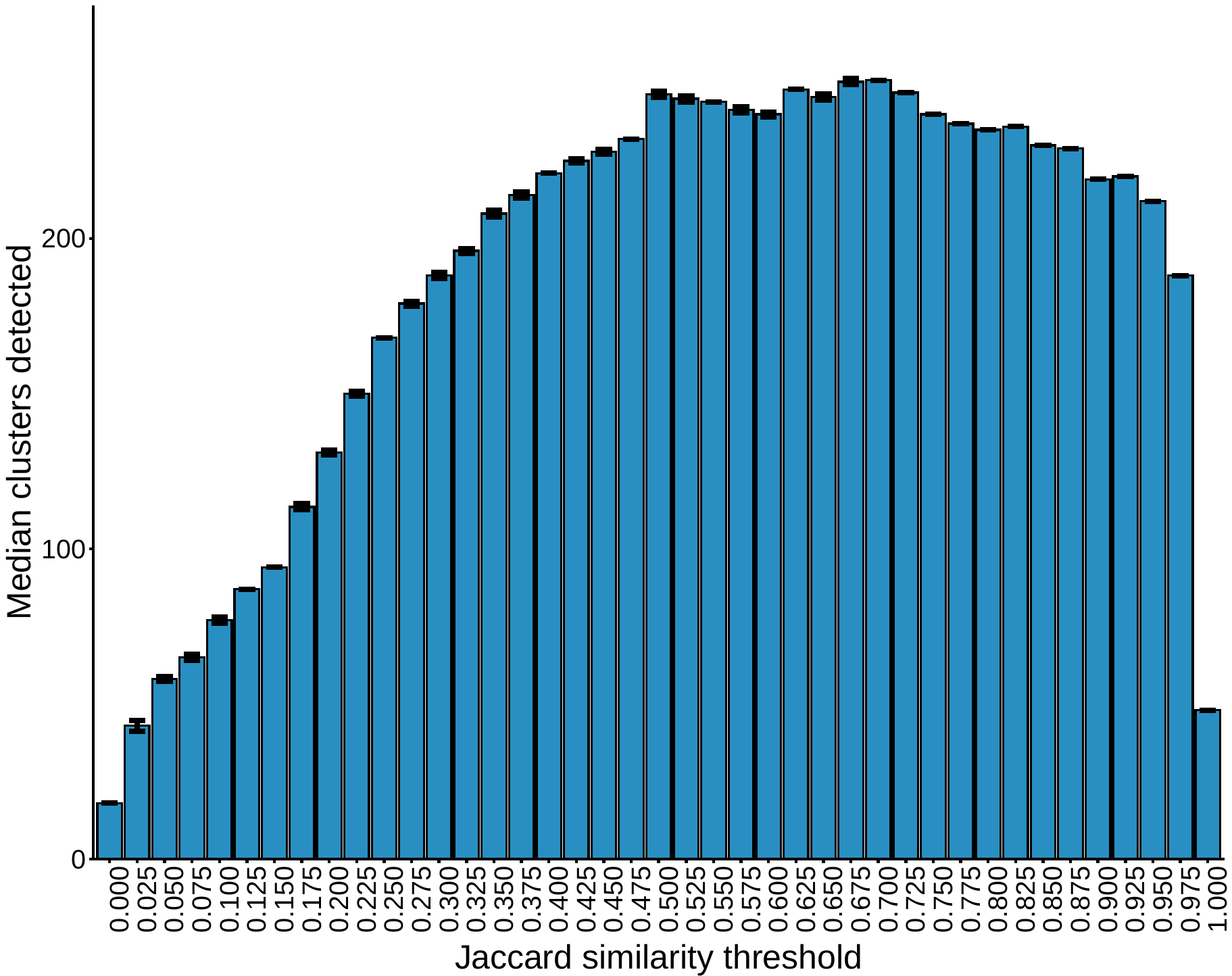

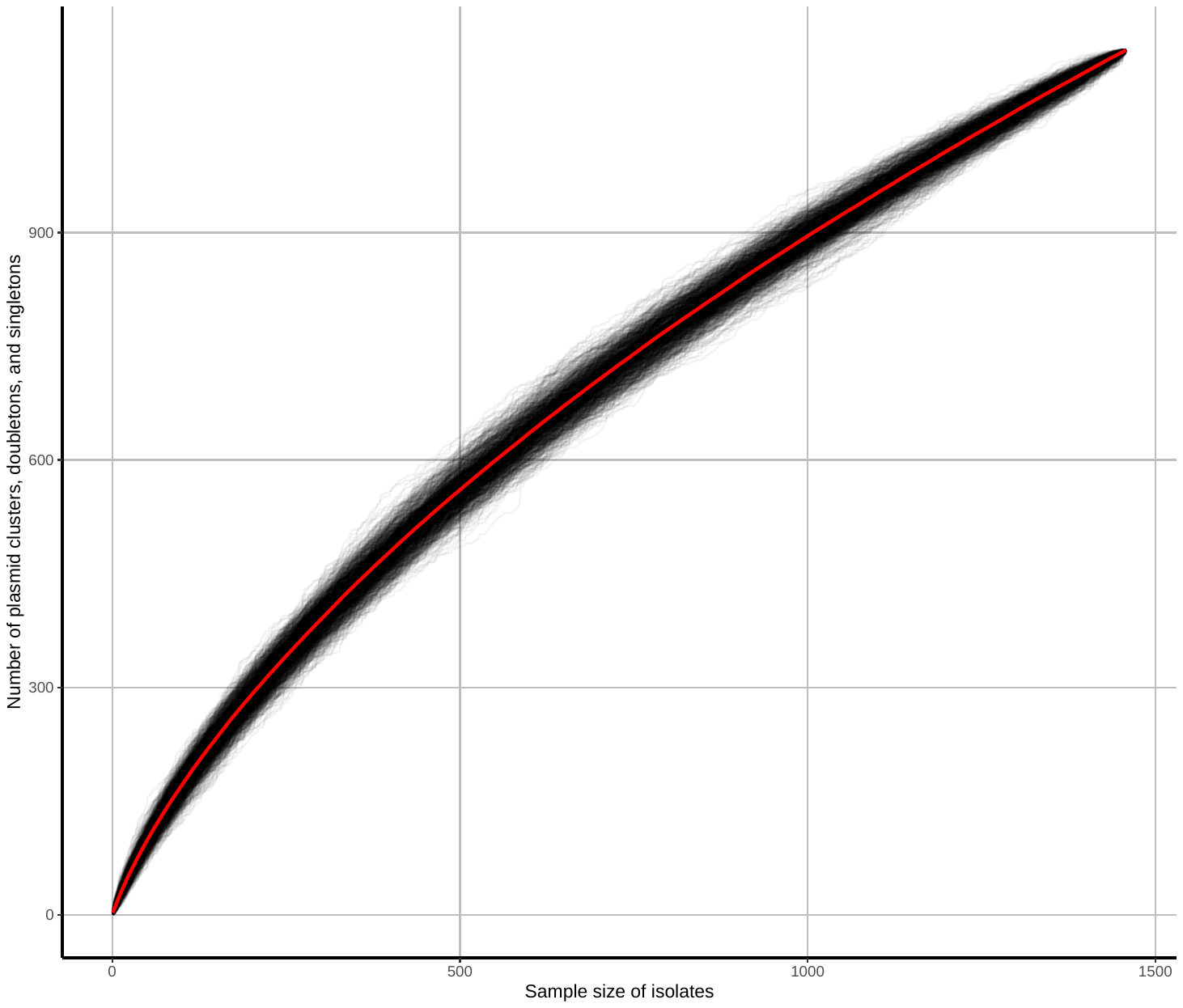
**Fig. S7.** Accumulation curves of plasmid clusters, doubletons, and singletons against isolate sample size. Black lines represent *b*=1000 bootstrap simulations, red line represents their average.

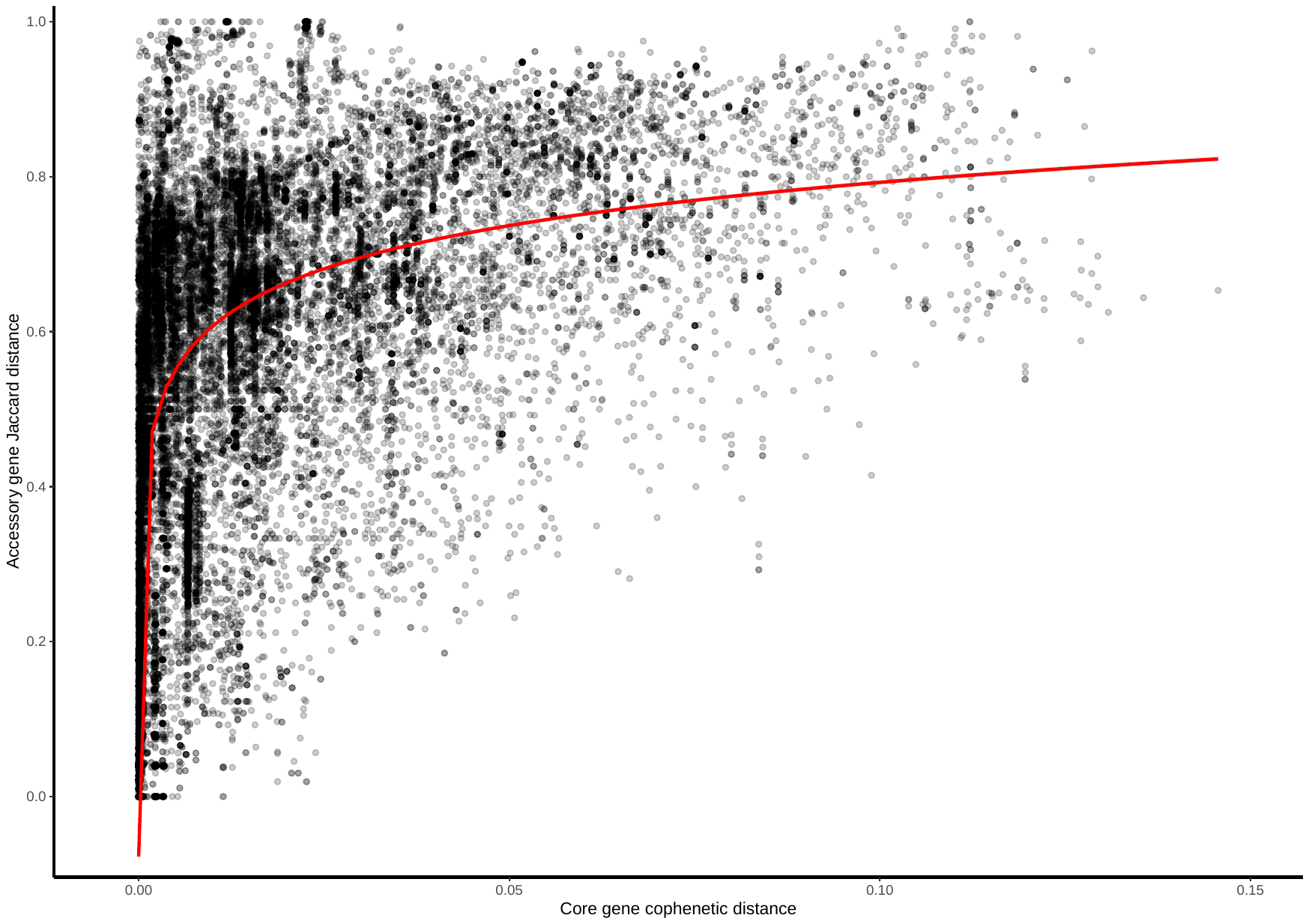
**Fig. S8.** Plasmid accessory gene presence/absence Jaccard distance against core gene cophenetic distance. Presented are data points from 27/247 clusters for which (i) all plasmids had at least 1 core gene, and (ii) the cluster contained at least 50 accessory genes. The red line is a statistically significant (p-value<2.2e-16) log-transformed linear regression.

**Fig. S9.** Accumulation curves of near-identical plasmid matches with BSI plasmids and singletons against livestock-associated (environmental soils/livestock) isolate sample size. Black lines represent *b*=1000 bootstrap simulations, red line represents their average.

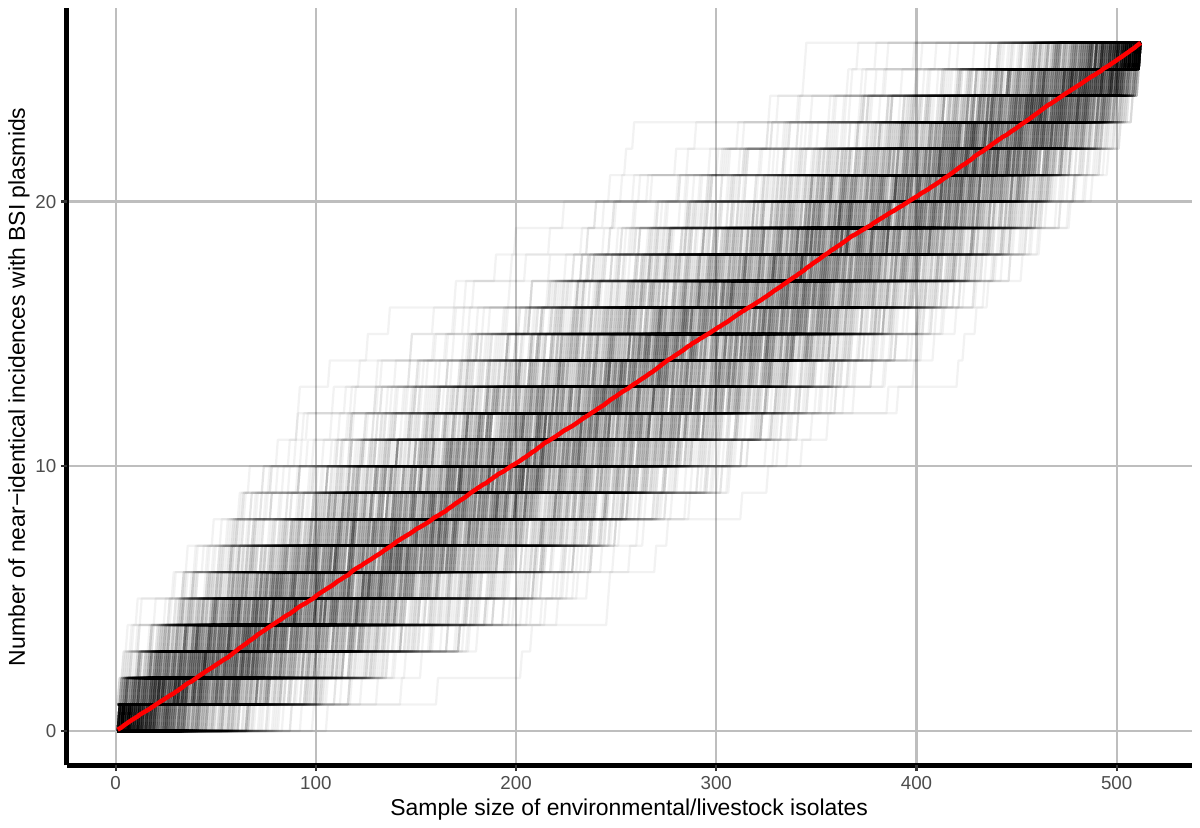
