## Supplementary File 1 for "*Enterobacterales* plasmid sharing amongst human bloodstream infections, livestock, wastewater, and waterway niches in Oxfordshire, UK"

In order, this document contains

(i) A metadata table for all plasmid clusters where every plasmid has at least one core gene. Presented are the cluster name, cluster size, mlst PubMLST genera of plasmid hosts, plasmid PlasmidFinder annotations, and plasmid NCBIAMRFinder annotations.

(ii) A core gene phylogeny and consensus gene synteny heatmap for all clusters in the metadata table, in cluster size decreasing order. Core gene phylogeny scales are in single nucleotide polymorphisms (SNPs).

| Cluster name | Cluster size | Host-PubMLST(s) | PlasmidFinder | AMRFinderPlus |
|---|---|---|---|---|
| 1 | 123 | ecoli,cfreundii | Col156_1,IncFIB(AP001918)_1,IncFII(29)_1_pUTI89,IncFII_1,IncFIC(FII)_1 | aac(3)-IId,aadA5,aph(3'')-Ib,aph(6)-Id,blaTEM-1,dfrA17,mph(A),qacEdelta1,sul1,sul2,tet(A),catA1,tet(B),blaTEM,blaCTX-M-27,blaTEM-30,dfrA8 |
| 2 | 100 | ecoli,senterica | IncFIA_1,IncFIB(AP001918)_1,IncFIC(FII)_1,IncFII_1 | aph(3'')-Ib,aph(6)-Id,blaTEM-1,dfrA14,sul2,tet(A),dfrA5,aac(3)-IId,catA1,tet(B),aph(3')-Ia,aadA1,aadA2,cmlA1,dfrA12,qacL,sul3,qacEdelta1,sul1,blaTEM-40,blaTEM,aadA5,dfrA17,catA2,floR |
| 3 | 73 | ecoli,senterica,cfreundii | IncI1_1_Alpha,IncI_Gamma_1 | blaTEM-1,qnrS1,blaCTX-M-15,blaCTX-M-1,sul2,tet(A),mcr-5.1,aph(3')-Ia,aph(3'')-Ib,aph(6)-Id,tet(B),aadA1,dfrA1,qacEdelta1,sul1,aadA2,cmlA1,qacL,sul3,blaCMY-2,blaCTX-M-14,blaLAP-2,dfrA14,sat2,blaCTX-M-55,blaTEM |
| 4 | 69 | ecoli,cfreundii,kpneumoniae | Col156_1 | blaTEM-1,blaCTX-M-27 |
| 5 | 68 | ecoli,kpneumoniae,senterica | IncFIA_1,IncFIB(AP001918)_1,IncFII(pRSB107)_1_pRSB107,Col156_1,IncFIB(pB171)_1_pB171,IncFII(pAMA1167-NDM-5)_1_pAMA1167-NDM-5,IncFIC(FII)_1,ColRNAI_1 | aph(3'')-Ib,aph(6)-Id,blaTEM-1,sul2,dfrA14,mph(A),tet(B),blaCTX-M-27,catA1,aph(3')-Ia,aadA1,dfrA1,qacEdelta1,sul1,tet(A),aadA5,dfrA17,aadA2,blaCARB-2,aac(3)-IIe,aac(6')-Ib-cr5,blaCTX-M-15,blaOXA-1,catB3,aac(3)-IId,erm(B),dfrA12 |
| 7 | 58 | ecoli | IncB/O/K/Z_2,IncB/O/K/Z_3,IncB/O/K/Z_1 | aph(3'')-Ib,blaTEM-1,sul2,aph(6)-Id,aadA1,dfrA1,qacEdelta1,sul1,tet(A),dfrA14,blaTEM-30,tet(B),blaCTX-M-14,erm(B),qacE,aadA5,dfrA17,mph(A),blaTEM |
| 9 | 51 | ecoli,senterica,kpneumoniae | Col8282_1 | |
| 10 | 47 | ecoli | IncFIB(AP001918)_1,IncFIA_1,IncFIC(FII)_1 | |
| 11 | 43 | ecoli,cfreundii,senterica | ColRNAI_1 | |
| 12 | 42 | ecoli | IncFIC(FII)_1,IncFII_1,NA,IncFII(pHN7A8)_1_pHN7A8,IncFII(pRSB107)_1_pRSB107,IncFII(pCoo)_1_pCoo,IncFIA_1,IncFIB(AP001918)_1 | blaTEM-1,NA,aph(3'')-Ib,aph(6)-Id,sul2 |
| 13 | 41 | kpneumoniae,koxytoca,ecoli | IncFII_1_pKP91 | aac(3)-IIe,aac(6')-Ib-cr5,aph(3'')-Ib,aph(6)-Id,blaCTX-M-15,blaOXA-1,blaTEM-1,catB3,dfrA14,sul2,qnrB1,tet(A),blaTEM |
| 14 | 40 | ecoli | IncFIB(AP001918)_1,IncFIB(pB171)_1_pB171,IncFIA_1,IncFIC(FII)_1 | |
| 15 | 38 | ecoli,cfreundii | Col156_1,IncFIB(AP001918)_1,IncFIC(FII)_1,IncFIA_1,IncFII(29)_1_pUTI89 | blaTEM-1,tet(B),aph(3'')-Ib,aph(6)-Id,sul2,mph(A),aadA1,blaSHV-1,qacEdelta1,sul1,tet(D),aadA2,dfrA12,blaSHV |
| 16 | 37 | kpneumoniae,koxytoca | IncFIB(K)_1_Kpn3,IncFII_1_pKP91 | aadA1,blaSHV-1,dfrA1,qacEdelta1,sul1,tet(D),aph(3'')-Ib,aph(6)-Id,blaTEM-1,dfrA14,mph(A),sul2,tet(A),aph(3')-Ia,aac(3)-IIe,blaSCO-1,catA1,dfrA8,aac(6')-Ib-cr5,blaCTX-M-15,blaOXA-1,catB3,qnrB1,blaSHV,dfrA7,aac(3)-IId,aadA2,blaAAK,blaSHV-12,blaSHV-2A |
| 17 | 35 | ecoli | ColRNAI_1 | blaTEM |
| 18 | 34 | kpneumoniae | | blaCTX-M-15 |
| 19 | 33 | kpneumoniae | IncFIB(K)_1_Kpn3 | |
| 20 | 32 | kpneumoniae | IncFIA(HI1)_1_HI1 | |
| 21 | 29 | ecoli,kpneumoniae | Col8282_1,Col(MG828)_1 | |
| 22 | 29 | ecoli | Col(BS512)_1 | |
| 23 | 26 | ecoli | IncX1_1 | blaTEM-1,sul2,tet(A) |
| 24 | 25 | ecoli | Col156_1 | |
| 25 | 24 | ecoli,cfreundii | RepA_1_pKPC-CAV1321 | aac(3)-IId,aadA2,aph(3'')-Ib,aph(6)-Id,blaTEM-1,dfrA12,mph(A),qacEdelta1,sul1,sul2,tet(B) |
| 27 | 23 | ecoli | IncFIB(pHCM2)_1_pHCM2 | blaCTX-M-15 |
| 28 | 21 | ecloacae,koxytoca | ColRNAI_1 | |
| 30 | 21 | ecoli | Col156_1,IncFIA_1,IncFIB(AP001918)_1,IncFIC(FII)_1 | aac(3)-IIe,aac(6')-Ib-cr5,aadA5,blaCTX-M-15,blaOXA-1,catB3,dfrA17,mph(A),qacEdelta1,sul1,blaTEM-1,tet(B),tet(A),aac(3)-IId,catA1,blaDHA-1,qacE,qnrB4 |
| 31 | 20 | ecoli,cfreundii | Col156_1 | |
| 33 | 19 | ecoli,koxytoca | IncHI2_1,IncHI2A_1,RepA_1_pKPC-CAV1321 | aadA1,aph(3'')-Ib,aph(6)-Id,blaTEM-1,dfrA1,qacEdelta1,sul1,sul2,tet(A),aac(3)-IVa,aadA2,aph(3')-Ia,aph(3')-IIa,aph(4)-Ia,ble,cmlA1,dfrA12,qacL,sul3,oqxB,tet(B),NA,mef(B),floR,mph(B) |
| 34 | 18 | koxytoca,ecloacae,ecoli,cfreundii | Col440II_1,ColRNAI_1 | |
| 35 | 17 | ecoli | IncI2_1_Delta,IncI2_1 | |
| 36 | 17 | ecoli | IncFIB(AP001918)_1,IncFIC(FII)_1,IncFIA_1,IncFIB(pB171)_1_pB171 | |
| 37 | 16 | ecoli | ColRNAI_1 | |
| 38 | 15 | ecoli,cfreundii,koxytoca | ColRNAI_1 | aph(3'')-Ib,aph(6)-Id,dfrA14,sul2,blaTEM-1,tet(A) |
| 39 | 15 | ecoli | | blaTEM-1 |
| 41 | 14 | ecoli | ColRNAI_1 | |
| 42 | 14 | ecloacae | IncFIB(pECLA)_1_pECLA,IncFII(pECLA)_1_pECLA,IncR_1 | |
| 43 | 14 | ecoli,kpneumoniae | IncX4_1,IncX4_2 | aadA1,blaTEM-31,lnu(G),mcr-1,blaCTX-M-14 |
| 44 | 14 | ecoli | Col440I_1,ColRNAI_1 | |
| 45 | 14 | ecoli | | |
| 46 | 14 | ecoli | ColRNAI_1 | |
| 47 | 13 | ecoli,cfreundii | | |
| 48 | 13 | kpneumoniae,ecloacae,koxytoca | Col440I_1,Col440II_1 | |
| 49 | 13 | kpneumoniae | Col440I_1 | |
| 50 | 13 | ecoli | IncX1_1 | aadA1,aadA2,blaCTX-M-32,cmlA1,dfrA12,qacL,qnrB19,sul3 |
| 51 | 12 | koxytoca,ecloacae | Col440II_1,ColRNAI_1,Col440I_1 | |
| 52 | 11 | ecoli | ColRNAI_1 | |
| 53 | 11 | koxytoca,kpneumoniae | Col440I_1,Col440II_1 | |
| 54 | 11 | ecoli | Col156_1 | |
| 55 | 11 | ecoli | Col156_1 | |
| 56 | 11 | kpneumoniae,koxytoca | Col440I_1,IncFIB(K)_1_Kpn3,IncFII_1_pKP91,IncR_1 | |
| 57 | 11 | ecoli,koxytoca | IncFIA(HI1)_1_HI1,IncHI1A_1,IncHI1B(R27)_1_R27 | aph(3'')-Ib,aph(3')-Ia,aph(6)-Id,blaCTX-M-1,mph(A),sul2,tet(B) |
| 58 | 11 | ecoli,kpneumoniae | IncN_1,ColE10_1 | aph(3'')-Ib,aph(6)-Id,blaTEM-54,dfrA14,sul2,tet(A),aadA22,dfrA1,qacEdelta1,sul1,blaCTX-M-15,blaTEM-1,aac(6')-Ib-cr5,aadA16,arr-3,dfrA27,qnrB6 |
| 59 | 11 | ecoli | IncHI1B(CIT)_1_pNDM-CIT,p0111_1,IncFIB(K)_1_Kpn3 | blaTEM,aadA1,dfrA1,qacEdelta1,sul1,tet(A),blaTEM-1 |
| 60 | 10 | ecoli | | |
| 61 | 10 | ecoli | ColRNAI_1 | |
| 62 | 10 | ecloacae,kpneumoniae | Col440II_1,ColRNAI_1 | |
| 63 | 10 | ecoli,kpneumoniae,ecloacae | IncN_1 | |
| 64 | 10 | ecoli | IncY_1,p0111_1 | aadA5,blaCTX-M-15,dfrA17,mph(A),qacEdelta1,sul1 |
| 66 | 10 | ecoli | IncFIA(HI1)_1_HI1,IncFIB(K)_1_Kpn3 | tet(A) |
| 67 | 10 | ecoli | ColRNAI_1 | |
| 68 | 10 | ecloacae,kpneumoniae | Col440II_1,ColRNAI_1 | |
| 69 | 10 | cfreundii,ecloacae | ColRNAI_1,Col440I_1 | |

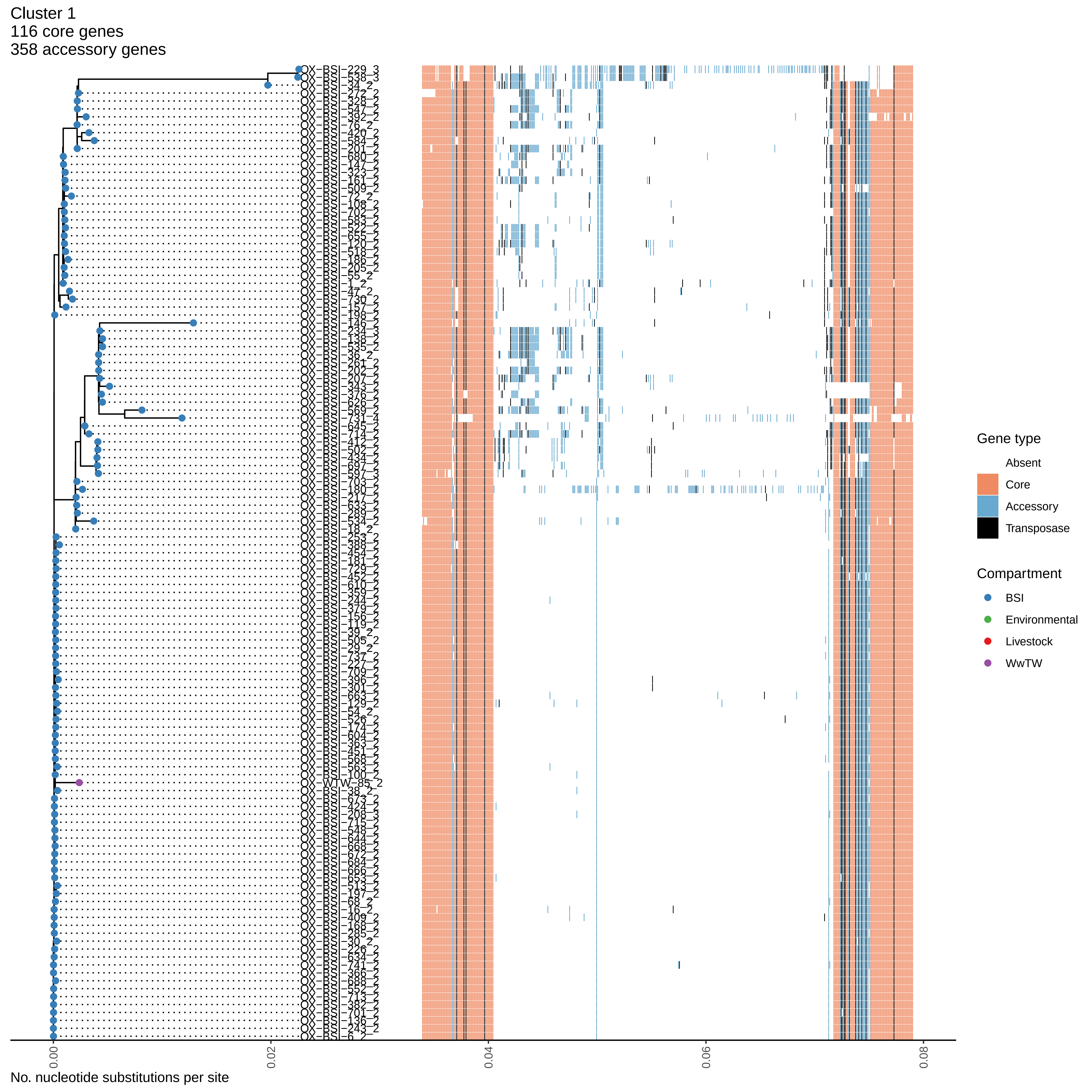
Cluster 1
116 core genes
358 accessory genes
Gene type
Absent
Core
Accessory
Transposase
Compartment
BSI
Environmental
Livestock
WwTW
0.00
0.02
0.04
0.06
0.08
No. nucleotide substitutions per site

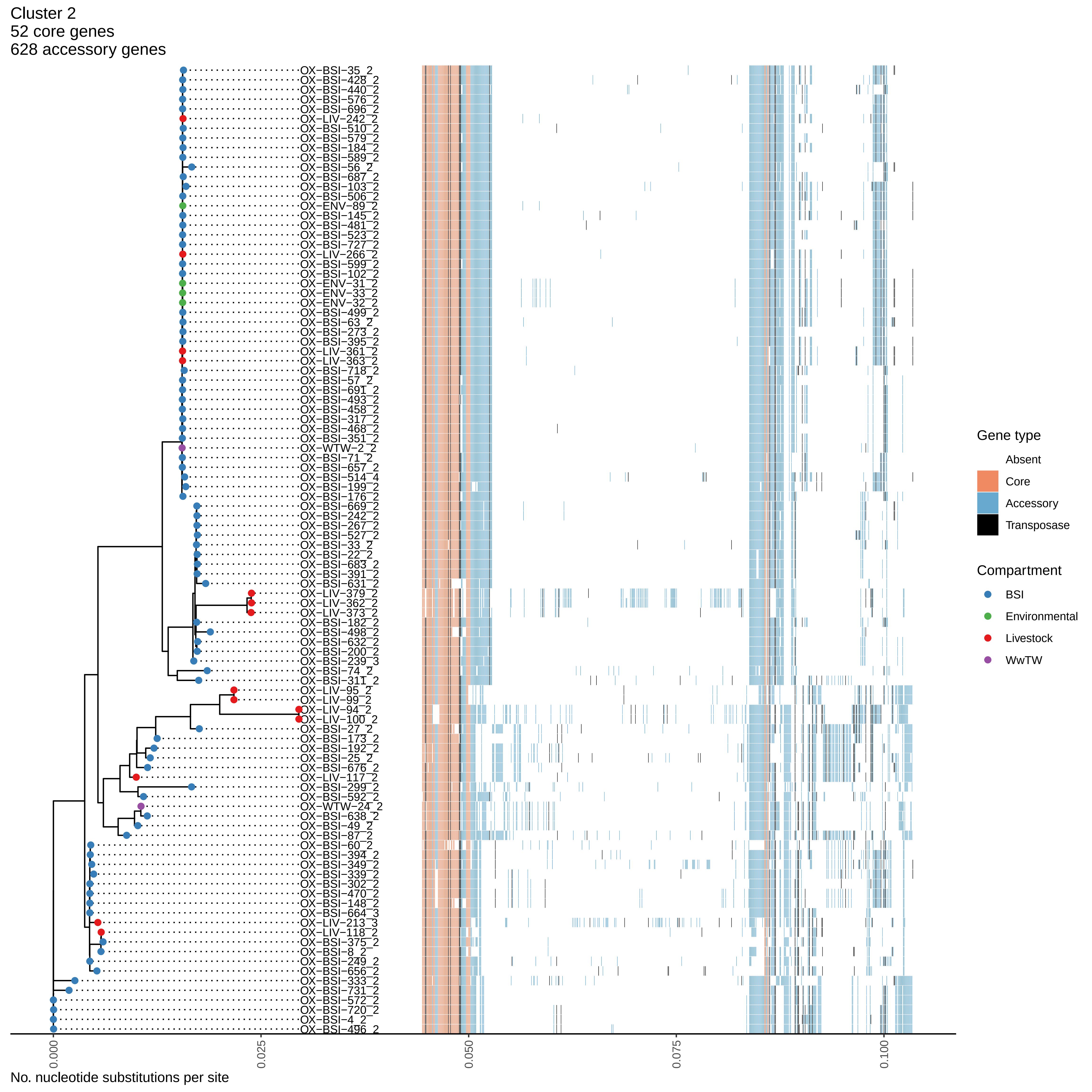

Cluster 2
52 core genes
628 accessory genes
OX-BSI-35_2
OX-BSI-428_2
OX-BSI-440_2
OX-BSI-576_2
OX-BSI-696_2
OX-LIV-242_2
OX-BSI-510_2
OX-BSI-579_2
OX-BSI-184_2
OX-BSI-589_2
OX-BSI-56_2
OX-BSI-687_2
OX-BSI-103_2
OX-BSI-506_2
OX-ENV-89_2
OX-BSI-145_2
OX-BSI-481_2
OX-BSI-523_2
OX-BSI-727_2
OX-LIV-266_2
OX-BSI-599_2
OX-BSI-102_2
OX-ENV-31_2
OX-ENV-33_2
OX-ENV-32_2
OX-BSI-499_2
OX-BSI-63_2
OX-BSI-273_2
OX-BSI-395_2
OX-LIV-361_2
OX-LIV-363_2
OX-BSI-718_2
OX-BSI-57_2
OX-BSI-691_2
OX-BSI-493_2
OX-BSI-458_2
OX-BSI-317_2
OX-BSI-468_2
OX-BSI-351_2
OX-WTW-2_2
OX-BSI-71_2
OX-BSI-657_2
OX-BSI-514_4
OX-BSI-199_2
OX-BSI-176_2
OX-BSI-669_2
OX-BSI-242_2
OX-BSI-267_2
OX-BSI-527_2
OX-BSI-33_2
OX-BSI-22_2
OX-BSI-683_2
OX-BSI-391_2
OX-BSI-631_2
OX-LIV-379_2
OX-LIV-362_2
OX-LIV-373_2
OX-BSI-182_2
OX-BSI-498_2
OX-BSI-632_2
OX-BSI-200_2
OX-BSI-239_3
OX-BSI-74_2
OX-BSI-311_2
OX-LIV-95_2
OX-LIV-99_2
OX-LIV-94_2
OX-LIV-100_2
OX-BSI-27_2
OX-BSI-173_2
OX-BSI-192_2
OX-BSI-25_2
OX-BSI-676_2
OX-LIV-117_2
OX-BSI-299_2
OX-BSI-592_2
OX-WTW-24_2
OX-BSI-638_2
OX-BSI-49_2
OX-BSI-87_2
OX-BSI-60_2
OX-BSI-394_2
OX-BSI-349_2
OX-BSI-339_2
OX-BSI-302_2
OX-BSI-470_2
OX-BSI-148_2
OX-BSI-664_3
OX-LIV-213_3
OX-LIV-118_2
OX-BSI-375_2
OX-BSI-8_2
OX-BSI-249_2
OX-BSI-656_2
OX-BSI-333_2
OX-BSI-731_2
OX-BSI-572_2
OX-BSI-720_2
OX-BSI-4_2
OX-BSI-496_2
0.000
0.025
0.050
0.075
0.100
No. nucleotide substitutions per site
Gene type
Absent
Core
Accessory
Transposase
Compartment
BSI
Environmental
Livestock
WwTW

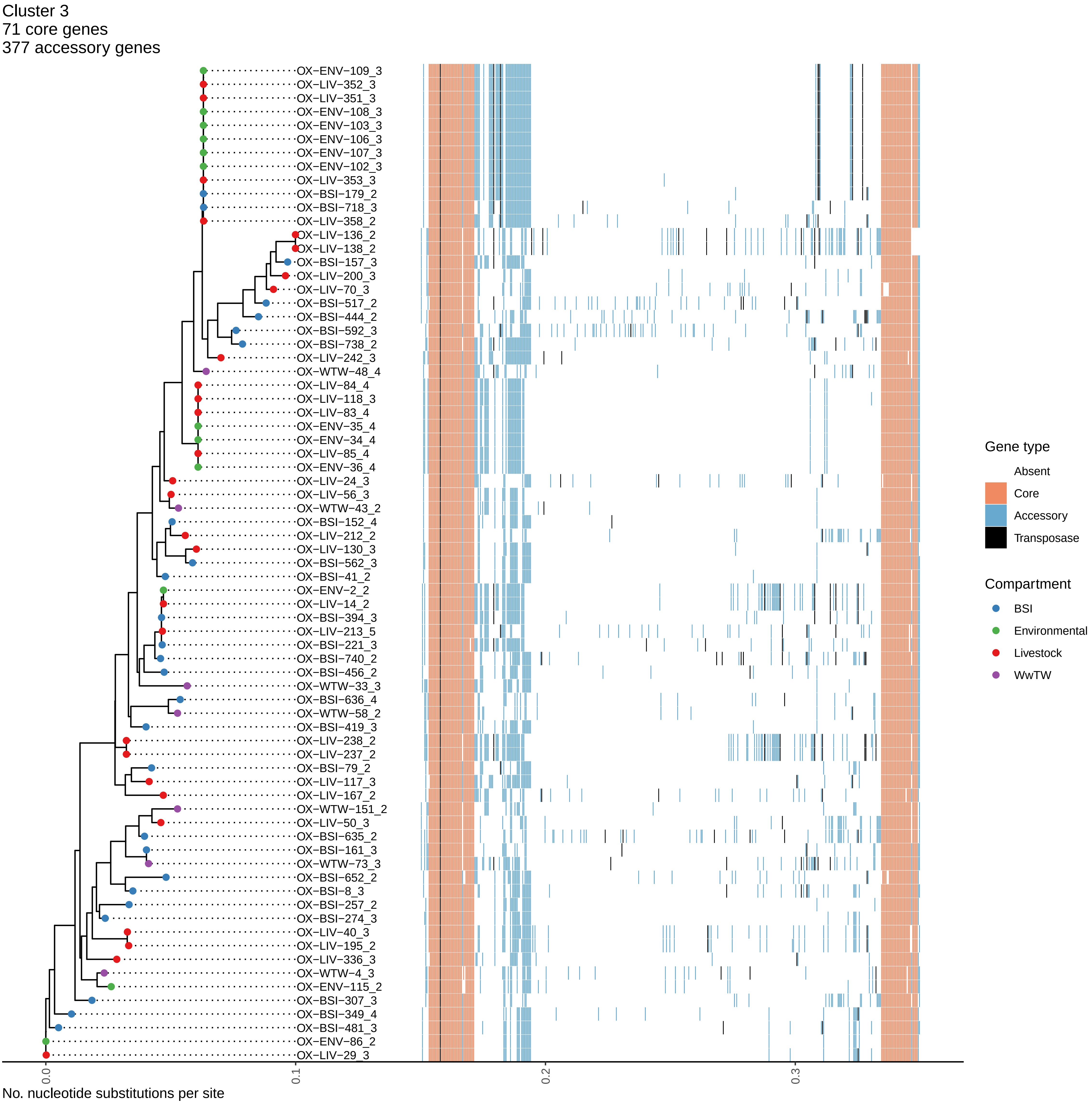
Cluster 3
71 core genes
377 accessory genes
OX−ENV−109_3
OX−LIV−352_3
OX−LIV−351_3
OX−ENV−108_3
OX−ENV−103_3
OX−ENV−106_3
OX−ENV−107_3
OX−ENV−102_3
OX−LIV−353_3
OX−BSI−179_2
OX−BSI−718_3
OX−LIV−358_2
OX−LIV−136_2
OX−LIV−138_2
OX−BSI−157_3
OX−LIV−200_3
OX−LIV−70_3
OX−BSI−517_2
OX−BSI−444_2
OX−BSI−592_3
OX−BSI−738_2
OX−LIV−242_3
OX−WTW−48_4
OX−LIV−84_4
OX−LIV−118_3
OX−LIV−83_4
OX−ENV−35_4
OX−ENV−34_4
OX−LIV−85_4
OX−ENV−36_4
OX−LIV−24_3
OX−LIV−56_3
OX−WTW−43_2
OX−BSI−152_4
OX−LIV−212_2
OX−LIV−130_3
OX−BSI−562_3
OX−BSI−41_2
OX−ENV−2_2
OX−LIV−14_2
OX−BSI−394_3
OX−LIV−213_5
OX−BSI−221_3
OX−BSI−740_2
OX−BSI−456_2
OX−WTW−33_3
OX−BSI−636_4
OX−WTW−58_2
OX−BSI−419_3
OX−LIV−238_2
OX−LIV−237_2
OX−BSI−79_2
OX−LIV−117_3
OX−LIV−167_2
OX−WTW−151_2
OX−LIV−50_3
OX−BSI−635_2
OX−BSI−161_3
OX−WTW−73_3
OX−BSI−652_2
OX−BSI−8_3
OX−BSI−257_2
OX−BSI−274_3
OX−LIV−40_3
OX−LIV−195_2
OX−LIV−336_3
OX−WTW−4_3
OX−ENV−115_2
OX−BSI−307_3
OX−BSI−349_4
OX−BSI−481_3
OX−ENV−86_2
OX−LIV−29_3
0.0
0.1
0.2
0.3
No. nucleotide substitutions per site
Gene type
Absent
Core
Accessory
Transposase
Compartment
BSI
Environmental
Livestock
WwTW

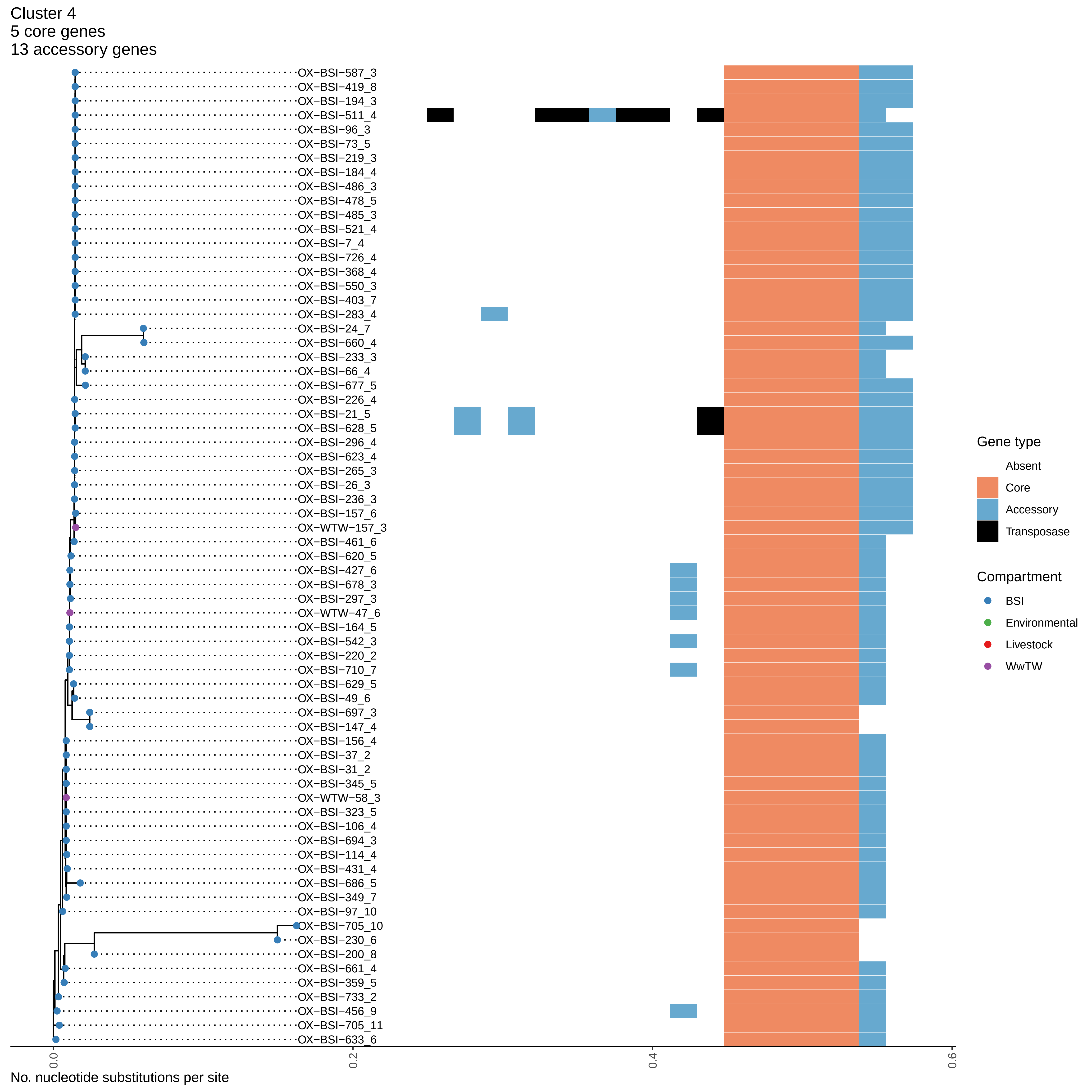
Cluster 4
5 core genes
13 accessory genes
OX−BSI−587_3
OX−BSI−419_8
OX−BSI−194_3
OX−BSI−511_4
OX−BSI−96_3
OX−BSI−73_5
OX−BSI−219_3
OX−BSI−184_4
OX−BSI−486_3
OX−BSI−478_5
OX−BSI−485_3
OX−BSI−521_4
OX−BSI−7_4
OX−BSI−726_4
OX−BSI−368_4
OX−BSI−550_3
OX−BSI−403_7
OX−BSI−283_4
OX−BSI−24_7
OX−BSI−660_4
OX−BSI−233_3
OX−BSI−66_4
OX−BSI−677_5
OX−BSI−226_4
OX−BSI−21_5
OX−BSI−628_5
OX−BSI−296_4
OX−BSI−623_4
OX−BSI−265_3
OX−BSI−26_3
OX−BSI−236_3
OX−BSI−157_6
OX−WTW−157_3
OX−BSI−461_6
OX−BSI−620_5
OX−BSI−427_6
OX−BSI−678_3
OX−BSI−297_3
OX−WTW−47_6
OX−BSI−164_5
OX−BSI−542_3
OX−BSI−220_2
OX−BSI−710_7
OX−BSI−629_5
OX−BSI−49_6
OX−BSI−697_3
OX−BSI−147_4
OX−BSI−156_4
OX−BSI−37_2
OX−BSI−31_2
OX−BSI−345_5
OX−WTW−58_3
OX−BSI−323_5
OX−BSI−106_4
OX−BSI−694_3
OX−BSI−114_4
OX−BSI−431_4
OX−BSI−686_5
OX−BSI−349_7
OX−BSI−97_10
OX−BSI−705_10
OX−BSI−230_6
OX−BSI−200_8
OX−BSI−661_4
OX−BSI−359_5
OX−BSI−733_2
OX−BSI−456_9
OX−BSI−705_11
OX−BSI−633_6
0.0
0.2
0.4
0.6
No. nucleotide substitutions per site
Gene type
Absent
Core
Accessory
Transposase
Compartment
BSI
Environmental
Livestock
WwTW

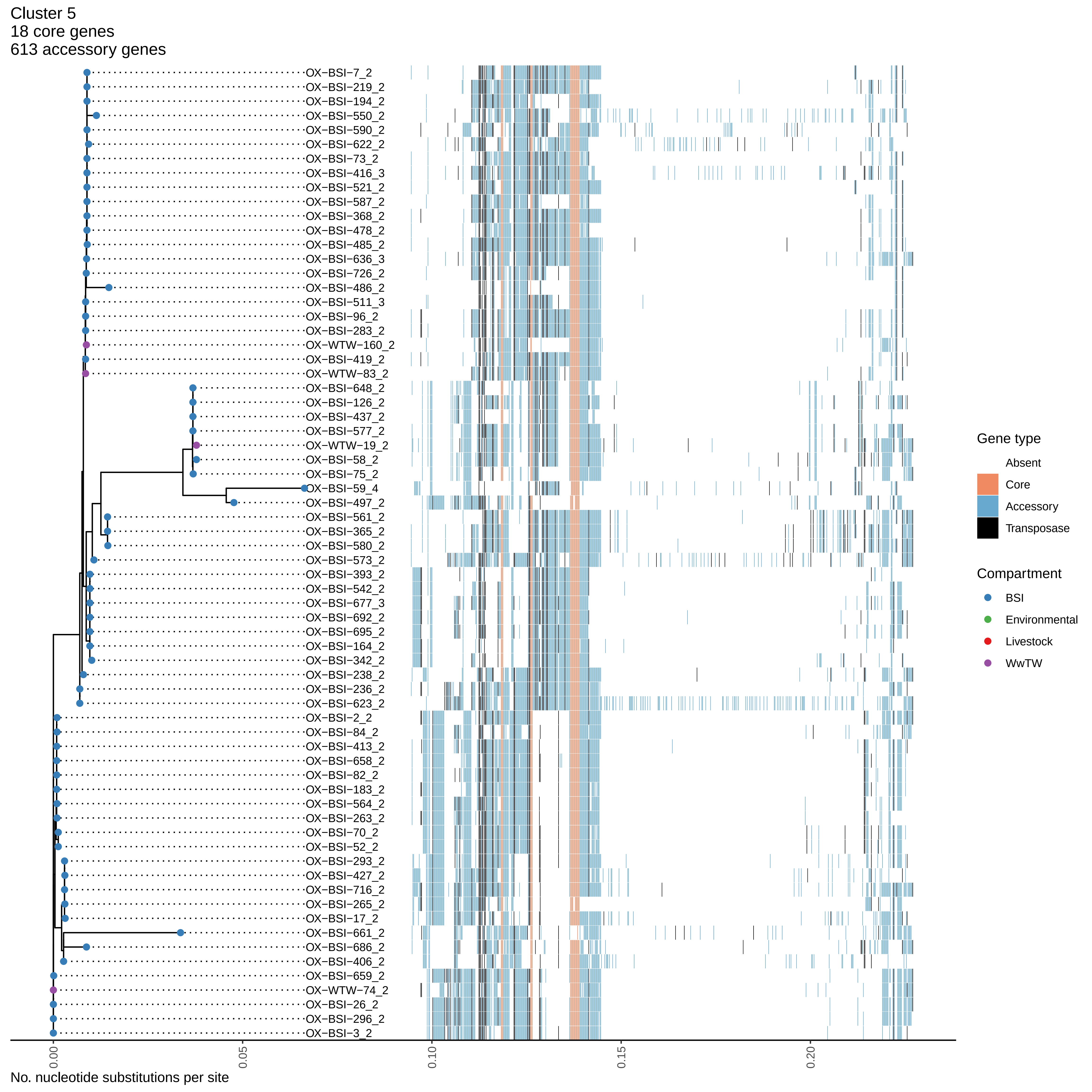

Cluster 5
18 core genes
613 accessory genes
OX−BSI−7_2
OX−BSI−219_2
OX−BSI−194_2
OX−BSI−550_2
OX−BSI−590_2
OX−BSI−622_2
OX−BSI−73_2
OX−BSI−416_3
OX−BSI−521_2
OX−BSI−587_2
OX−BSI−368_2
OX−BSI−478_2
OX−BSI−485_2
OX−BSI−636_3
OX−BSI−726_2
OX−BSI−486_2
OX−BSI−511_3
OX−BSI−96_2
OX−BSI−283_2
OX−WTW−160_2
OX−BSI−419_2
OX−WTW−83_2
OX−BSI−648_2
OX−BSI−126_2
OX−BSI−437_2
OX−BSI−577_2
OX−WTW−19_2
OX−BSI−58_2
OX−BSI−75_2
OX−BSI−59_4
OX−BSI−497_2
OX−BSI−561_2
OX−BSI−365_2
OX−BSI−580_2
OX−BSI−573_2
OX−BSI−393_2
OX−BSI−542_2
OX−BSI−677_3
OX−BSI−692_2
OX−BSI−695_2
OX−BSI−164_2
OX−BSI−342_2
OX−BSI−238_2
OX−BSI−236_2
OX−BSI−623_2
OX−BSI−2_2
OX−BSI−84_2
OX−BSI−413_2
OX−BSI−658_2
OX−BSI−82_2
OX−BSI−183_2
OX−BSI−564_2
OX−BSI−263_2
OX−BSI−70_2
OX−BSI−52_2
OX−BSI−293_2
OX−BSI−427_2
OX−BSI−716_2
OX−BSI−265_2
OX−BSI−17_2
OX−BSI−661_2
OX−BSI−686_2
OX−BSI−406_2
OX−BSI−659_2
OX−WTW−74_2
OX−BSI−26_2
OX−BSI−296_2
OX−BSI−3_2
Gene type
Absent
Core
Accessory
Transposase
Compartment
BSI
Environmental
Livestock
WwTW
0.00
0.05
0.10
0.15
0.20
No. nucleotide substitutions per site

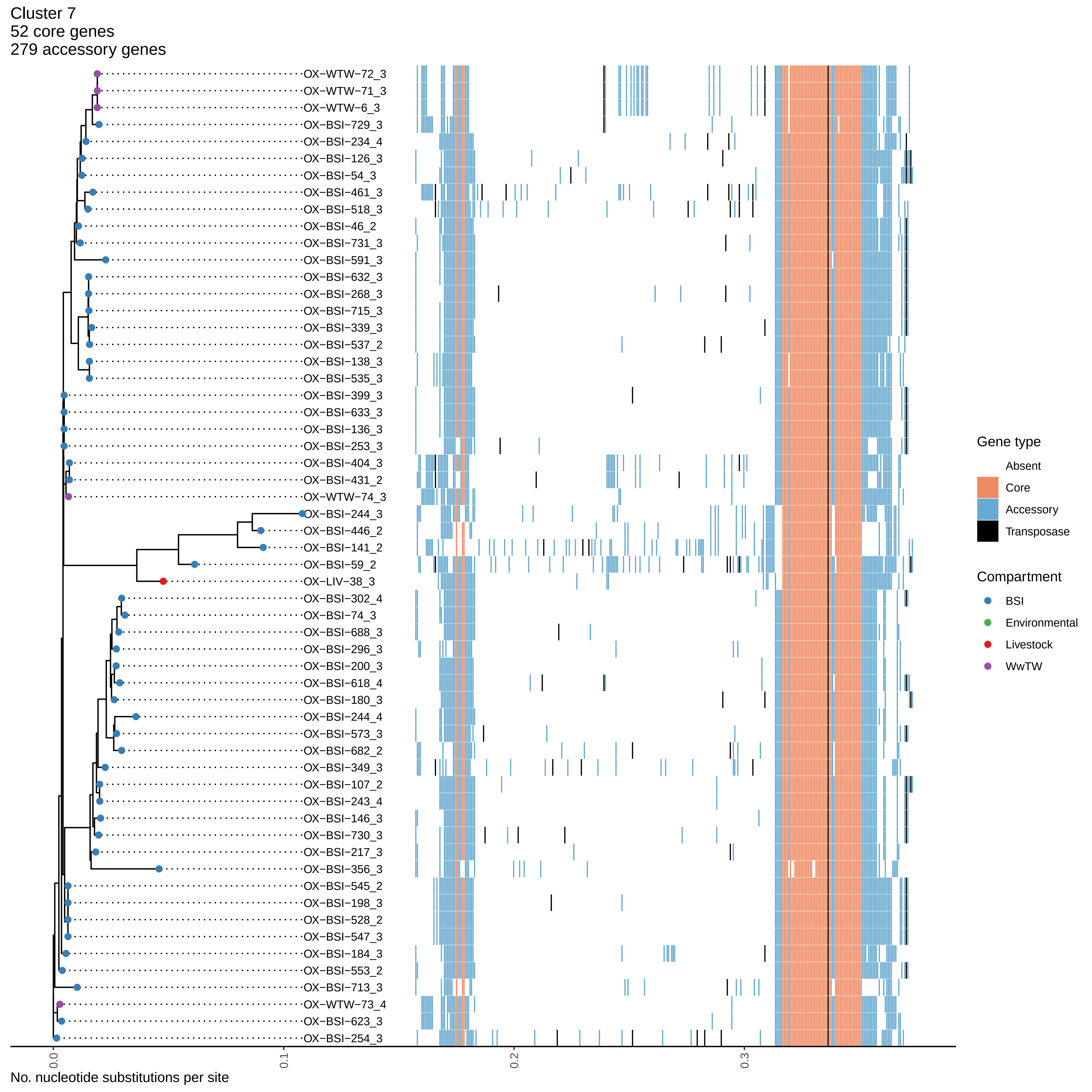
Cluster 7
52 core genes
279 accessory genes
OX−WTW−72_3
OX−WTW−71_3
OX−WTW−6_3
OX−BSI−729_3
OX−BSI−234_4
OX−BSI−126_3
OX−BSI−54_3
OX−BSI−461_3
OX−BSI−518_3
OX−BSI−46_2
OX−BSI−731_3
OX−BSI−591_3
OX−BSI−632_3
OX−BSI−268_3
OX−BSI−715_3
OX−BSI−339_3
OX−BSI−537_2
OX−BSI−138_3
OX−BSI−535_3
OX−BSI−399_3
OX−BSI−633_3
OX−BSI−136_3
OX−BSI−253_3
OX−BSI−404_3
OX−BSI−431_2
OX−WTW−74_3
OX−BSI−244_3
OX−BSI−446_2
OX−BSI−141_2
OX−BSI−59_2
OX−LIV−38_3
OX−BSI−302_4
OX−BSI−74_3
OX−BSI−688_3
OX−BSI−296_3
OX−BSI−200_3
OX−BSI−618_4
OX−BSI−180_3
OX−BSI−244_4
OX−BSI−573_3
OX−BSI−682_2
OX−BSI−349_3
OX−BSI−107_2
OX−BSI−243_4
OX−BSI−146_3
OX−BSI−730_3
OX−BSI−217_3
OX−BSI−356_3
OX−BSI−545_2
OX−BSI−198_3
OX−BSI−528_2
OX−BSI−547_3
OX−BSI−184_3
OX−BSI−553_2
OX−BSI−713_3
OX−WTW−73_4
OX−BSI−623_3
OX−BSI−254_3
0.0
0.1
0.2
0.3
No. nucleotide substitutions per site
Gene type
Absent
Core
Accessory
Transposase
Compartment
BSI
Environmental
Livestock
WwTW

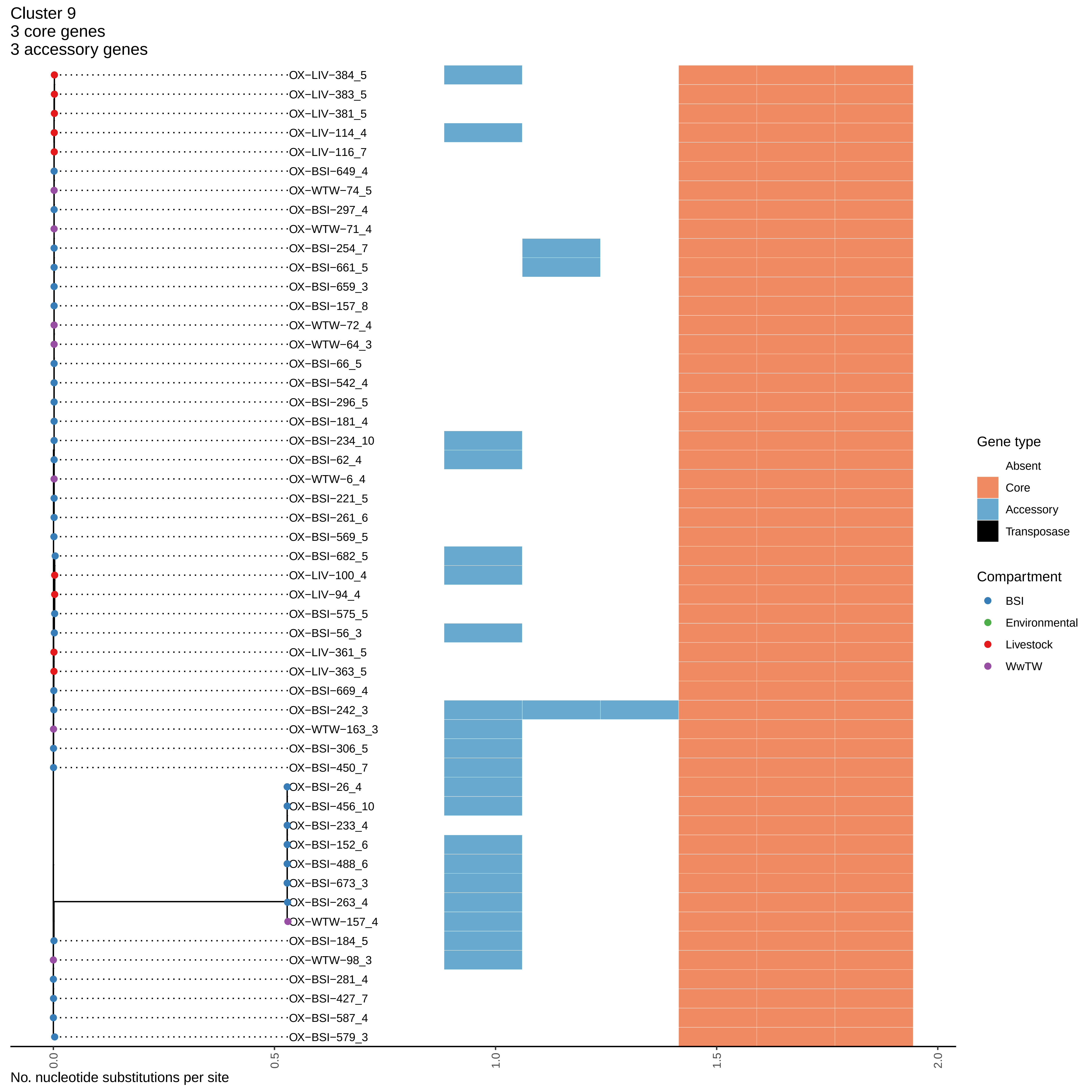

Cluster 9
3 core genes
3 accessory genes
OX−LIV−384_5
OX−LIV−383_5
OX−LIV−381_5
OX−LIV−114_4
OX−LIV−116_7
OX−BSI−649_4
OX−WTW−74_5
OX−BSI−297_4
OX−WTW−71_4
OX−BSI−254_7
OX−BSI−661_5
OX−BSI−659_3
OX−BSI−157_8
OX−WTW−72_4
OX−WTW−64_3
OX−BSI−66_5
OX−BSI−542_4
OX−BSI−296_5
OX−BSI−181_4
OX−BSI−234_10
OX−BSI−62_4
OX−WTW−6_4
OX−BSI−221_5
OX−BSI−261_6
OX−BSI−569_5
OX−BSI−682_5
OX−LIV−100_4
OX−LIV−94_4
OX−BSI−575_5
OX−BSI−56_3
OX−LIV−361_5
OX−LIV−363_5
OX−BSI−669_4
OX−BSI−242_3
OX−WTW−163_3
OX−BSI−306_5
OX−BSI−450_7
OX−BSI−26_4
OX−BSI−456_10
OX−BSI−233_4
OX−BSI−152_6
OX−BSI−488_6
OX−BSI−673_3
OX−BSI−263_4
OX−WTW−157_4
OX−BSI−184_5
OX−WTW−98_3
OX−BSI−281_4
OX−BSI−427_7
OX−BSI−587_4
OX−BSI−579_3
0.0
0.5
1.0
1.5
2.0
No. nucleotide substitutions per site
Gene type
Absent
Core
Accessory
Transposase
Compartment
BSI
Environmental
Livestock
WwTW

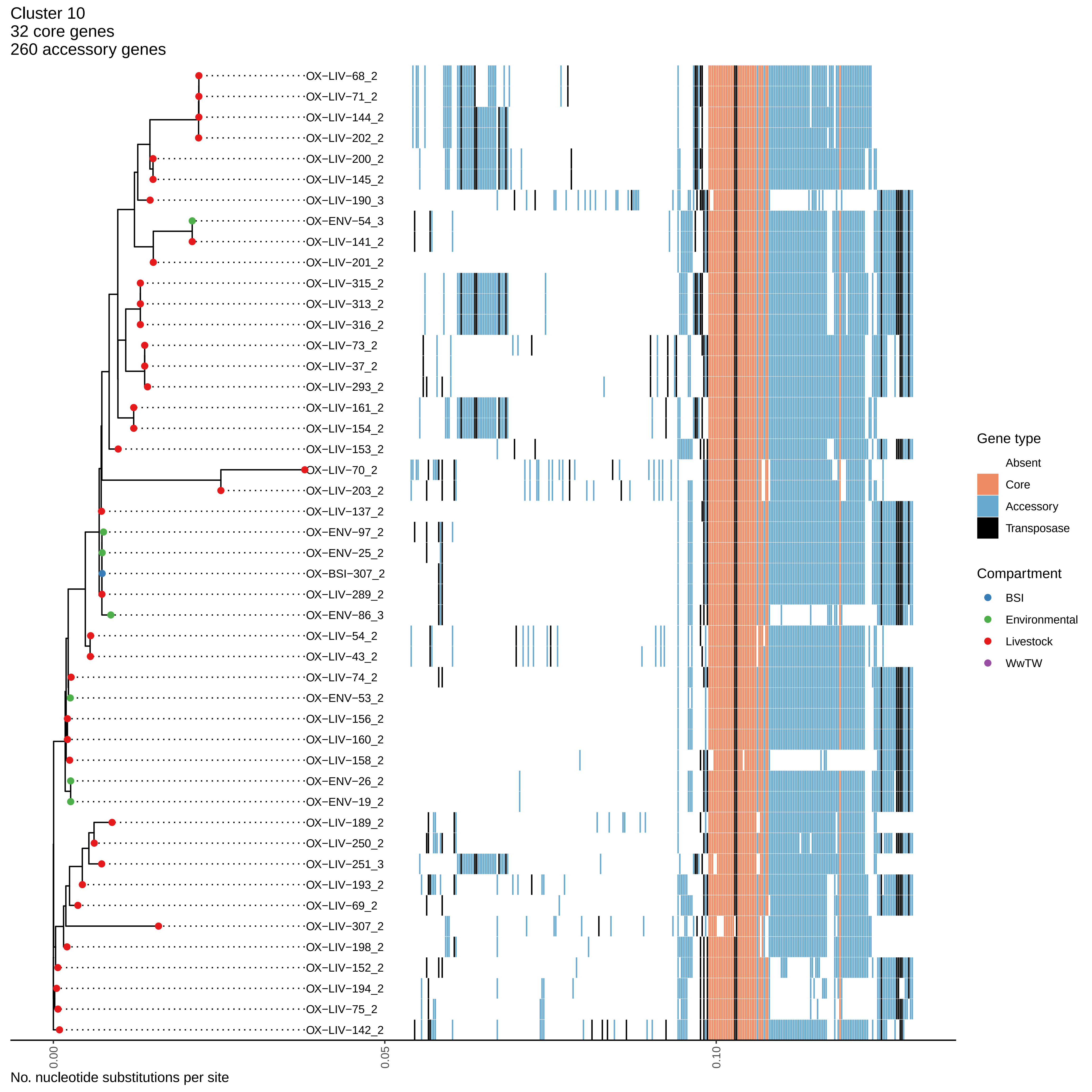

Cluster 10
32 core genes
260 accessory genes
OX−LIV−68_2
OX−LIV−71_2
OX−LIV−144_2
OX−LIV−202_2
OX−LIV−200_2
OX−LIV−145_2
OX−LIV−190_3
OX−ENV−54_3
OX−LIV−141_2
OX−LIV−201_2
OX−LIV−315_2
OX−LIV−313_2
OX−LIV−316_2
OX−LIV−73_2
OX−LIV−37_2
OX−LIV−293_2
OX−LIV−161_2
OX−LIV−154_2
OX−LIV−153_2
OX−LIV−70_2
OX−LIV−203_2
OX−LIV−137_2
OX−ENV−97_2
OX−ENV−25_2
OX−BSI−307_2
OX−LIV−289_2
OX−ENV−86_3
OX−LIV−54_2
OX−LIV−43_2
OX−LIV−74_2
OX−ENV−53_2
OX−LIV−156_2
OX−LIV−160_2
OX−LIV−158_2
OX−ENV−26_2
OX−ENV−19_2
OX−LIV−189_2
OX−LIV−250_2
OX−LIV−251_3
OX−LIV−193_2
OX−LIV−69_2
OX−LIV−307_2
OX−LIV−198_2
OX−LIV−152_2
OX−LIV−194_2
OX−LIV−75_2
OX−LIV−142_2
0.00
0.05
0.10
No. nucleotide substitutions per site
Gene type
Absent
Core
Accessory
Transposase
Compartment
BSI
Environmental
Livestock
WwTW

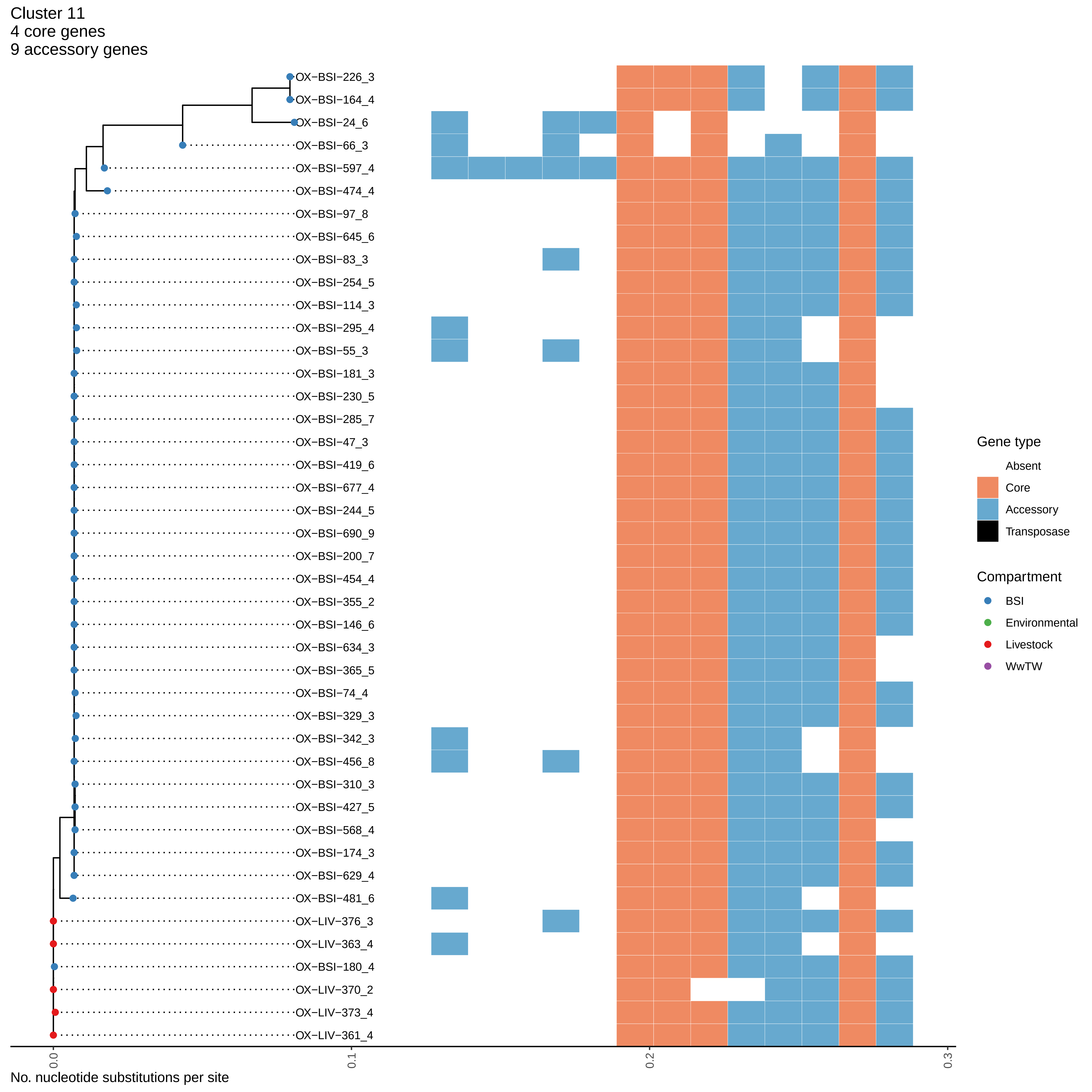

Cluster 11
4 core genes
9 accessory genes
OX−BSI−226_3
OX−BSI−164_4
OX−BSI−24_6
OX−BSI−66_3
OX−BSI−597_4
OX−BSI−474_4
OX−BSI−97_8
OX−BSI−645_6
OX−BSI−83_3
OX−BSI−254_5
OX−BSI−114_3
OX−BSI−295_4
OX−BSI−55_3
OX−BSI−181_3
OX−BSI−230_5
OX−BSI−285_7
OX−BSI−47_3
OX−BSI−419_6
OX−BSI−677_4
OX−BSI−244_5
OX−BSI−690_9
OX−BSI−200_7
OX−BSI−454_4
OX−BSI−355_2
OX−BSI−146_6
OX−BSI−634_3
OX−BSI−365_5
OX−BSI−74_4
OX−BSI−329_3
OX−BSI−342_3
OX−BSI−456_8
OX−BSI−310_3
OX−BSI−427_5
OX−BSI−568_4
OX−BSI−174_3
OX−BSI−629_4
OX−BSI−481_6
OX−LIV−376_3
OX−LIV−363_4
OX−BSI−180_4
OX−LIV−370_2
OX−LIV−373_4
OX−LIV−361_4
Gene type
Absent
Core
Accessory
Transposase
Compartment
BSI
Environmental
Livestock
WwTW
0.0
0.1
0.2
0.3
No. nucleotide substitutions per site

Cluster 12
49 core genes
198 accessory genes

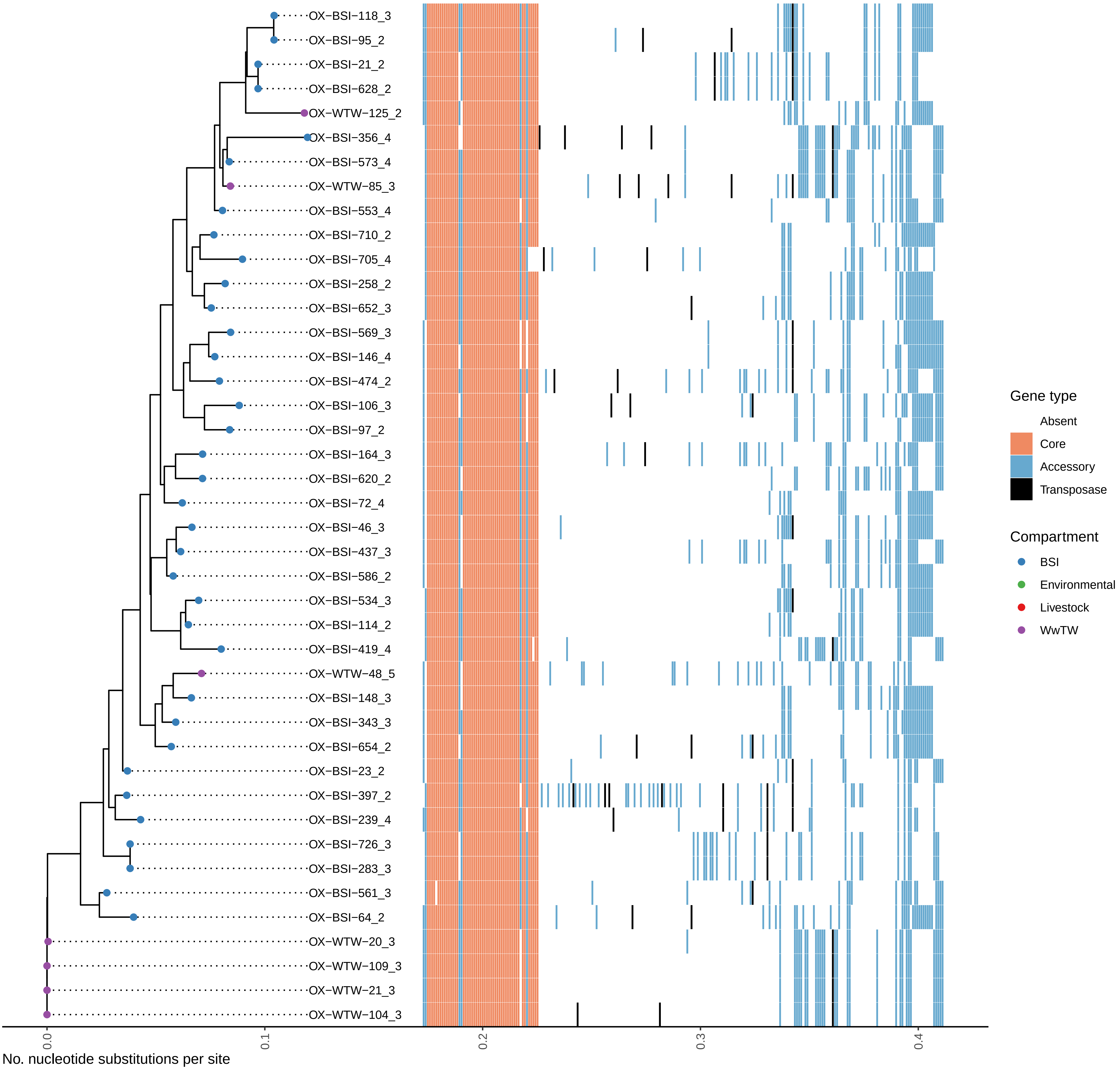

Cluster 13
117 core genes
86 accessory genes

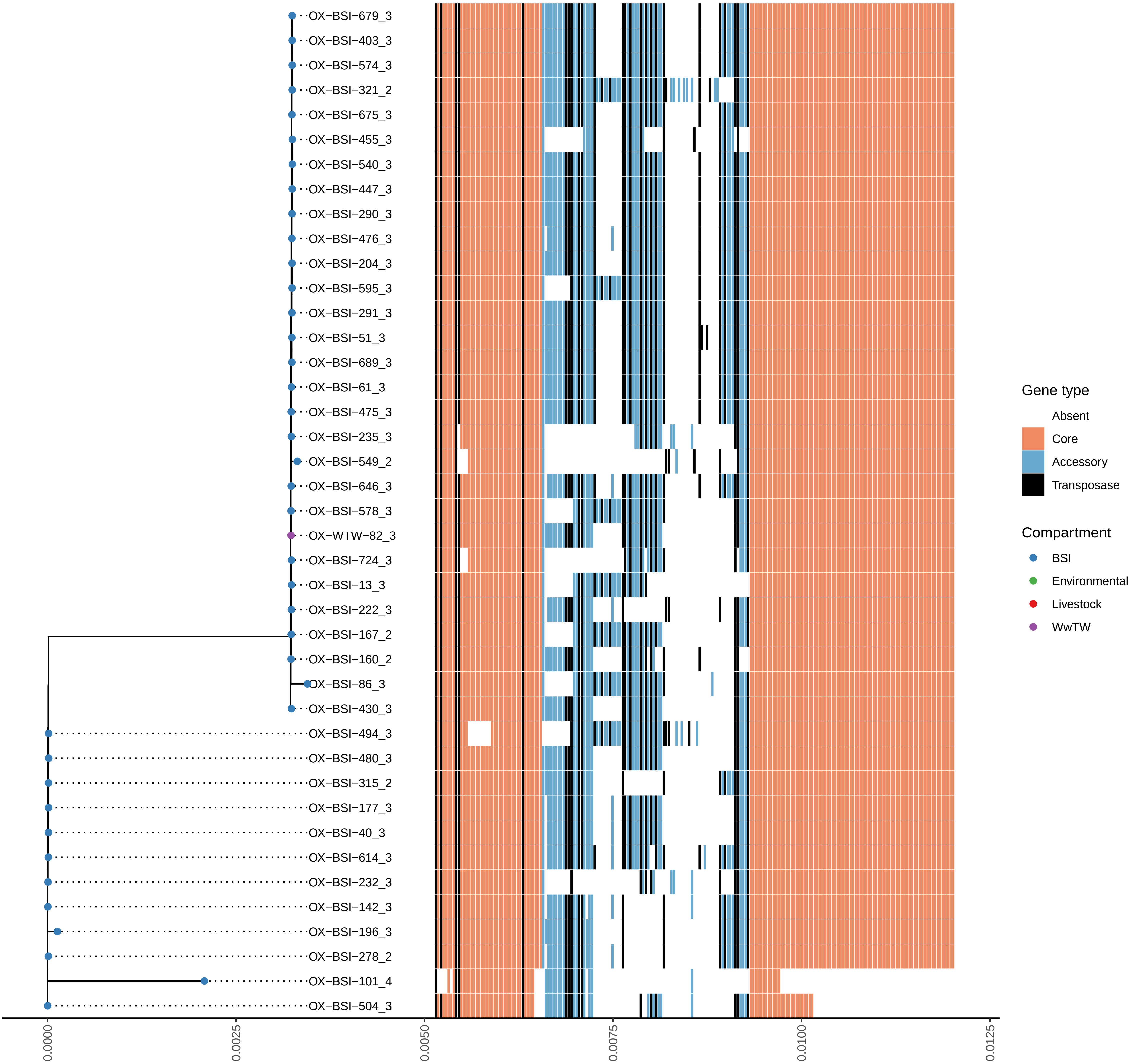

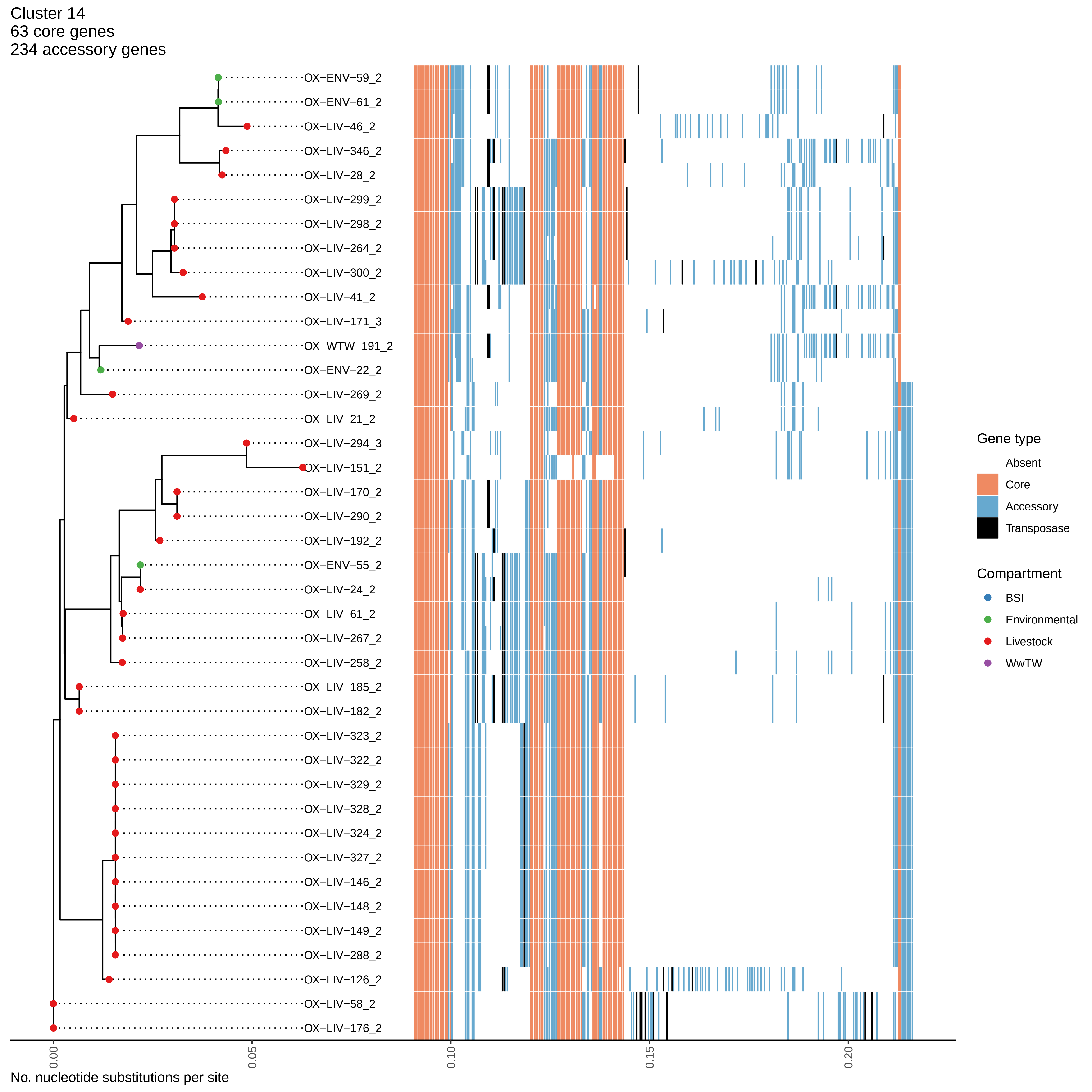

Cluster 14
63 core genes
234 accessory genes
OX−ENV−59_2
OX−ENV−61_2
OX−LIV−46_2
OX−LIV−346_2
OX−LIV−28_2
OX−LIV−299_2
OX−LIV−298_2
OX−LIV−264_2
OX−LIV−300_2
OX−LIV−41_2
OX−LIV−171_3
OX−WTW−191_2
OX−ENV−22_2
OX−LIV−269_2
OX−LIV−21_2
OX−LIV−294_3
OX−LIV−151_2
OX−LIV−170_2
OX−LIV−290_2
OX−LIV−192_2
OX−ENV−55_2
OX−LIV−24_2
OX−LIV−61_2
OX−LIV−267_2
OX−LIV−258_2
OX−LIV−185_2
OX−LIV−182_2
OX−LIV−323_2
OX−LIV−322_2
OX−LIV−329_2
OX−LIV−328_2
OX−LIV−324_2
OX−LIV−327_2
OX−LIV−146_2
OX−LIV−148_2
OX−LIV−149_2
OX−LIV−288_2
OX−LIV−126_2
OX−LIV−58_2
OX−LIV−176_2
0.00
0.05
0.10
0.15
0.20
No. nucleotide substitutions per site
Gene type
Absent
Core
Accessory
Transposase
Compartment
BSI
Environmental
Livestock
WwTW

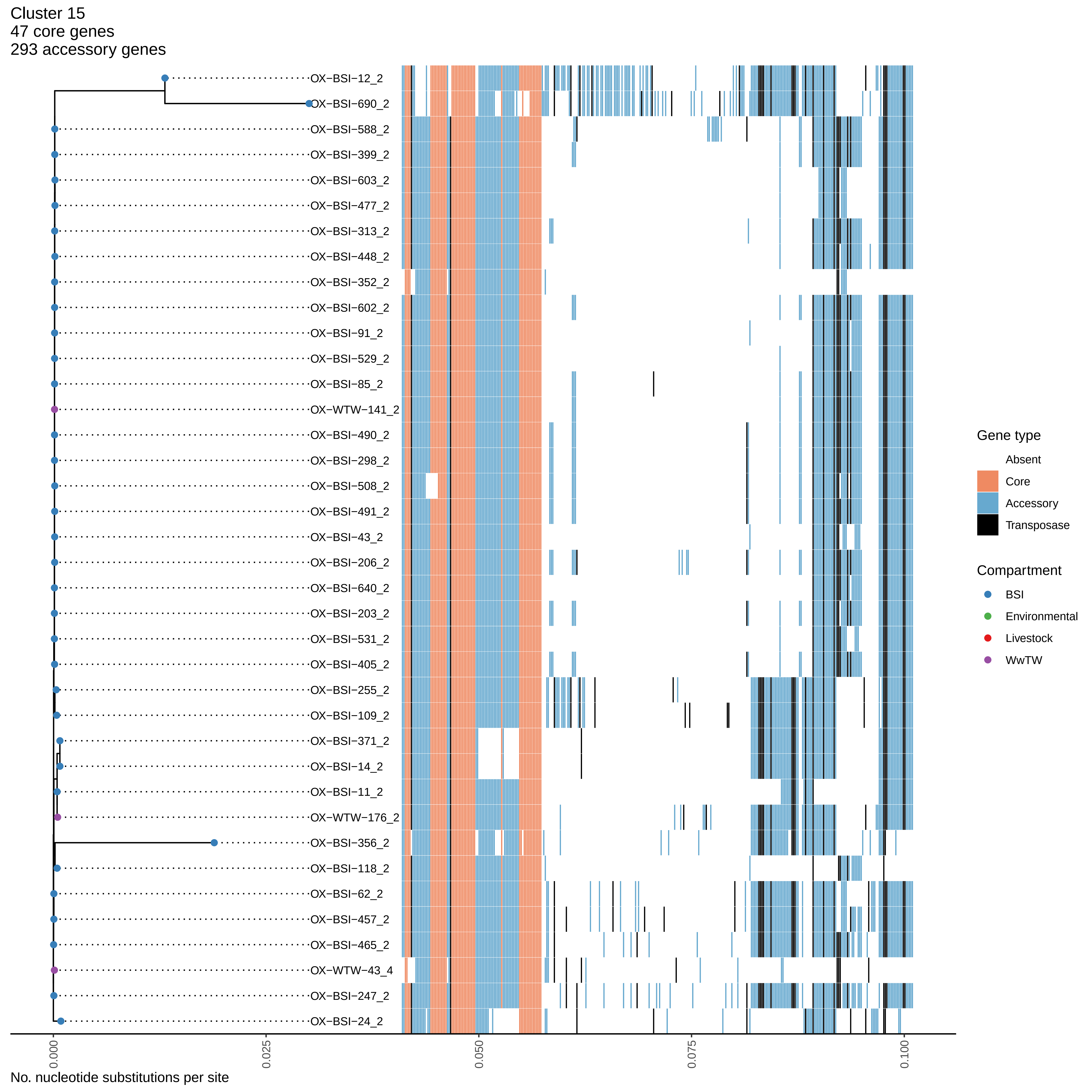

Cluster 15
47 core genes
293 accessory genes
OX−BSI−12_2
OX−BSI−690_2
OX−BSI−588_2
OX−BSI−399_2
OX−BSI−603_2
OX−BSI−477_2
OX−BSI−313_2
OX−BSI−448_2
OX−BSI−352_2
OX−BSI−602_2
OX−BSI−91_2
OX−BSI−529_2
OX−BSI−85_2
OX−WTW−141_2
OX−BSI−490_2
OX−BSI−298_2
OX−BSI−508_2
OX−BSI−491_2
OX−BSI−43_2
OX−BSI−206_2
OX−BSI−640_2
OX−BSI−203_2
OX−BSI−531_2
OX−BSI−405_2
OX−BSI−255_2
OX−BSI−109_2
OX−BSI−371_2
OX−BSI−14_2
OX−BSI−11_2
OX−WTW−176_2
OX−BSI−356_2
OX−BSI−118_2
OX−BSI−62_2
OX−BSI−457_2
OX−BSI−465_2
OX−WTW−43_4
OX−BSI−247_2
OX−BSI−24_2
0.000
0.025
0.050
0.075
0.100
No. nucleotide substitutions per site
Gene type
Absent
Core
Accessory
Transposase
Compartment
BSI
Environmental
Livestock
WwTW

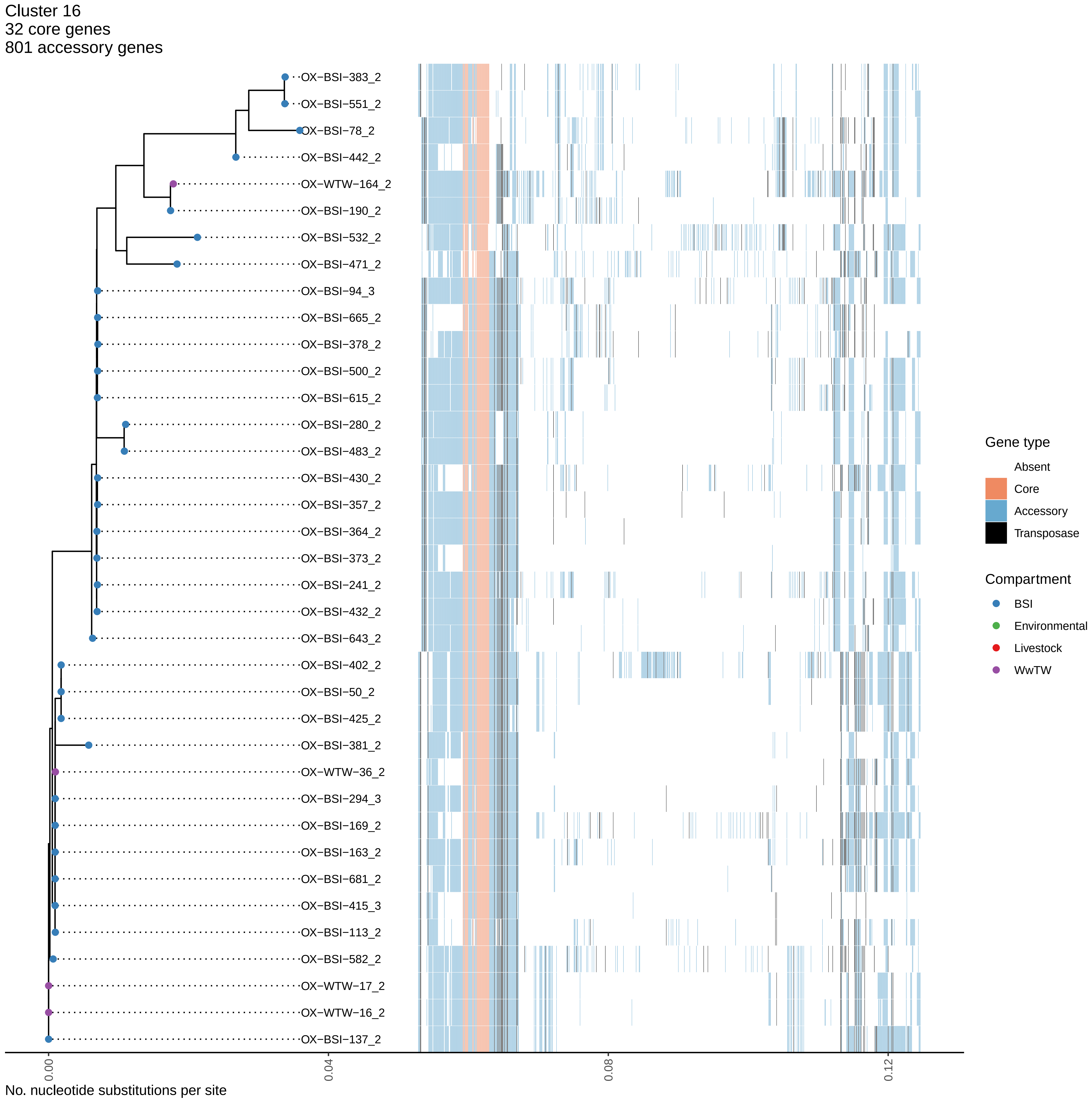
Cluster 16
32 core genes
801 accessory genes
OX−BSI−383_2
OX−BSI−551_2
OX−BSI−78_2
OX−BSI−442_2
OX−WTW−164_2
OX−BSI−190_2
OX−BSI−532_2
OX−BSI−471_2
OX−BSI−94_3
OX−BSI−665_2
OX−BSI−378_2
OX−BSI−500_2
OX−BSI−615_2
OX−BSI−280_2
OX−BSI−483_2
OX−BSI−430_2
OX−BSI−357_2
OX−BSI−364_2
OX−BSI−373_2
OX−BSI−241_2
OX−BSI−432_2
OX−BSI−643_2
OX−BSI−402_2
OX−BSI−50_2
OX−BSI−425_2
OX−BSI−381_2
OX−WTW−36_2
OX−BSI−294_3
OX−BSI−169_2
OX−BSI−163_2
OX−BSI−681_2
OX−BSI−415_3
OX−BSI−113_2
OX−BSI−582_2
OX−WTW−17_2
OX−WTW−16_2
OX−BSI−137_2
0.00
0.04
0.08
0.12
No. nucleotide substitutions per site
Gene type
Absent
Core
Accessory
Transposase
Compartment
BSI
Environmental
Livestock
WwTW

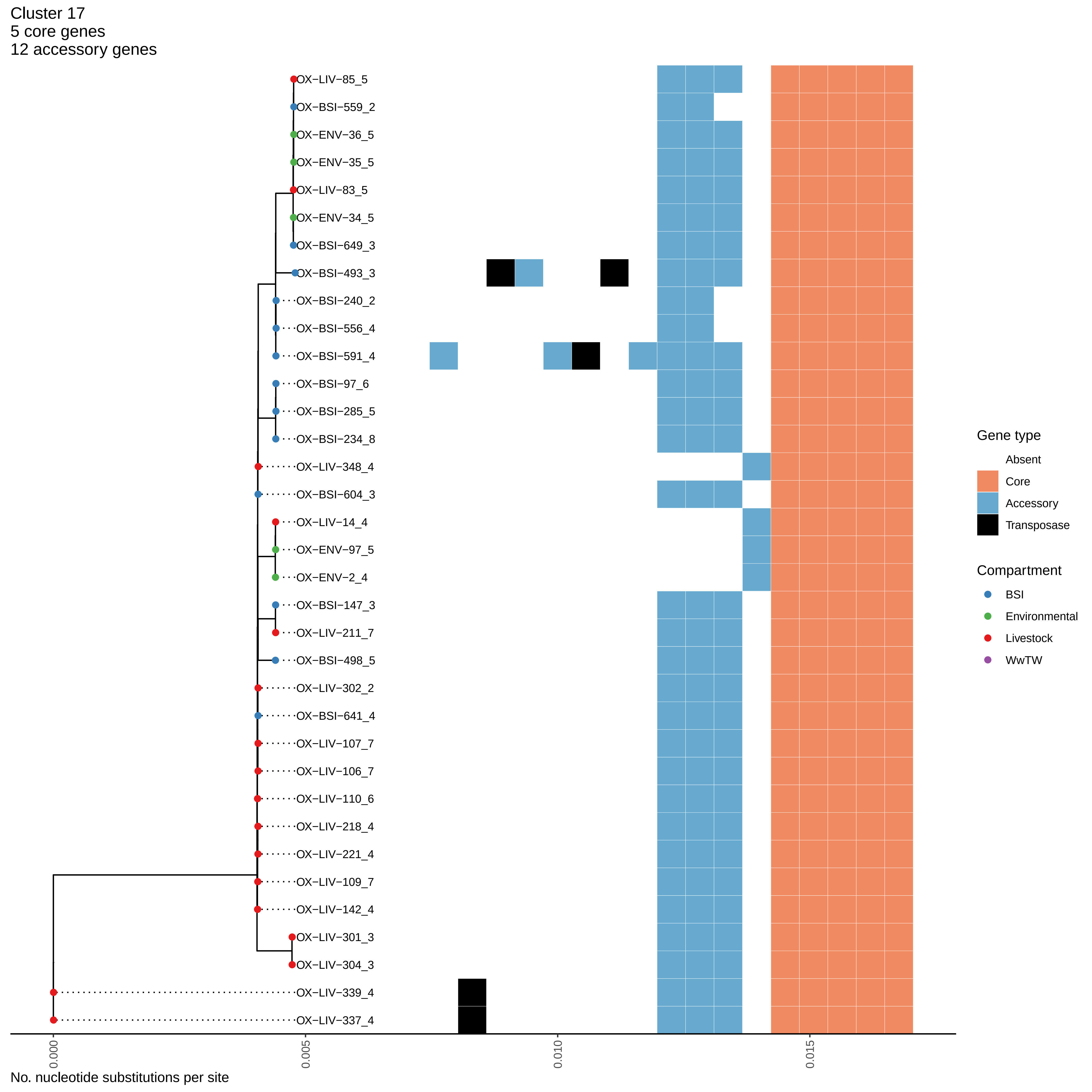
Cluster 17
5 core genes
12 accessory genes
OX−LIV−85_5
OX−BSI−559_2
OX−ENV−36_5
OX−ENV−35_5
OX−LIV−83_5
OX−ENV−34_5
OX−BSI−649_3
OX−BSI−493_3
OX−BSI−240_2
OX−BSI−556_4
OX−BSI−591_4
OX−BSI−97_6
OX−BSI−285_5
OX−BSI−234_8
OX−LIV−348_4
OX−BSI−604_3
OX−LIV−14_4
OX−ENV−97_5
OX−ENV−2_4
OX−BSI−147_3
OX−LIV−211_7
OX−BSI−498_5
OX−LIV−302_2
OX−BSI−641_4
OX−LIV−107_7
OX−LIV−106_7
OX−LIV−110_6
OX−LIV−218_4
OX−LIV−221_4
OX−LIV−109_7
OX−LIV−142_4
OX−LIV−301_3
OX−LIV−304_3
OX−LIV−339_4
OX−LIV−337_4
0.000
0.005
0.010
0.015
No. nucleotide substitutions per site
Gene type
Absent
Core
Accessory
Transposase
Compartment
BSI
Environmental
Livestock
WwTW

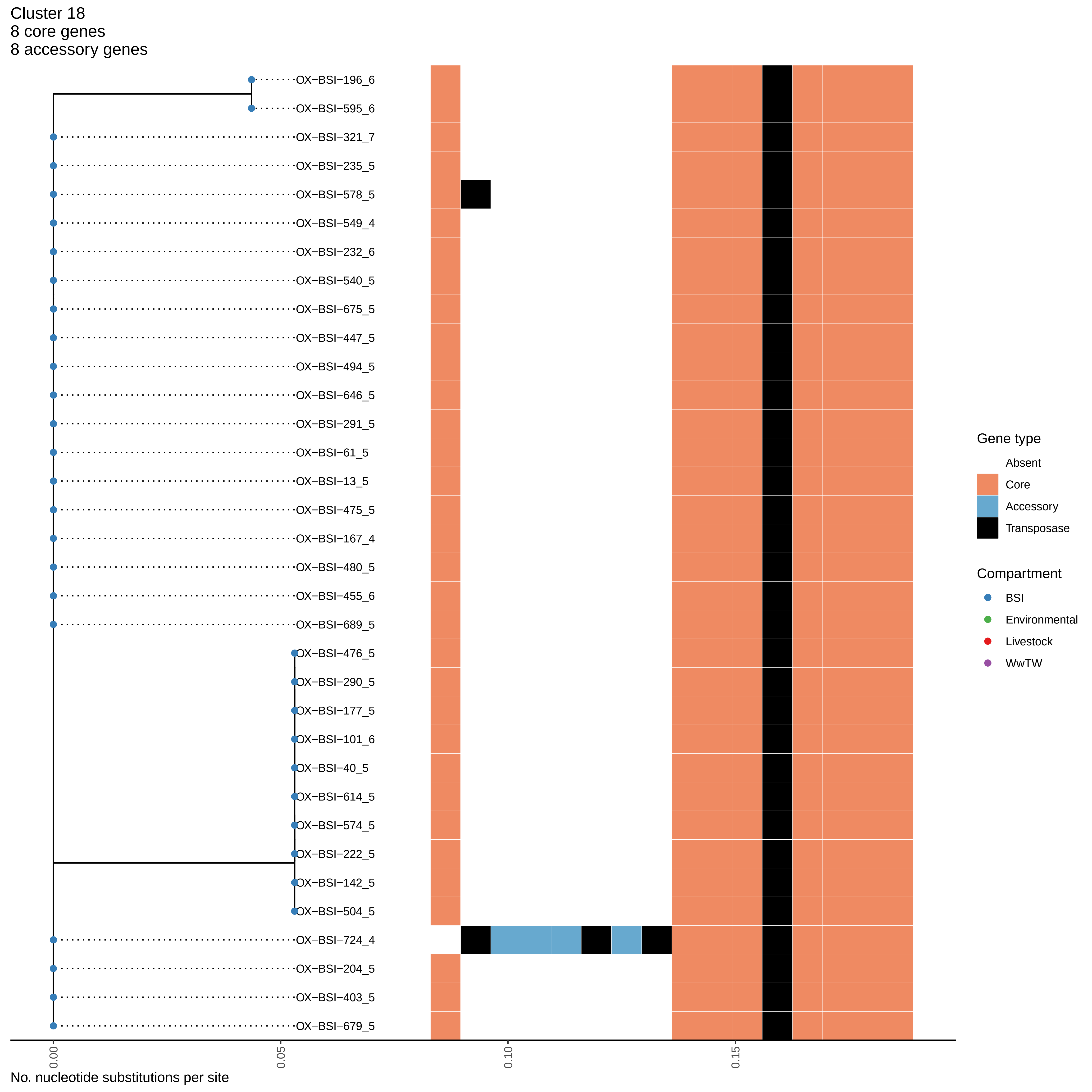
Cluster 18
8 core genes
8 accessory genes
OX−BSI−196_6
OX−BSI−595_6
OX−BSI−321_7
OX−BSI−235_5
OX−BSI−578_5
OX−BSI−549_4
OX−BSI−232_6
OX−BSI−540_5
OX−BSI−675_5
OX−BSI−447_5
OX−BSI−494_5
OX−BSI−646_5
OX−BSI−291_5
OX−BSI−61_5
OX−BSI−13_5
OX−BSI−475_5
OX−BSI−167_4
OX−BSI−480_5
OX−BSI−455_6
OX−BSI−689_5
OX−BSI−476_5
OX−BSI−290_5
OX−BSI−177_5
OX−BSI−101_6
OX−BSI−40_5
OX−BSI−614_5
OX−BSI−574_5
OX−BSI−222_5
OX−BSI−142_5
OX−BSI−504_5
OX−BSI−724_4
OX−BSI−204_5
OX−BSI−403_5
OX−BSI−679_5
0.00
0.05
0.10
0.15
No. nucleotide substitutions per site
Gene type
Absent
Core
Accessory
Transposase
Compartment
BSI
Environmental
Livestock
WwTW

Cluster 19
116 core genes
173 accessory genes

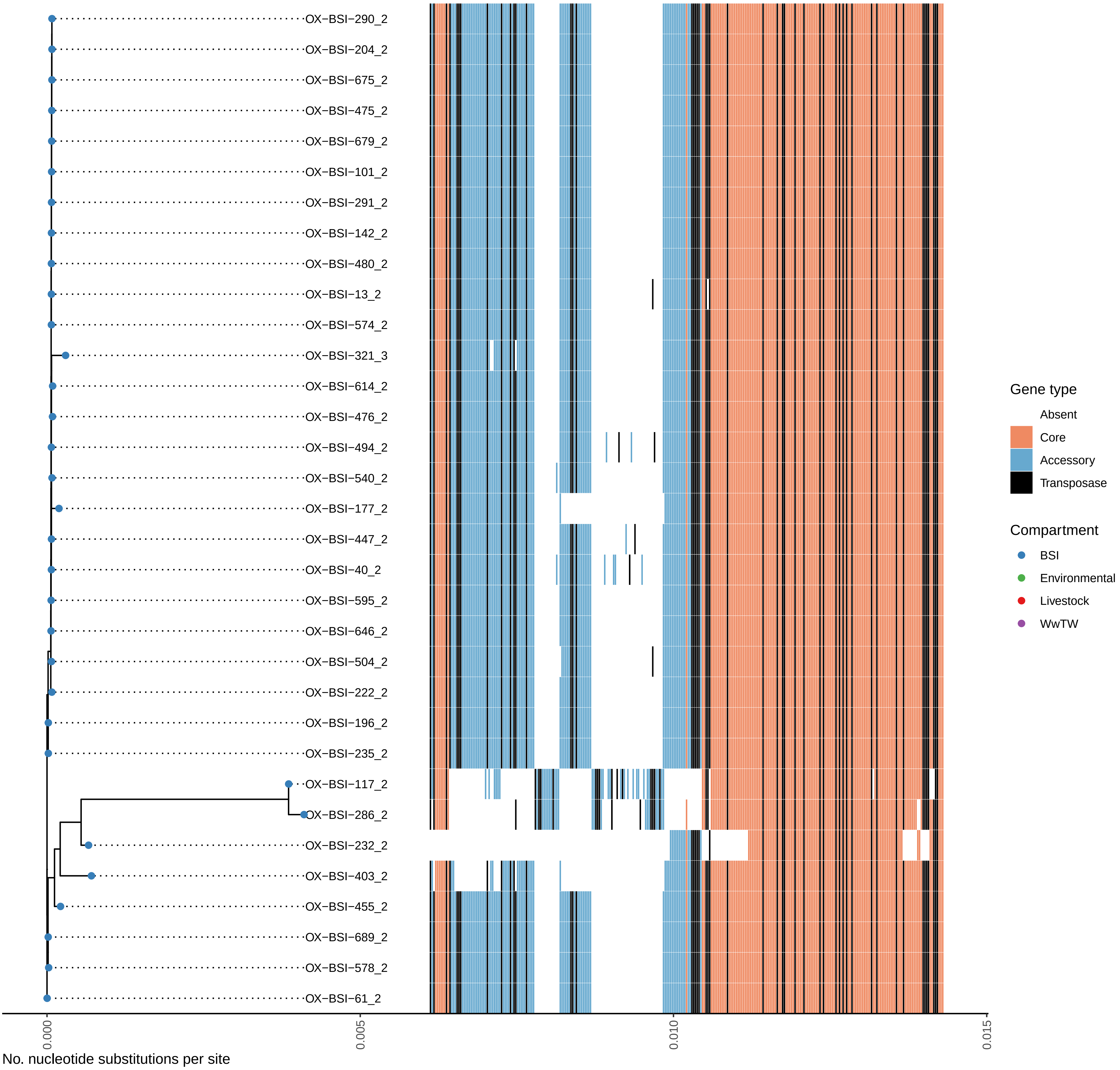

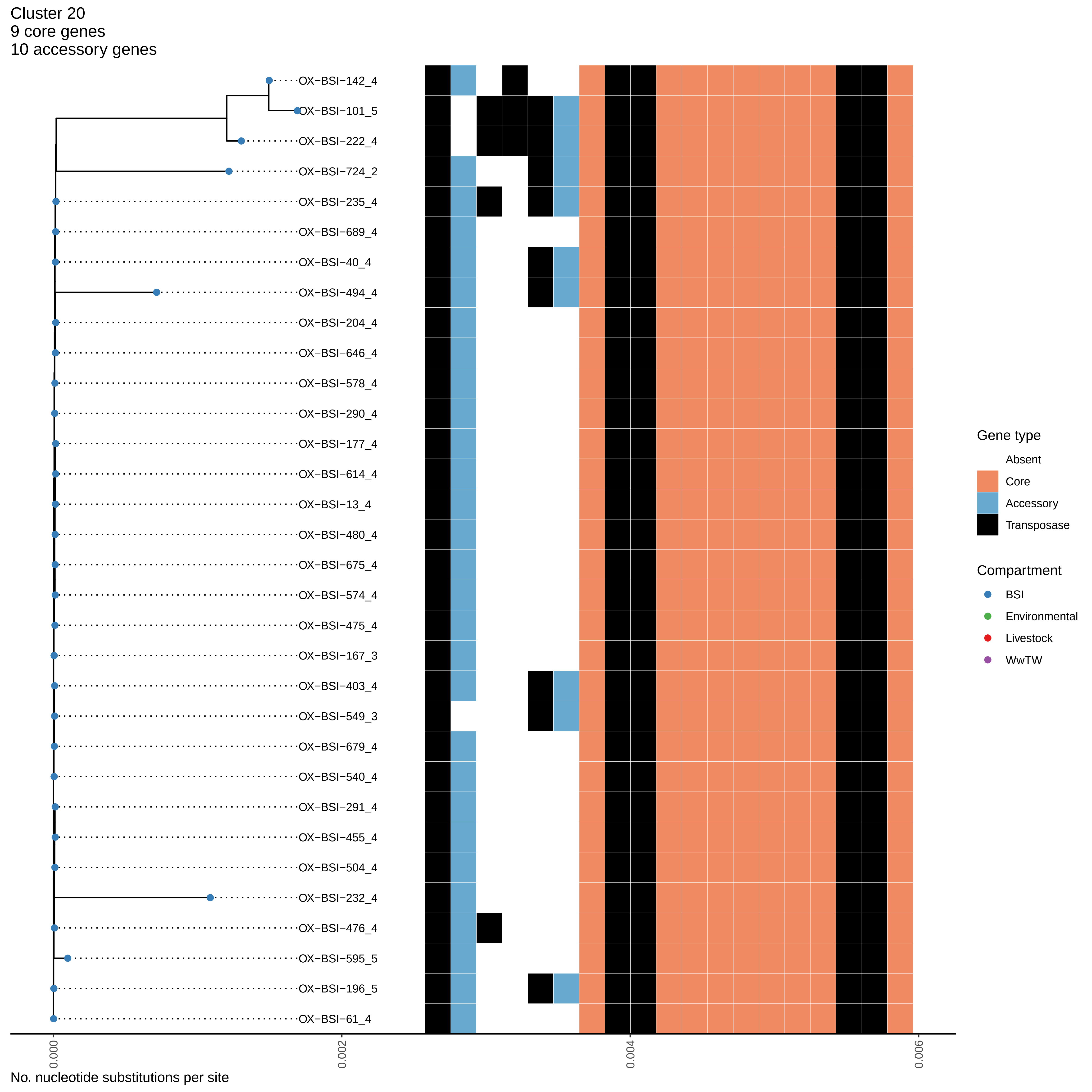

Cluster 20
9 core genes
10 accessory genes
OX−BSI−142_4
OX−BSI−101_5
OX−BSI−222_4
OX−BSI−724_2
OX−BSI−235_4
OX−BSI−689_4
OX−BSI−40_4
OX−BSI−494_4
OX−BSI−204_4
OX−BSI−646_4
OX−BSI−578_4
OX−BSI−290_4
OX−BSI−177_4
OX−BSI−614_4
OX−BSI−13_4
OX−BSI−480_4
OX−BSI−675_4
OX−BSI−574_4
OX−BSI−475_4
OX−BSI−167_3
OX−BSI−403_4
OX−BSI−549_3
OX−BSI−679_4
OX−BSI−540_4
OX−BSI−291_4
OX−BSI−455_4
OX−BSI−504_4
OX−BSI−232_4
OX−BSI−476_4
OX−BSI−595_5
OX−BSI−196_5
OX−BSI−61_4
0.000
0.002
0.004
0.006
No. nucleotide substitutions per site
Gene type
Absent
Core
Accessory
Transposase
Compartment
BSI
Environmental
Livestock
WwTW

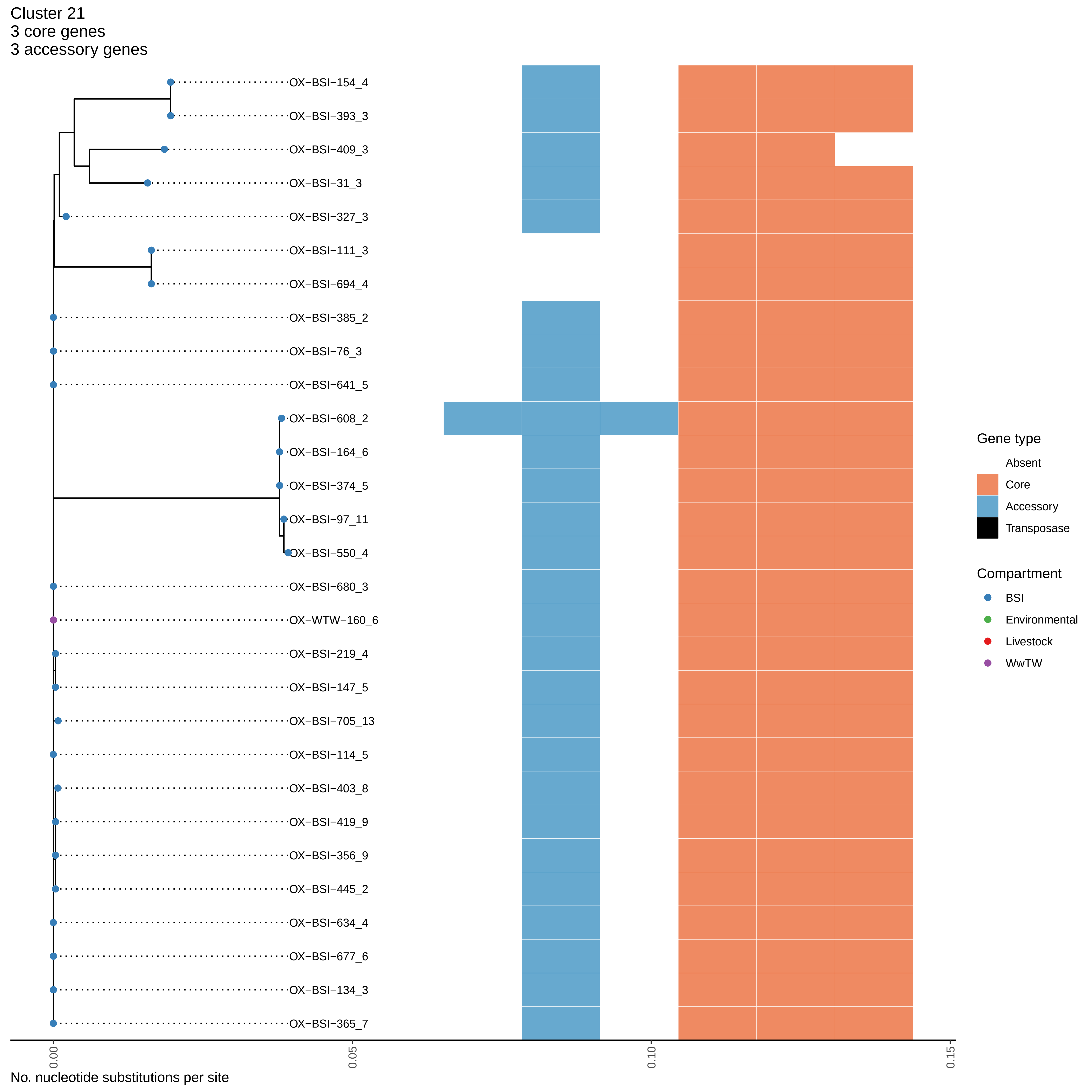

Cluster 21
3 core genes
3 accessory genes
OX−BSI−154_4
OX−BSI−393_3
OX−BSI−409_3
OX−BSI−31_3
OX−BSI−327_3
OX−BSI−111_3
OX−BSI−694_4
OX−BSI−385_2
OX−BSI−76_3
OX−BSI−641_5
OX−BSI−608_2
OX−BSI−164_6
OX−BSI−374_5
OX−BSI−97_11
OX−BSI−550_4
OX−BSI−680_3
OX−WTW−160_6
OX−BSI−219_4
OX−BSI−147_5
OX−BSI−705_13
OX−BSI−114_5
OX−BSI−403_8
OX−BSI−419_9
OX−BSI−356_9
OX−BSI−445_2
OX−BSI−634_4
OX−BSI−677_6
OX−BSI−134_3
OX−BSI−365_7
Gene type
Absent
Core
Accessory
Transposase
Compartment
BSI
Environmental
Livestock
WwTW
0.00
0.05
0.10
0.15
No. nucleotide substitutions per site

Cluster 22
2 core genes
0 accessory genes

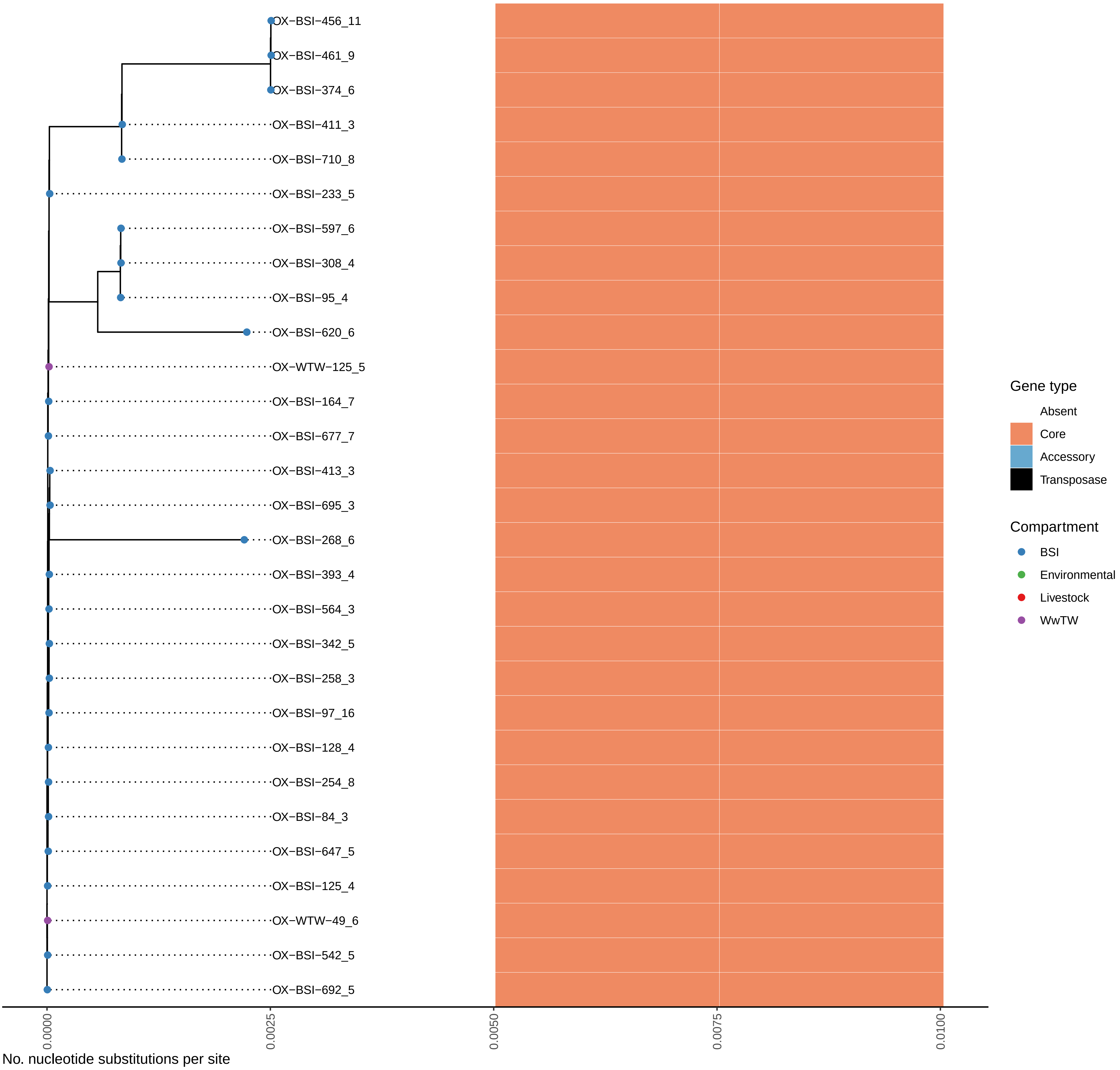

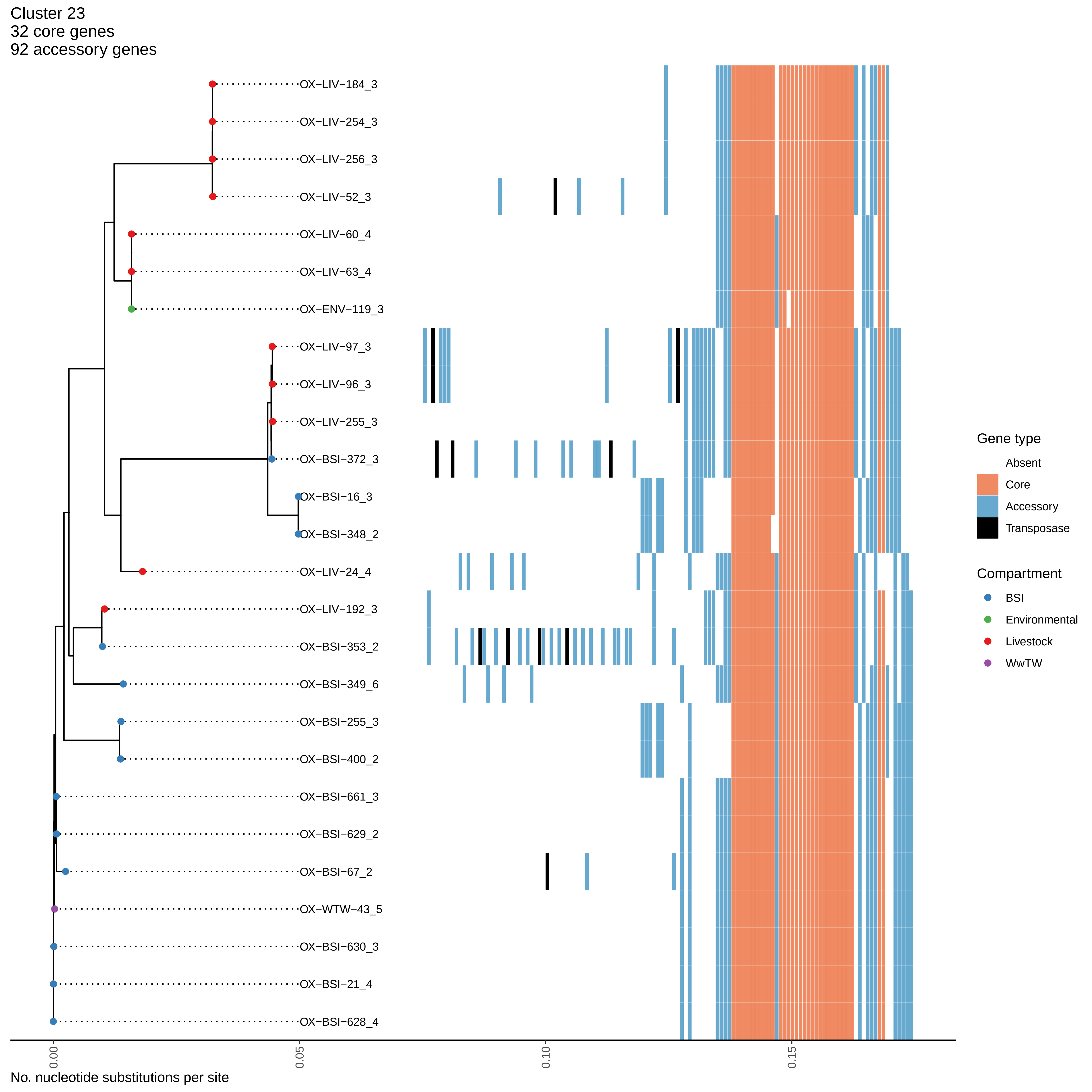

Cluster 23
32 core genes
92 accessory genes
OX−LIV−184_3
OX−LIV−254_3
OX−LIV−256_3
OX−LIV−52_3
OX−LIV−60_4
OX−LIV−63_4
OX−ENV−119_3
OX−LIV−97_3
OX−LIV−96_3
OX−LIV−255_3
OX−BSI−372_3
OX−BSI−16_3
OX−BSI−348_2
OX−LIV−24_4
OX−LIV−192_3
OX−BSI−353_2
OX−BSI−349_6
OX−BSI−255_3
OX−BSI−400_2
OX−BSI−661_3
OX−BSI−629_2
OX−BSI−67_2
OX−WTW−43_5
OX−BSI−630_3
OX−BSI−21_4
OX−BSI−628_4
0.00
0.05
0.10
0.15
No. nucleotide substitutions per site
Gene type
Absent
Core
Accessory
Transposase
Compartment
BSI
Environmental
Livestock
WwTW

Cluster 24
7 core genes
3 accessory genes

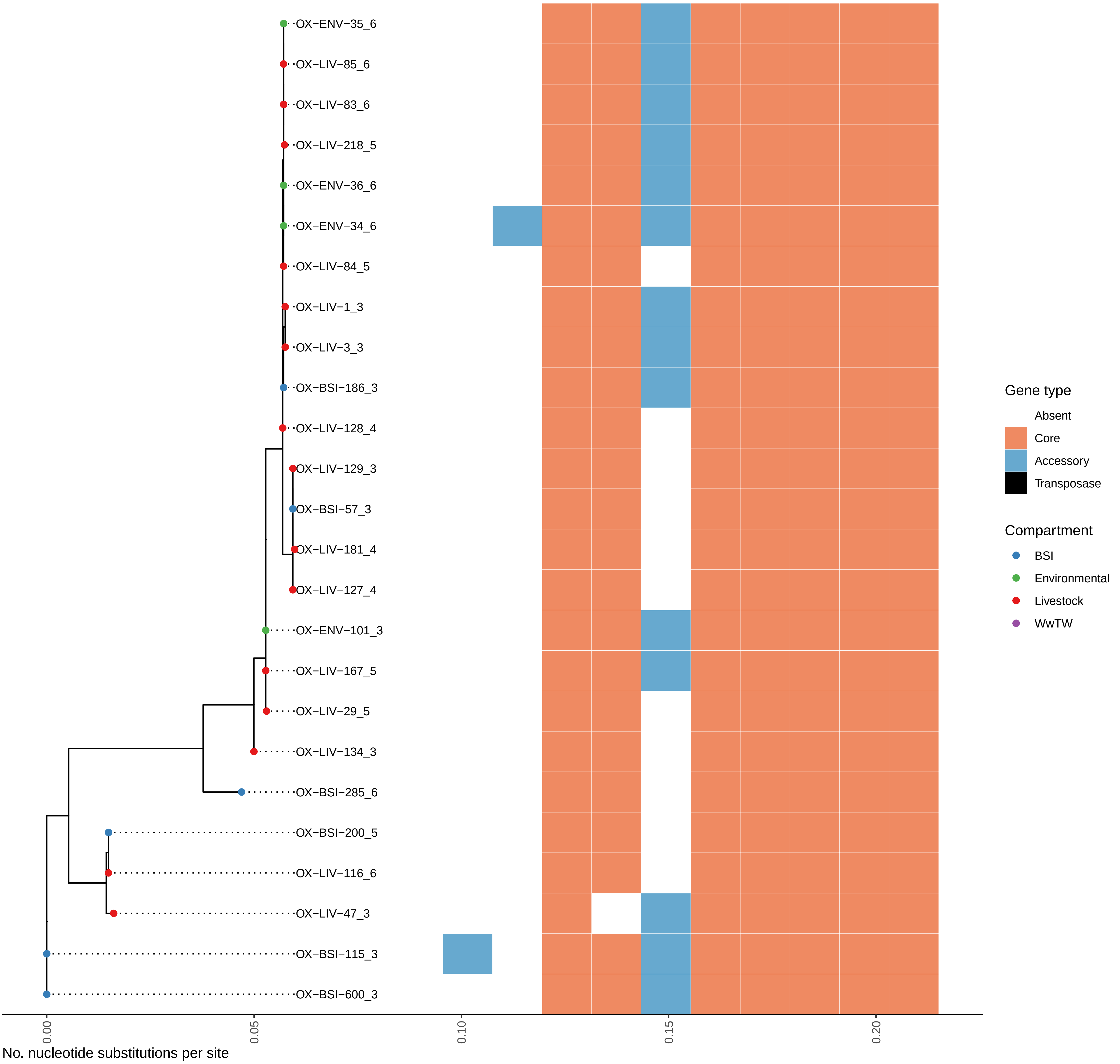

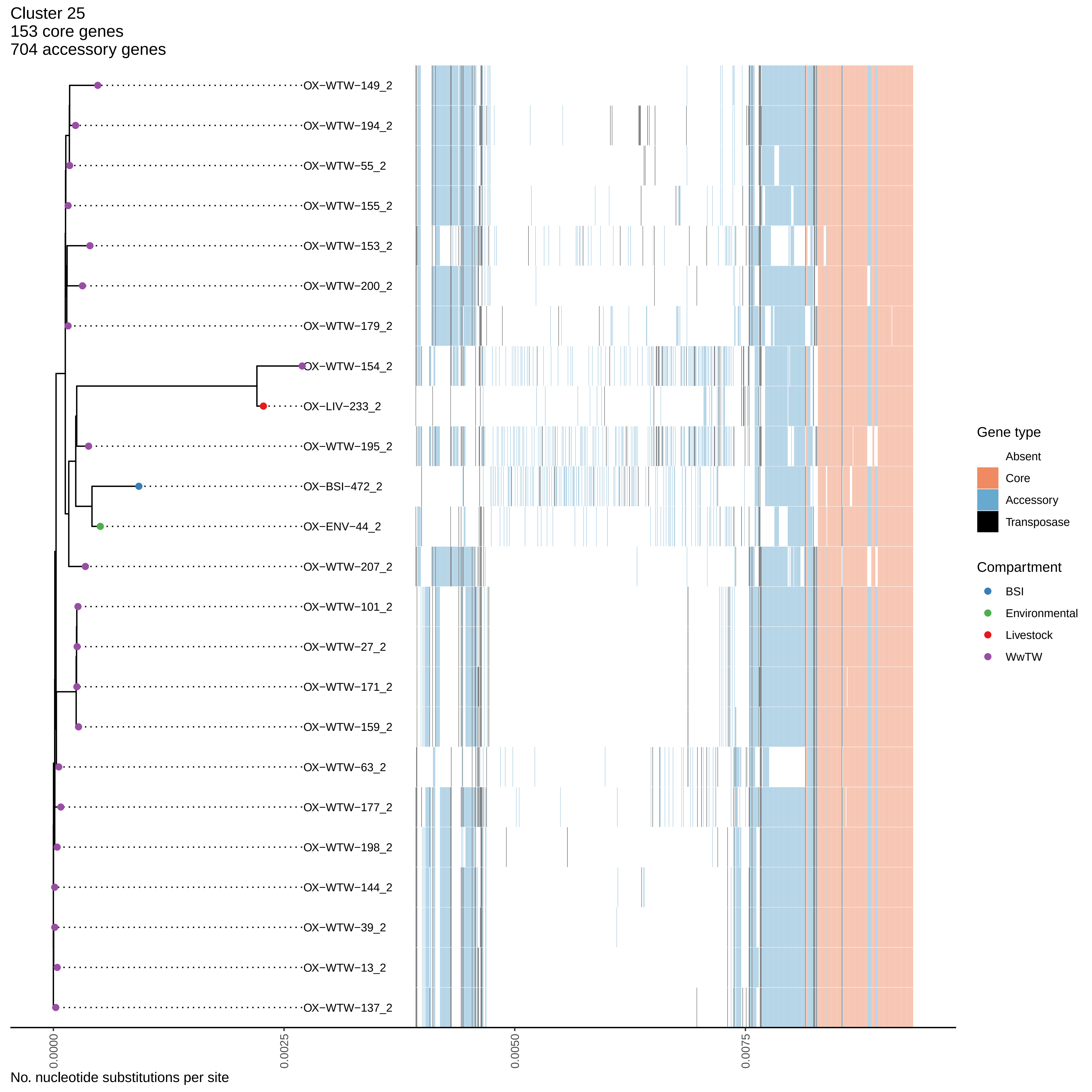
Cluster 25
153 core genes
704 accessory genes
OX−WTW−149_2
OX−WTW−194_2
OX−WTW−55_2
OX−WTW−155_2
OX−WTW−153_2
OX−WTW−200_2
OX−WTW−179_2
OX−WTW−154_2
OX−LIV−233_2
OX−WTW−195_2
OX−BSI−472_2
OX−ENV−44_2
OX−WTW−207_2
OX−WTW−101_2
OX−WTW−27_2
OX−WTW−171_2
OX−WTW−159_2
OX−WTW−63_2
OX−WTW−177_2
OX−WTW−198_2
OX−WTW−144_2
OX−WTW−39_2
OX−WTW−13_2
OX−WTW−137_2
0.0000
0.0025
0.0050
0.0075
No. nucleotide substitutions per site
Gene type
Absent
Core
Accessory
Transposase
Compartment
BSI
Environmental
Livestock
WwTW

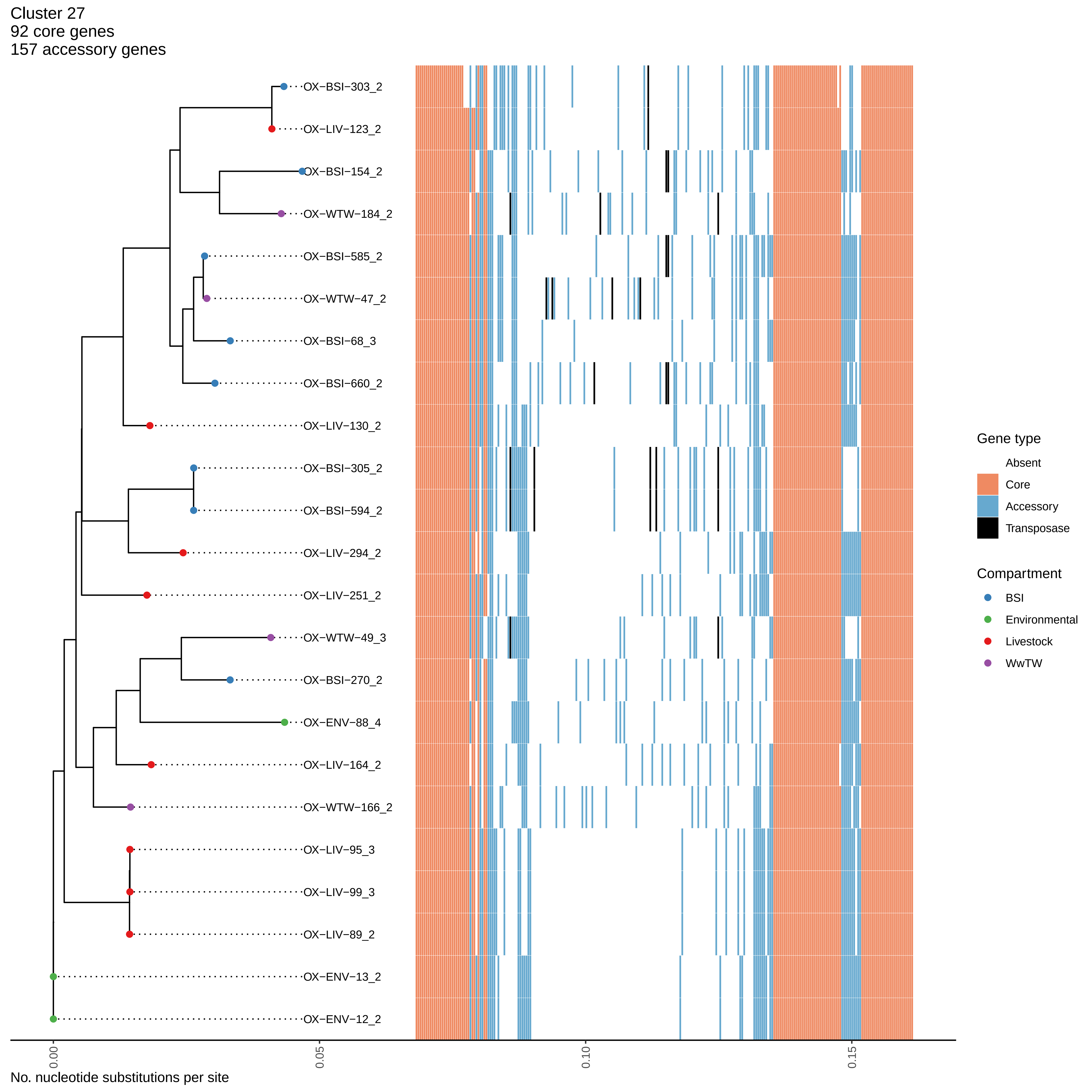

Cluster 27
92 core genes
157 accessory genes
OX−BSI−303_2
OX−LIV−123_2
OX−BSI−154_2
OX−WTW−184_2
OX−BSI−585_2
OX−WTW−47_2
OX−BSI−68_3
OX−BSI−660_2
OX−LIV−130_2
OX−BSI−305_2
OX−BSI−594_2
OX−LIV−294_2
OX−LIV−251_2
OX−WTW−49_3
OX−BSI−270_2
OX−ENV−88_4
OX−LIV−164_2
OX−WTW−166_2
OX−LIV−95_3
OX−LIV−99_3
OX−LIV−89_2
OX−ENV−13_2
OX−ENV−12_2
0.00
0.05
0.10
0.15
No. nucleotide substitutions per site
Gene type
Absent
Core
Accessory
Transposase
Compartment
BSI
Environmental
Livestock
WwTW

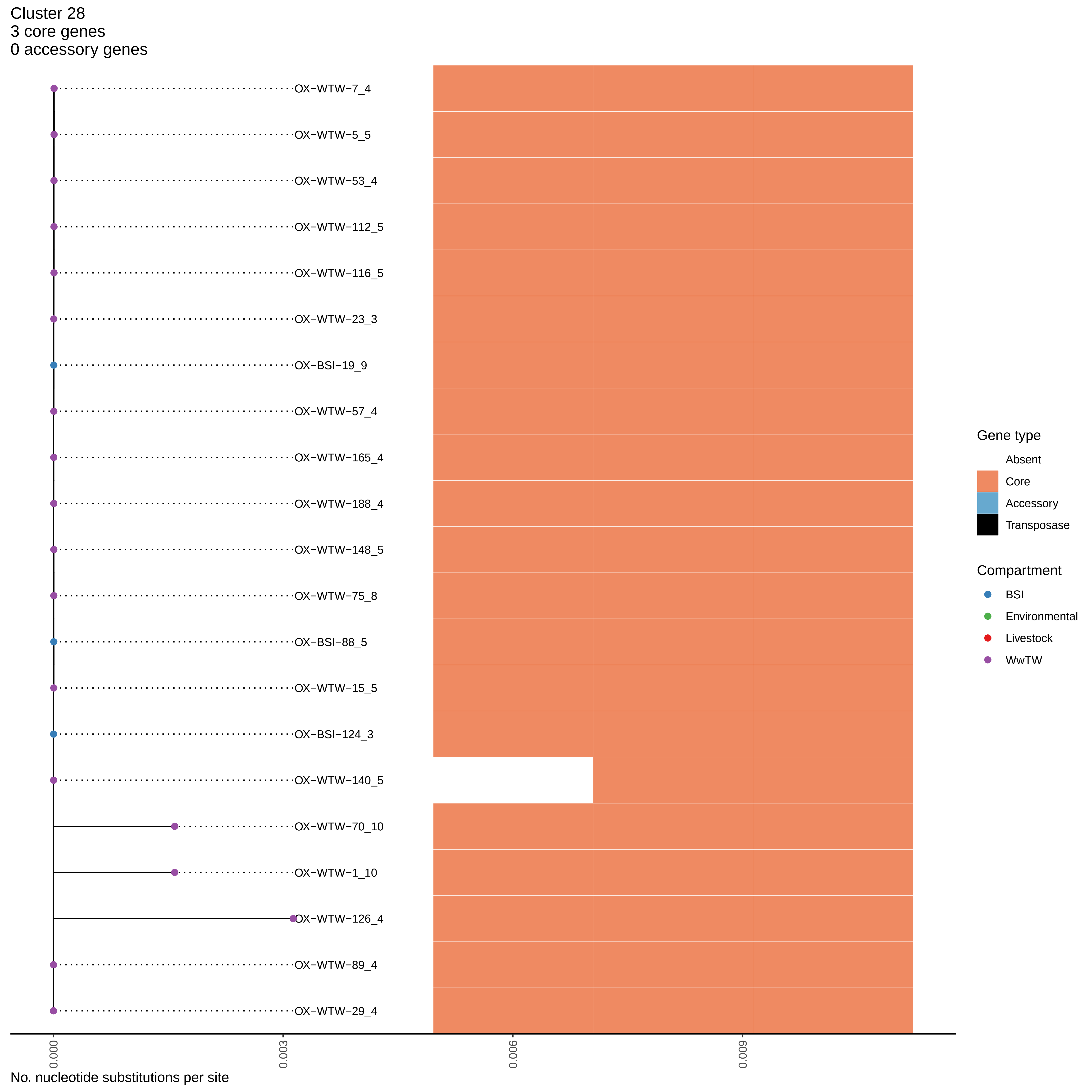

Cluster 28
3 core genes
0 accessory genes
OX−WTW−7_4
OX−WTW−5_5
OX−WTW−53_4
OX−WTW−112_5
OX−WTW−116_5
OX−WTW−23_3
OX−BSI−19_9
OX−WTW−57_4
OX−WTW−165_4
OX−WTW−188_4
OX−WTW−148_5
OX−WTW−75_8
OX−BSI−88_5
OX−WTW−15_5
OX−BSI−124_3
OX−WTW−140_5
OX−WTW−70_10
OX−WTW−1_10
OX−WTW−126_4
OX−WTW−89_4
OX−WTW−29_4
0.000
0.003
0.006
0.009
No. nucleotide substitutions per site
Gene type
Absent
Core
Accessory
Transposase
Compartment
BSI
Environmental
Livestock
WwTW

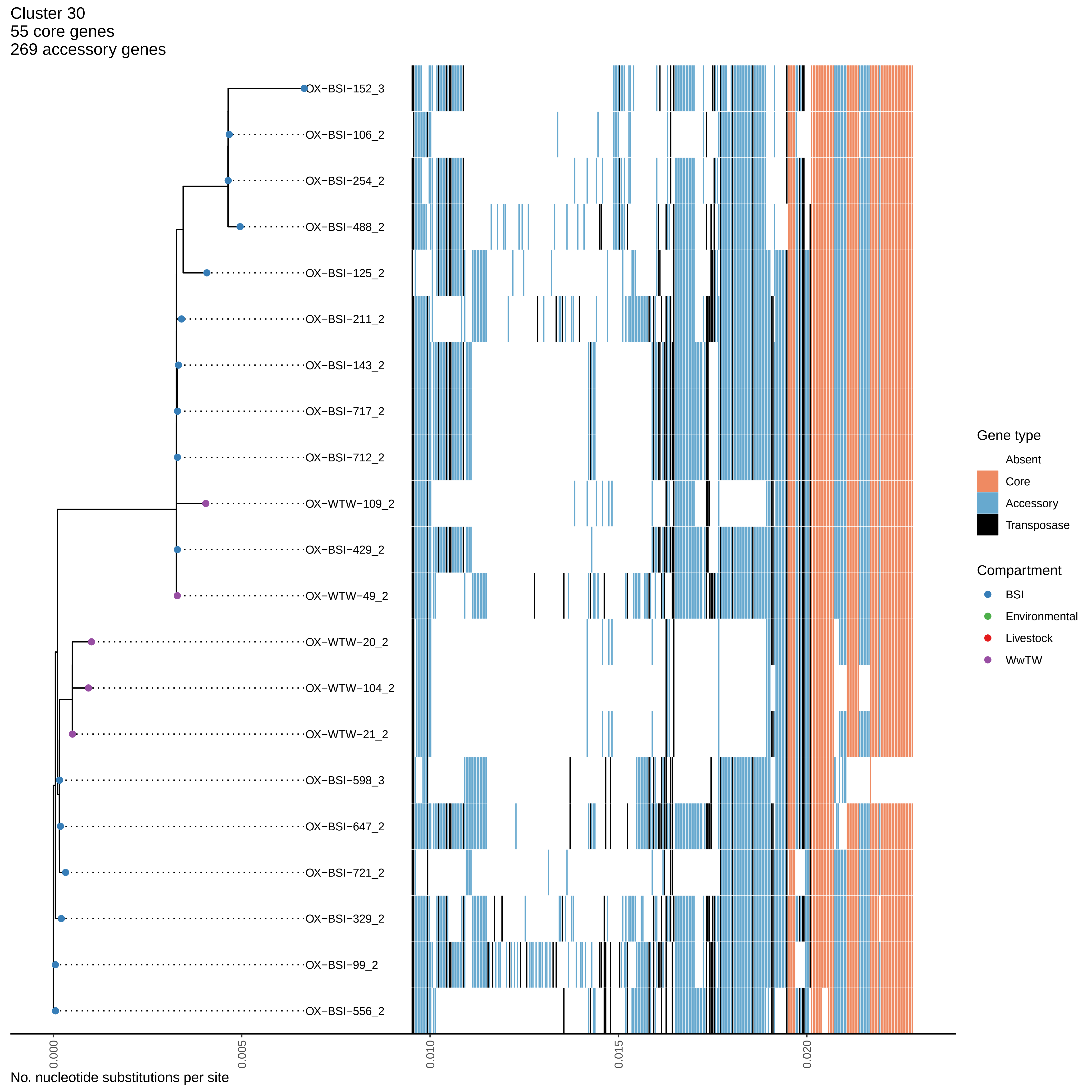

Cluster 30
55 core genes
269 accessory genes
OX−BSI−152_3
OX−BSI−106_2
OX−BSI−254_2
OX−BSI−488_2
OX−BSI−125_2
OX−BSI−211_2
OX−BSI−143_2
OX−BSI−717_2
OX−BSI−712_2
OX−WTW−109_2
OX−BSI−429_2
OX−WTW−49_2
OX−WTW−20_2
OX−WTW−104_2
OX−WTW−21_2
OX−BSI−598_3
OX−BSI−647_2
OX−BSI−721_2
OX−BSI−329_2
OX−BSI−99_2
OX−BSI−556_2
0.000
0.005
0.010
0.015
0.020
No. nucleotide substitutions per site
Gene type
Absent
Core
Accessory
Transposase
Compartment
BSI
Environmental
Livestock
WwTW

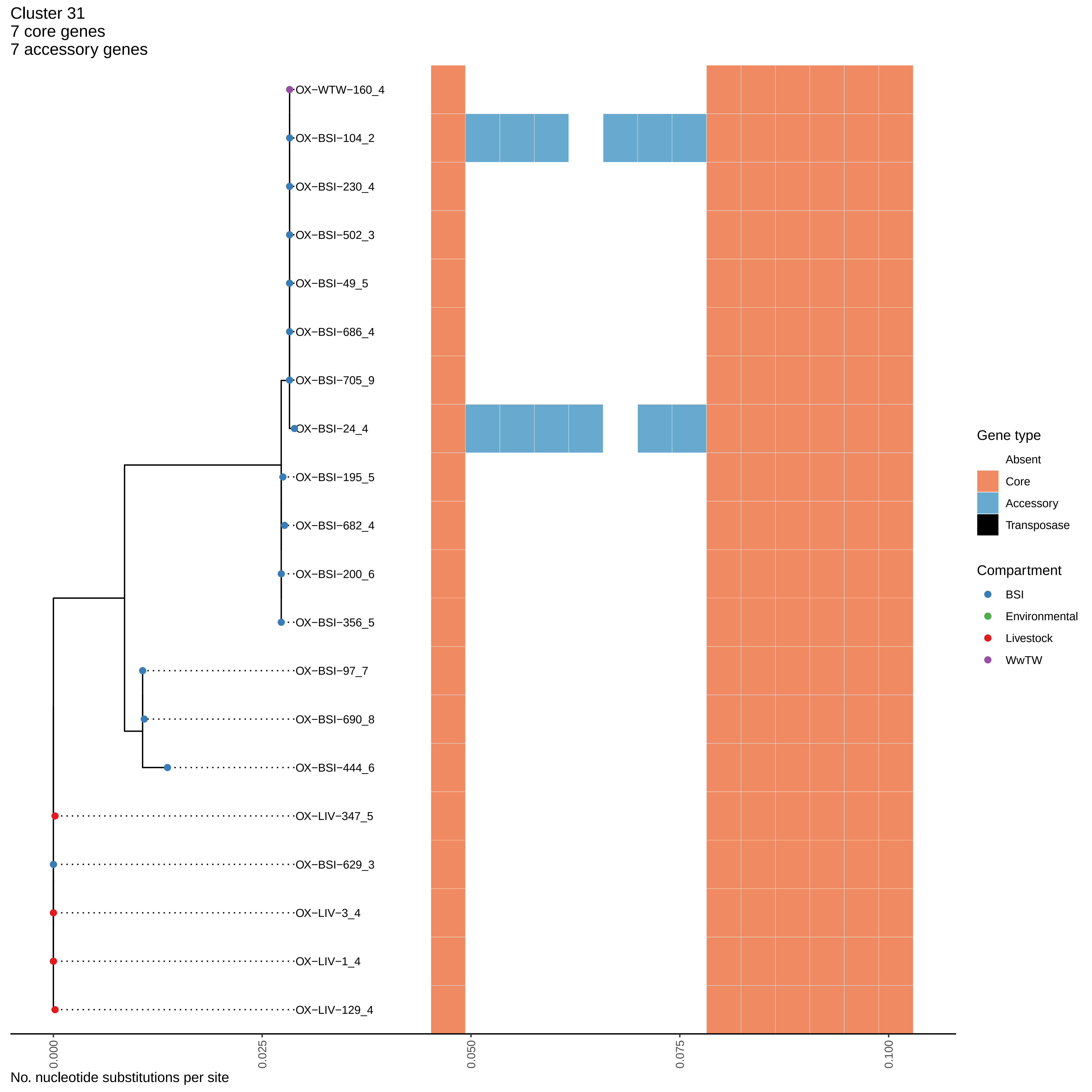
Cluster 31
7 core genes
7 accessory genes
OX−WTW−160_4
OX−BSI−104_2
OX−BSI−230_4
OX−BSI−502_3
OX−BSI−49_5
OX−BSI−686_4
OX−BSI−705_9
OX−BSI−24_4
OX−BSI−195_5
OX−BSI−682_4
OX−BSI−200_6
OX−BSI−356_5
OX−BSI−97_7
OX−BSI−690_8
OX−BSI−444_6
OX−LIV−347_5
OX−BSI−629_3
OX−LIV−3_4
OX−LIV−1_4
OX−LIV−129_4
0.000
0.025
0.050
0.075
0.100
No. nucleotide substitutions per site
Gene type
Absent
Core
Accessory
Transposase
Compartment
BSI
Environmental
Livestock
WwTW

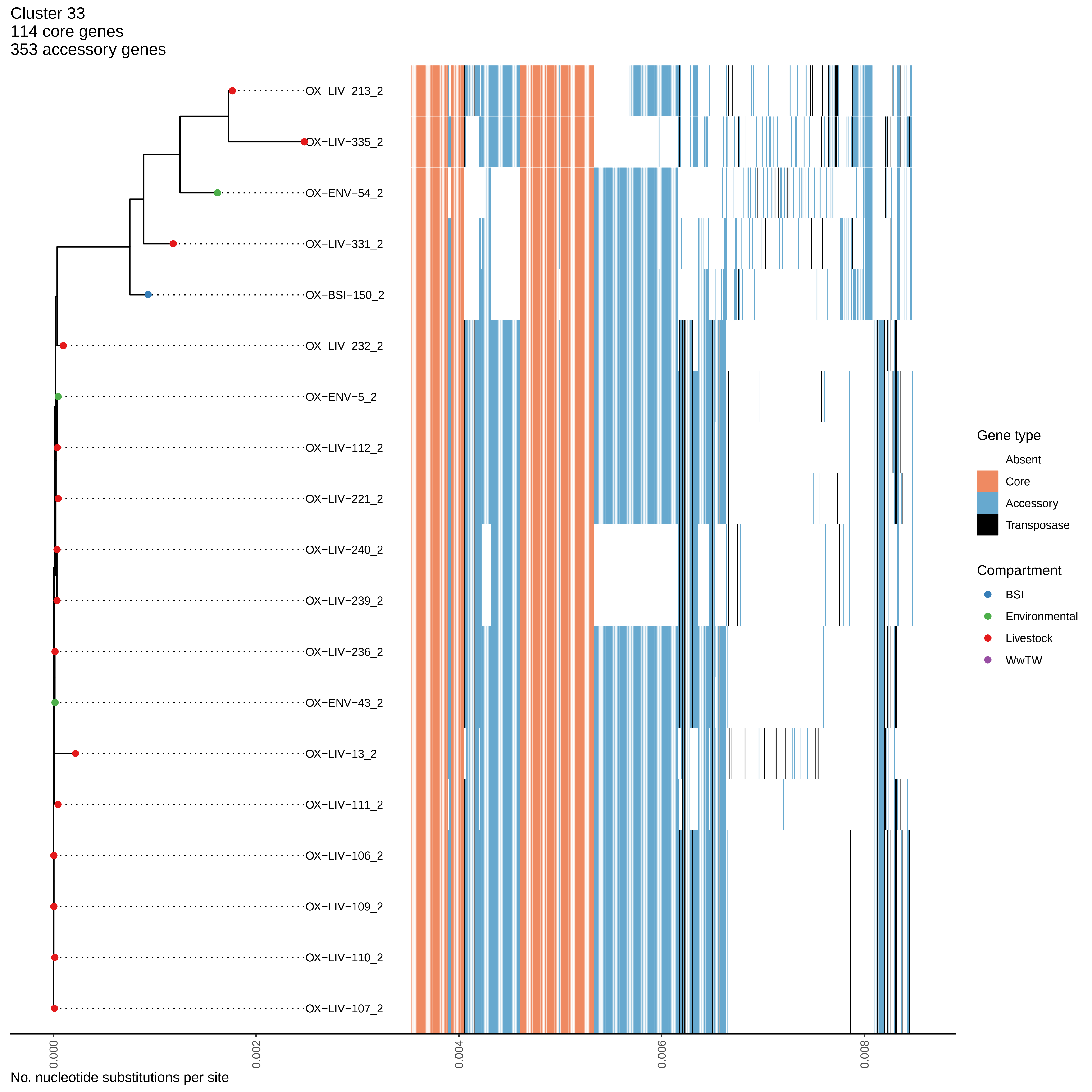

Cluster 33
114 core genes
353 accessory genes
OX−LIV−213_2
OX−LIV−335_2
OX−ENV−54_2
OX−LIV−331_2
OX−BSI−150_2
OX−LIV−232_2
OX−ENV−5_2
OX−LIV−112_2
OX−LIV−221_2
OX−LIV−240_2
OX−LIV−239_2
OX−LIV−236_2
OX−ENV−43_2
OX−LIV−13_2
OX−LIV−111_2
OX−LIV−106_2
OX−LIV−109_2
OX−LIV−110_2
OX−LIV−107_2
0.000
0.002
0.004
0.006
0.008
No. nucleotide substitutions per site
Gene type
Absent
Core
Accessory
Transposase
Compartment
BSI
Environmental
Livestock
WwTW

Cluster 34
2 core genes
3 accessory genes

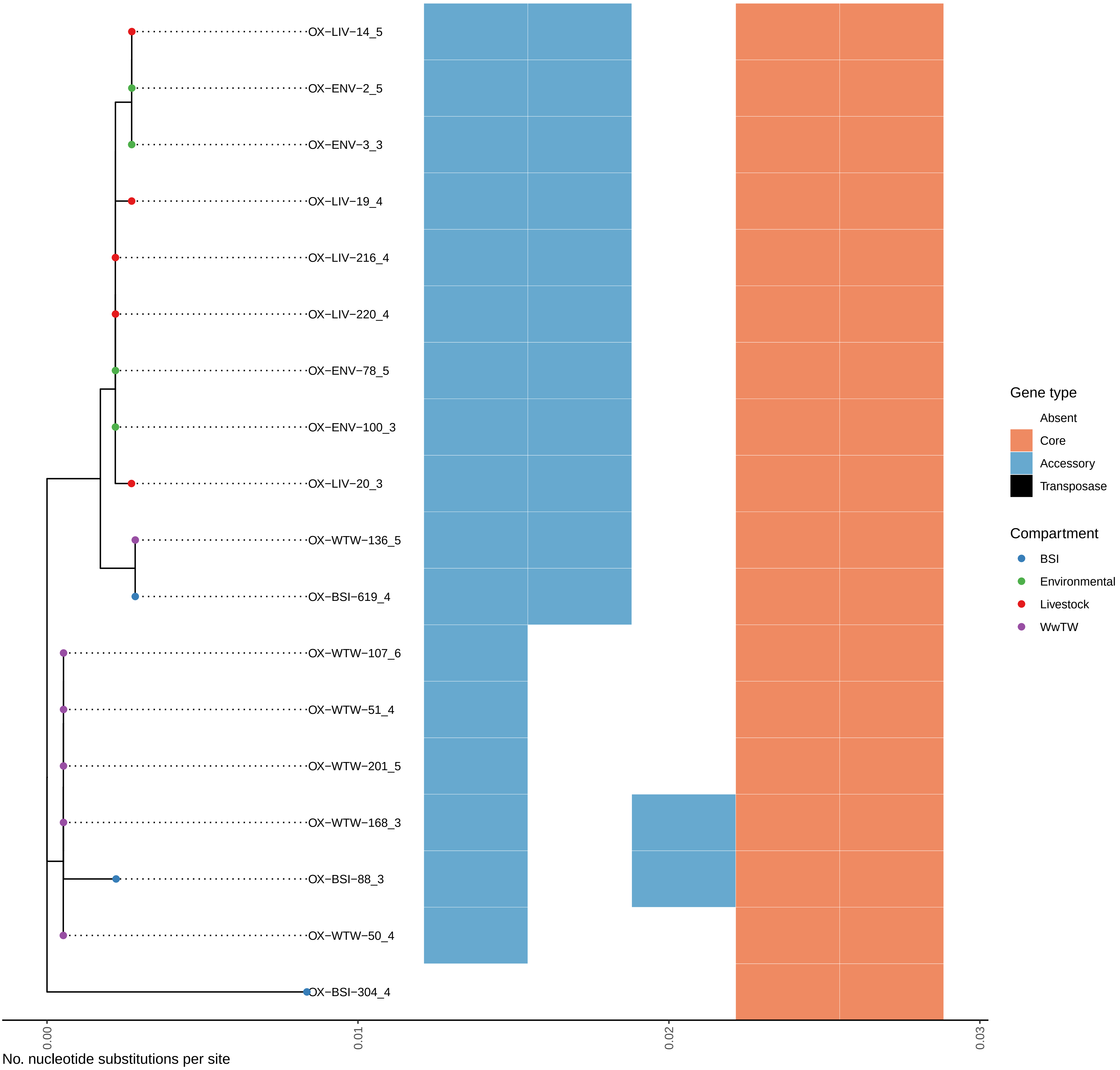

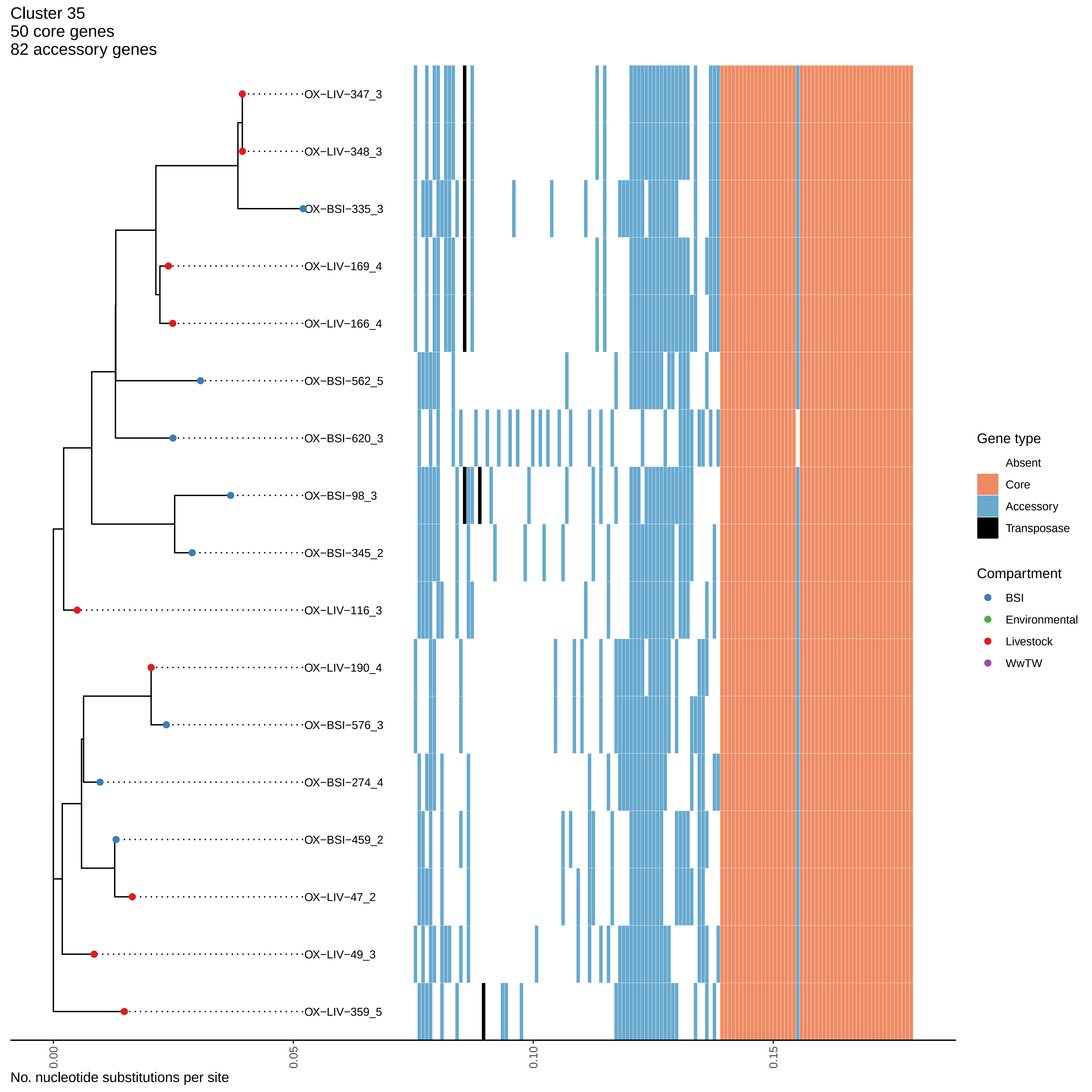

Cluster 35
50 core genes
82 accessory genes
OX−LIV−347_3
OX−LIV−348_3
OX−BSI−335_3
OX−LIV−169_4
OX−LIV−166_4
OX−BSI−562_5
OX−BSI−620_3
OX−BSI−98_3
OX−BSI−345_2
OX−LIV−116_3
OX−LIV−190_4
OX−BSI−576_3
OX−BSI−274_4
OX−BSI−459_2
OX−LIV−47_2
OX−LIV−49_3
OX−LIV−359_5
0.00
0.05
0.10
0.15
No. nucleotide substitutions per site
Gene type
Absent
Core
Accessory
Transposase
Compartment
BSI
Environmental
Livestock
WwTW

Cluster 36
103 core genes
131 accessory genes

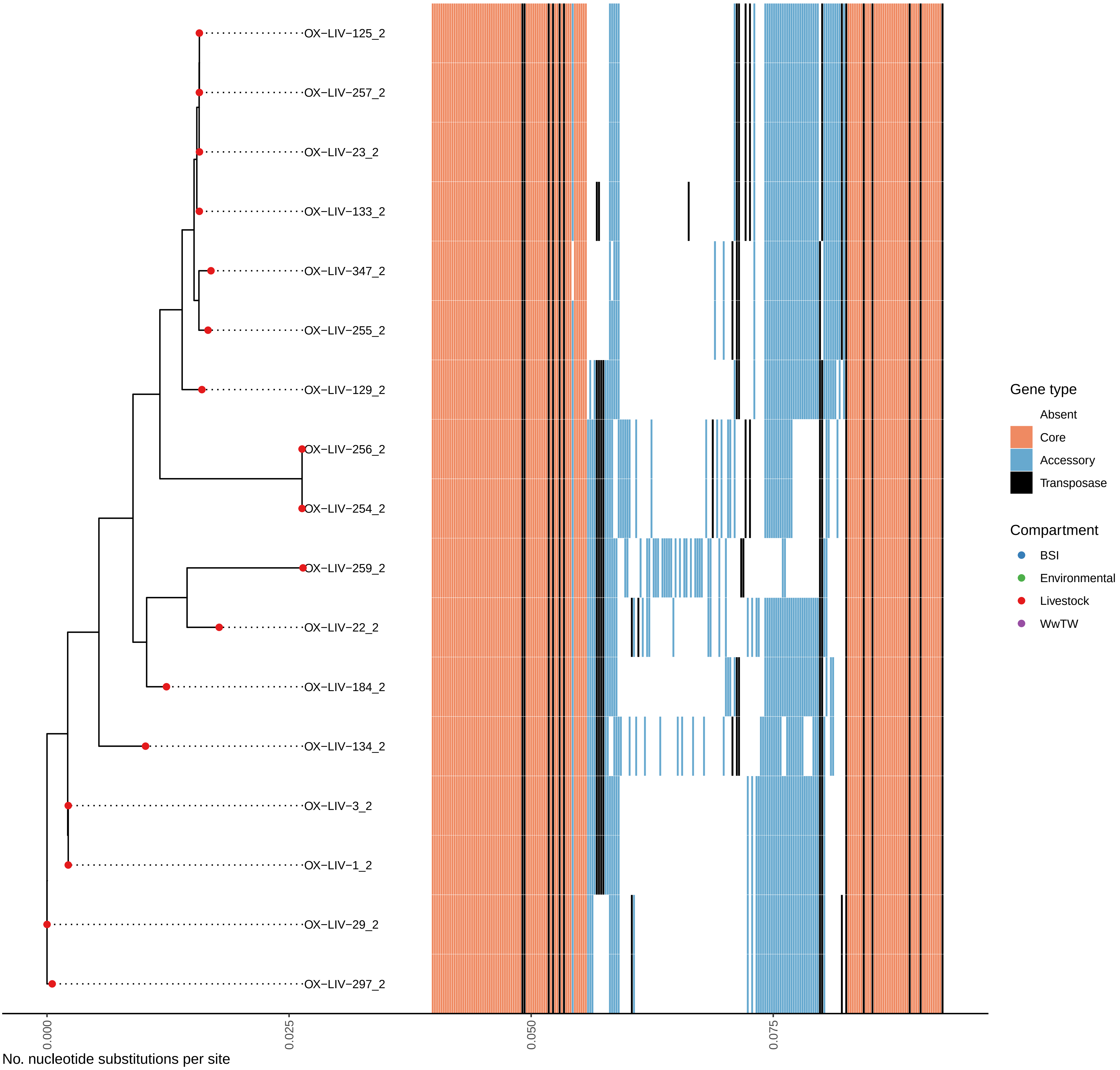

Cluster 37
5 core genes
0 accessory genes

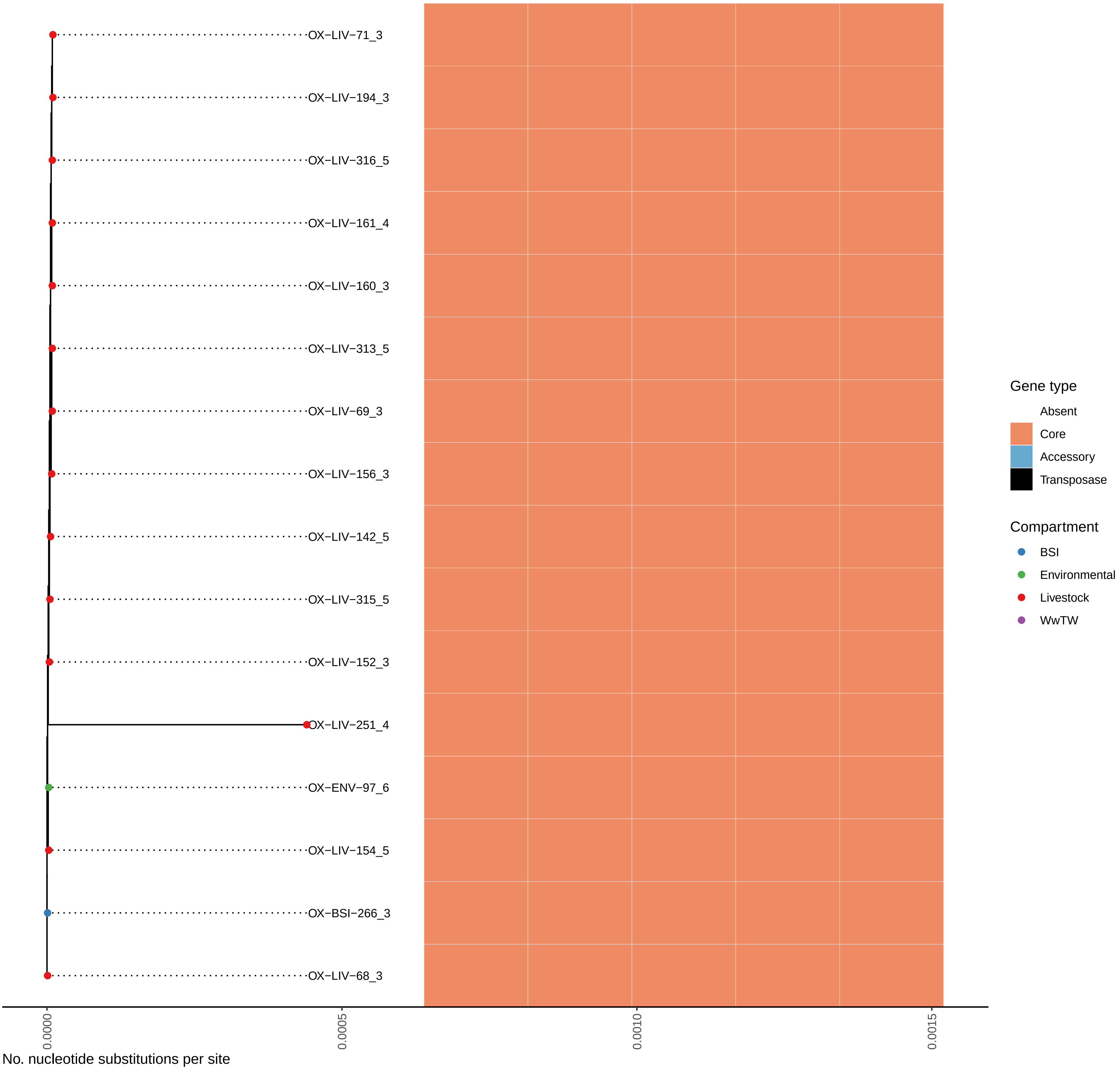

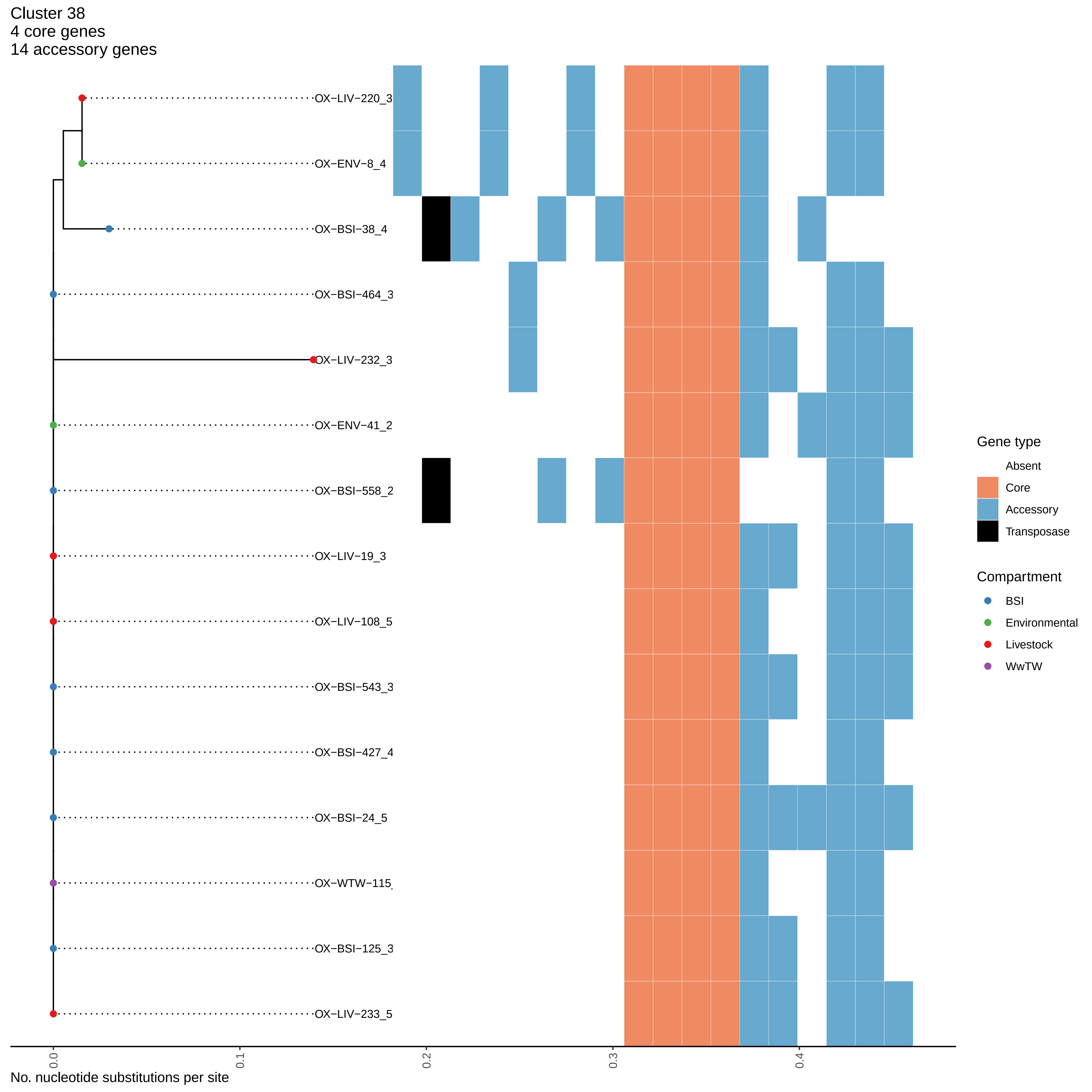

Cluster 38
4 core genes
14 accessory genes
OX−LIV−220_3
OX−ENV−8_4
OX−BSI−38_4
OX−BSI−464_3
OX−LIV−232_3
OX−ENV−41_2
OX−BSI−558_2
OX−LIV−19_3
OX−LIV−108_5
OX−BSI−543_3
OX−BSI−427_4
OX−BSI−24_5
OX−WTW−115
OX−BSI−125_3
OX−LIV−233_5
0.0
0.1
0.2
0.3
0.4
No. nucleotide substitutions per site
Gene type
Absent
Core
Accessory
Transposase
Compartment
BSI
Environmental
Livestock
WwTW

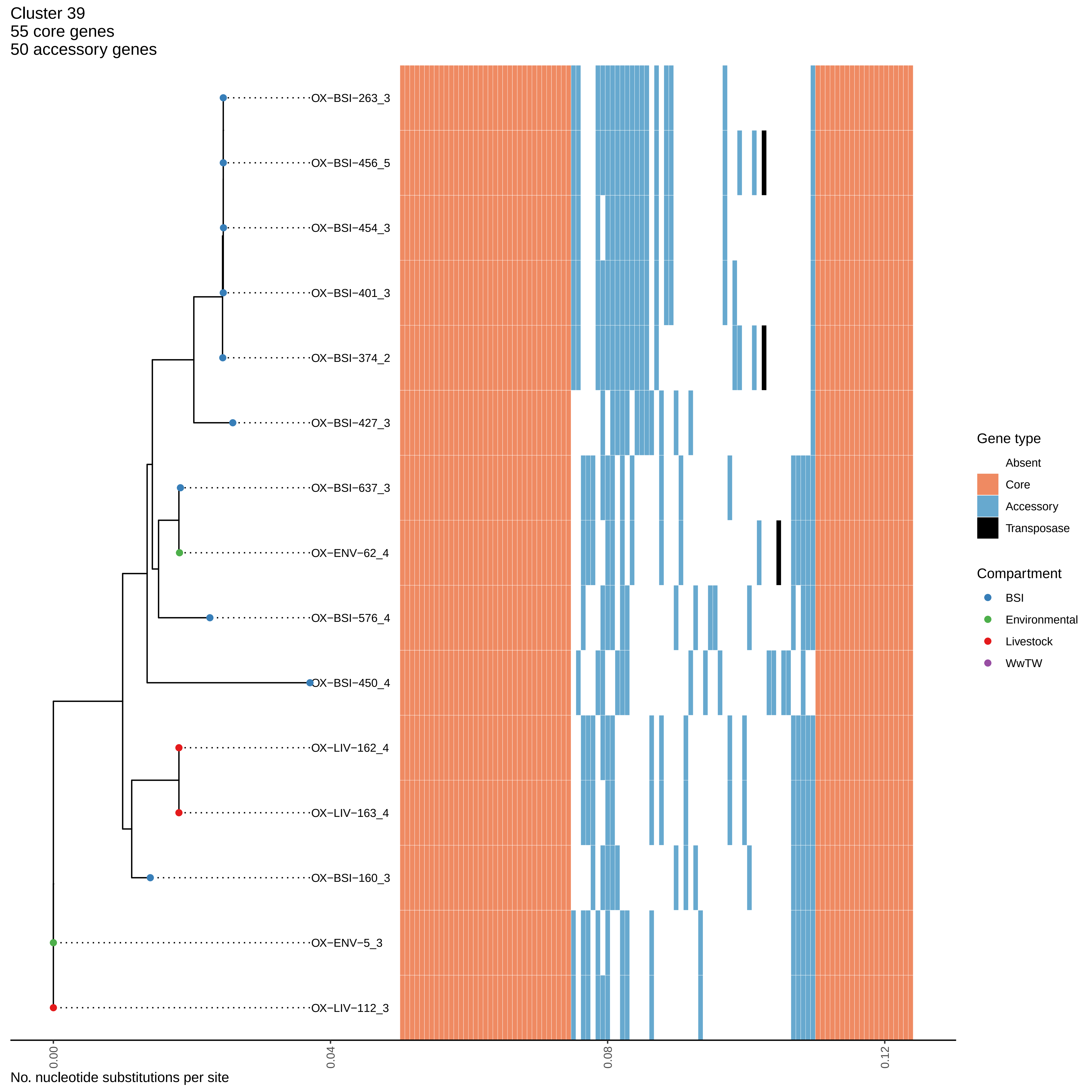

Cluster 39
55 core genes
50 accessory genes
OX−BSI−263_3
OX−BSI−456_5
OX−BSI−454_3
OX−BSI−401_3
OX−BSI−374_2
OX−BSI−427_3
OX−BSI−637_3
OX−ENV−62_4
OX−BSI−576_4
OX−BSI−450_4
OX−LIV−162_4
OX−LIV−163_4
OX−BSI−160_3
OX−ENV−5_3
OX−LIV−112_3
0.00
0.04
0.08
0.12
No. nucleotide substitutions per site
Gene type
Absent
Core
Accessory
Transposase
Compartment
BSI
Environmental
Livestock
WwTW

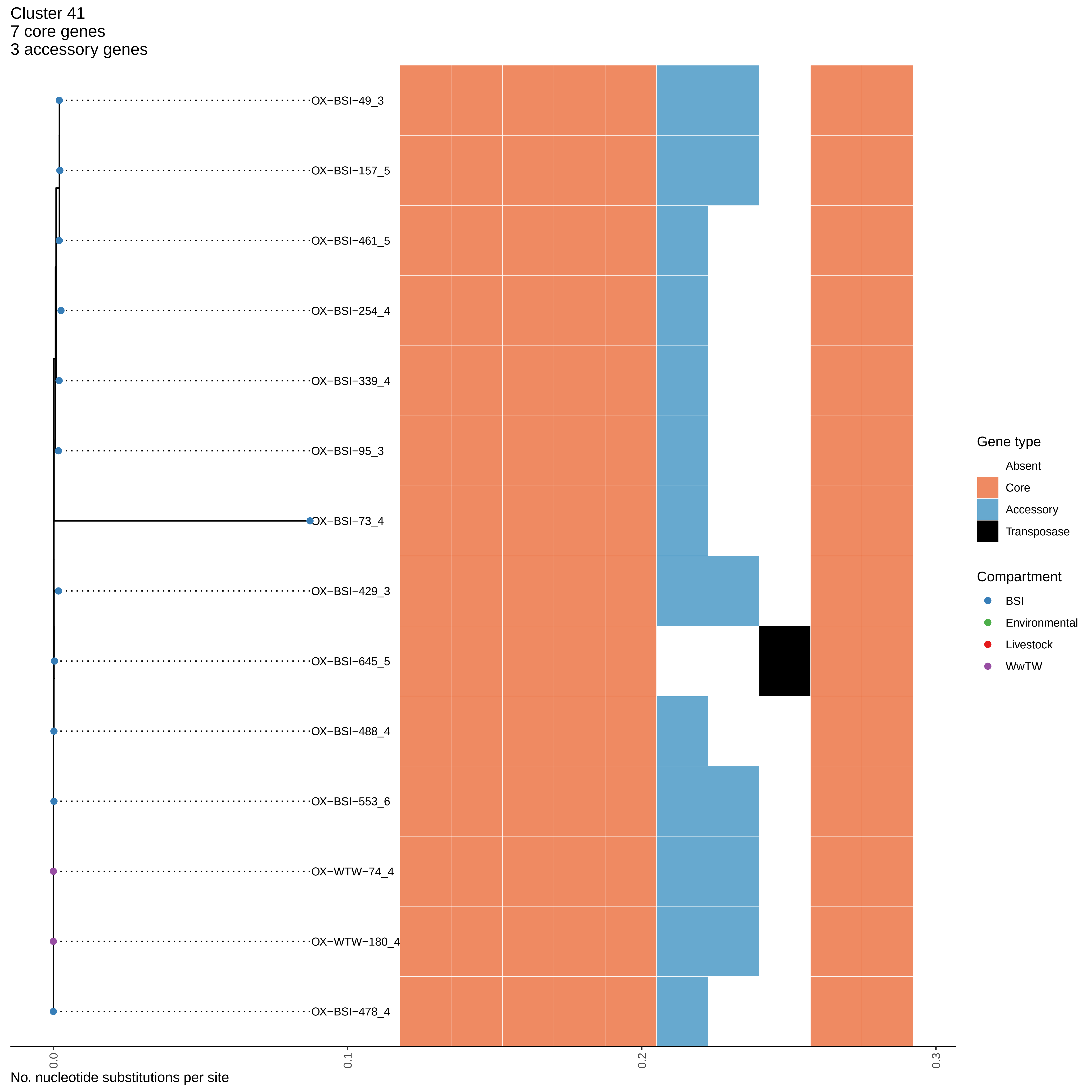

Cluster 41
7 core genes
3 accessory genes
OX−BSI−49_3
OX−BSI−157_5
OX−BSI−461_5
OX−BSI−254_4
OX−BSI−339_4
OX−BSI−95_3
OX−BSI−73_4
OX−BSI−429_3
OX−BSI−645_5
OX−BSI−488_4
OX−BSI−553_6
OX−WTW−74_4
OX−WTW−180_4
OX−BSI−478_4
Gene type
Absent
Core
Accessory
Transposase
Compartment
BSI
Environmental
Livestock
WwTW
0.0
0.1
0.2
0.3
No. nucleotide substitutions per site

Cluster 42
75 core genes
131 accessory genes

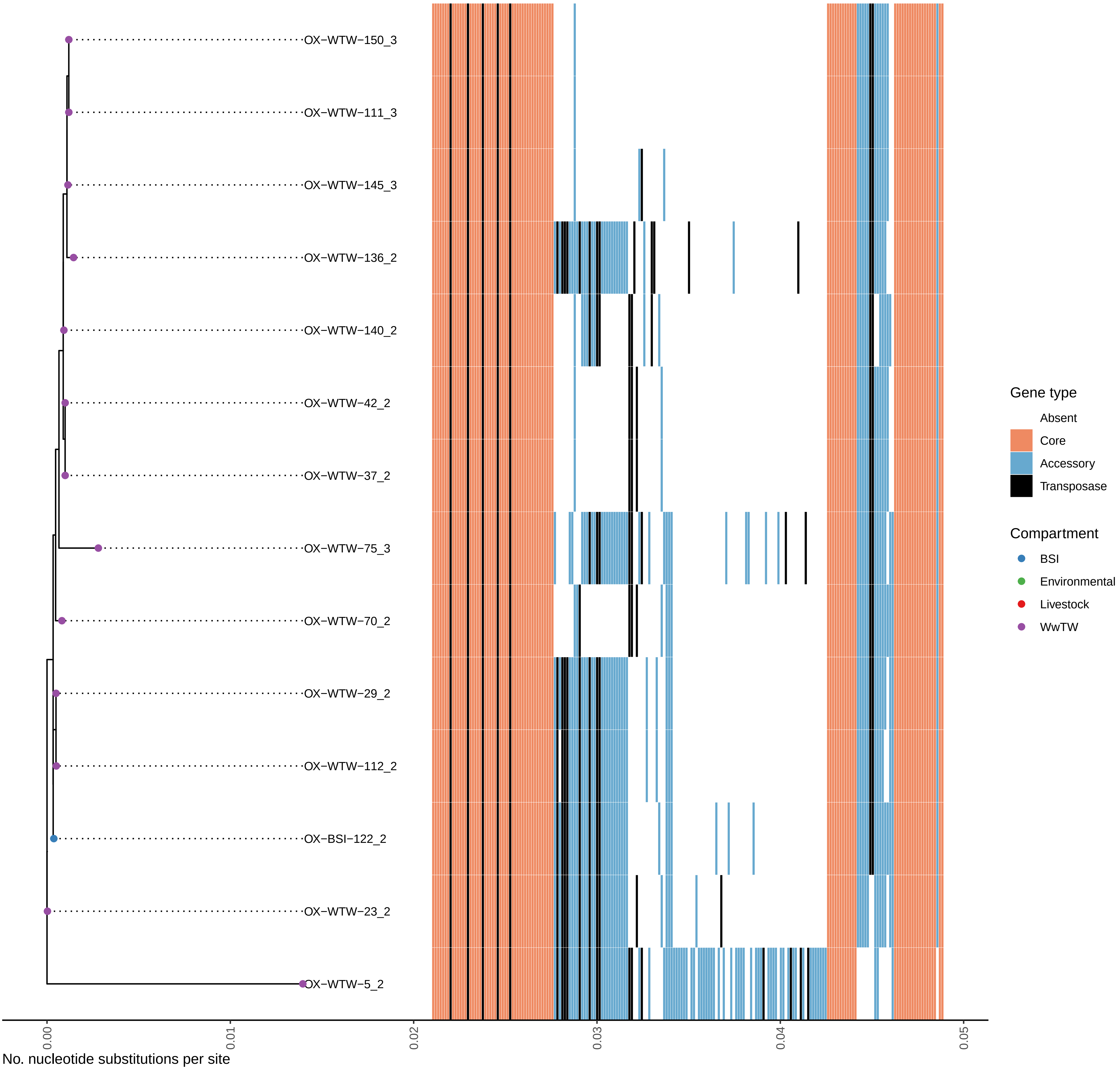

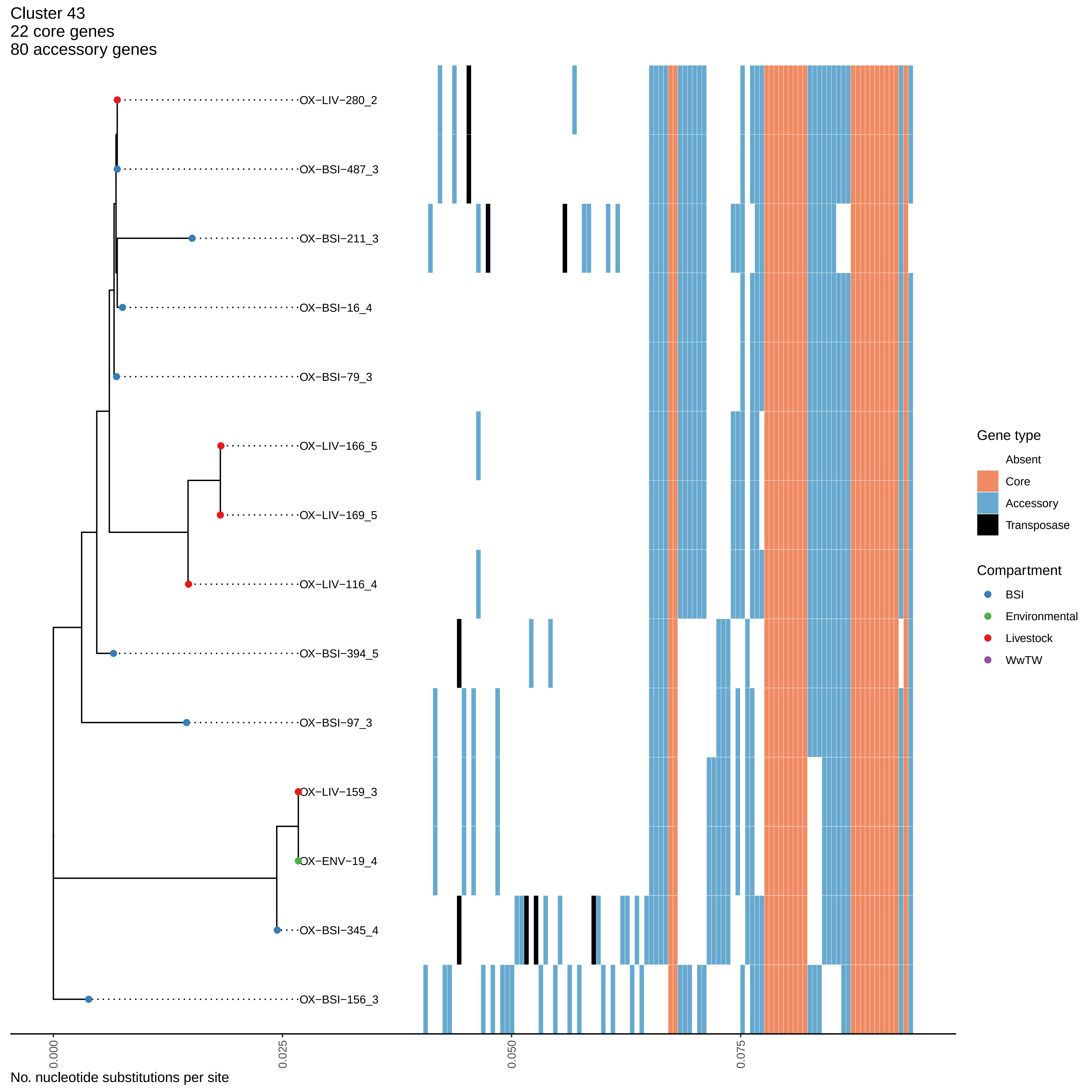
Cluster 43
22 core genes
80 accessory genes
OX−LIV−280_2
OX−BSI−487_3
OX−BSI−211_3
OX−BSI−16_4
OX−BSI−79_3
OX−LIV−166_5
OX−LIV−169_5
OX−LIV−116_4
OX−BSI−394_5
OX−BSI−97_3
OX−LIV−159_3
OX−ENV−19_4
OX−BSI−345_4
OX−BSI−156_3
Gene type
Absent
Core
Accessory
Transposase
Compartment
BSI
Environmental
Livestock
WwTW
0.000
0.025
0.050
0.075
No. nucleotide substitutions per site

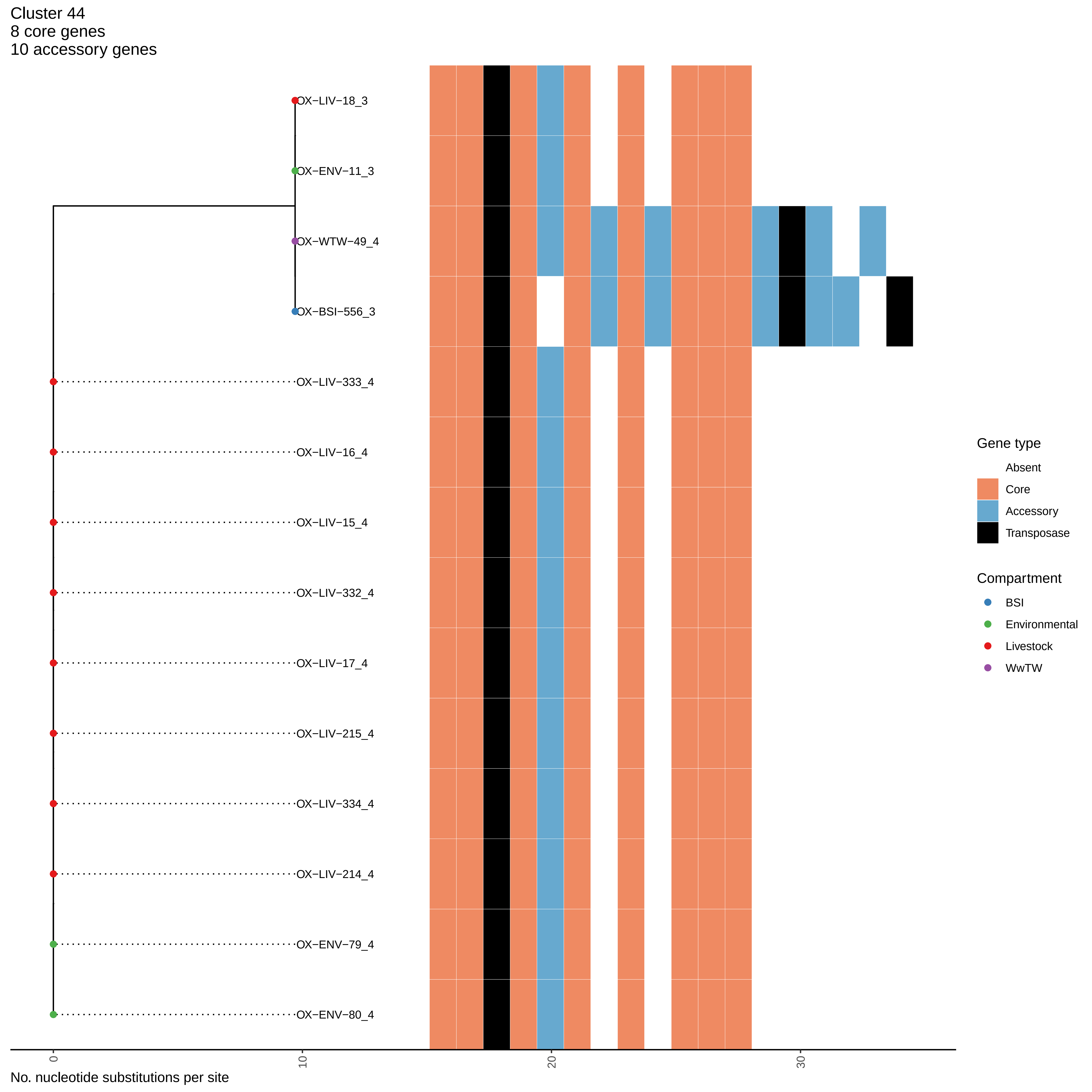
Cluster 44
8 core genes
10 accessory genes
OX−LIV−18_3
OX−ENV−11_3
OX−WTW−49_4
OX−BSI−556_3
OX−LIV−333_4
OX−LIV−16_4
OX−LIV−15_4
OX−LIV−332_4
OX−LIV−17_4
OX−LIV−215_4
OX−LIV−334_4
OX−LIV−214_4
OX−ENV−79_4
OX−ENV−80_4
0
10
20
30
No. nucleotide substitutions per site
Gene type
Absent
Core
Accessory
Transposase
Compartment
BSI
Environmental
Livestock
WwTW

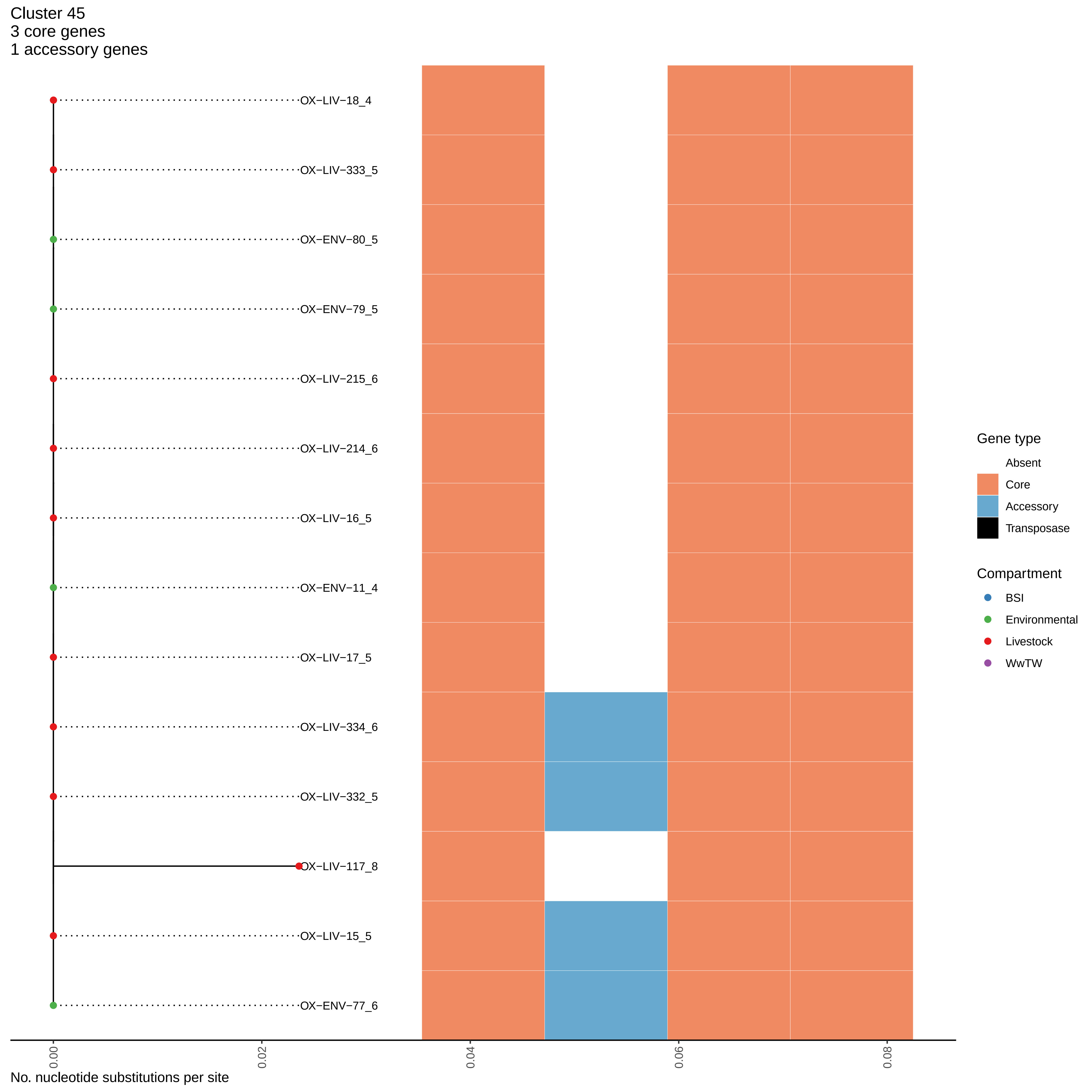

Cluster 45
3 core genes
1 accessory genes
OX−LIV−18_4
OX−LIV−333_5
OX−ENV−80_5
OX−ENV−79_5
OX−LIV−215_6
OX−LIV−214_6
OX−LIV−16_5
OX−ENV−11_4
OX−LIV−17_5
OX−LIV−334_6
OX−LIV−332_5
OX−LIV−117_8
OX−LIV−15_5
OX−ENV−77_6
0.00
0.02
0.04
0.06
0.08
No. nucleotide substitutions per site
Gene type
Absent
Core
Accessory
Transposase
Compartment
BSI
Environmental
Livestock
WwTW

Cluster 46
9 core genes
0 accessory genes

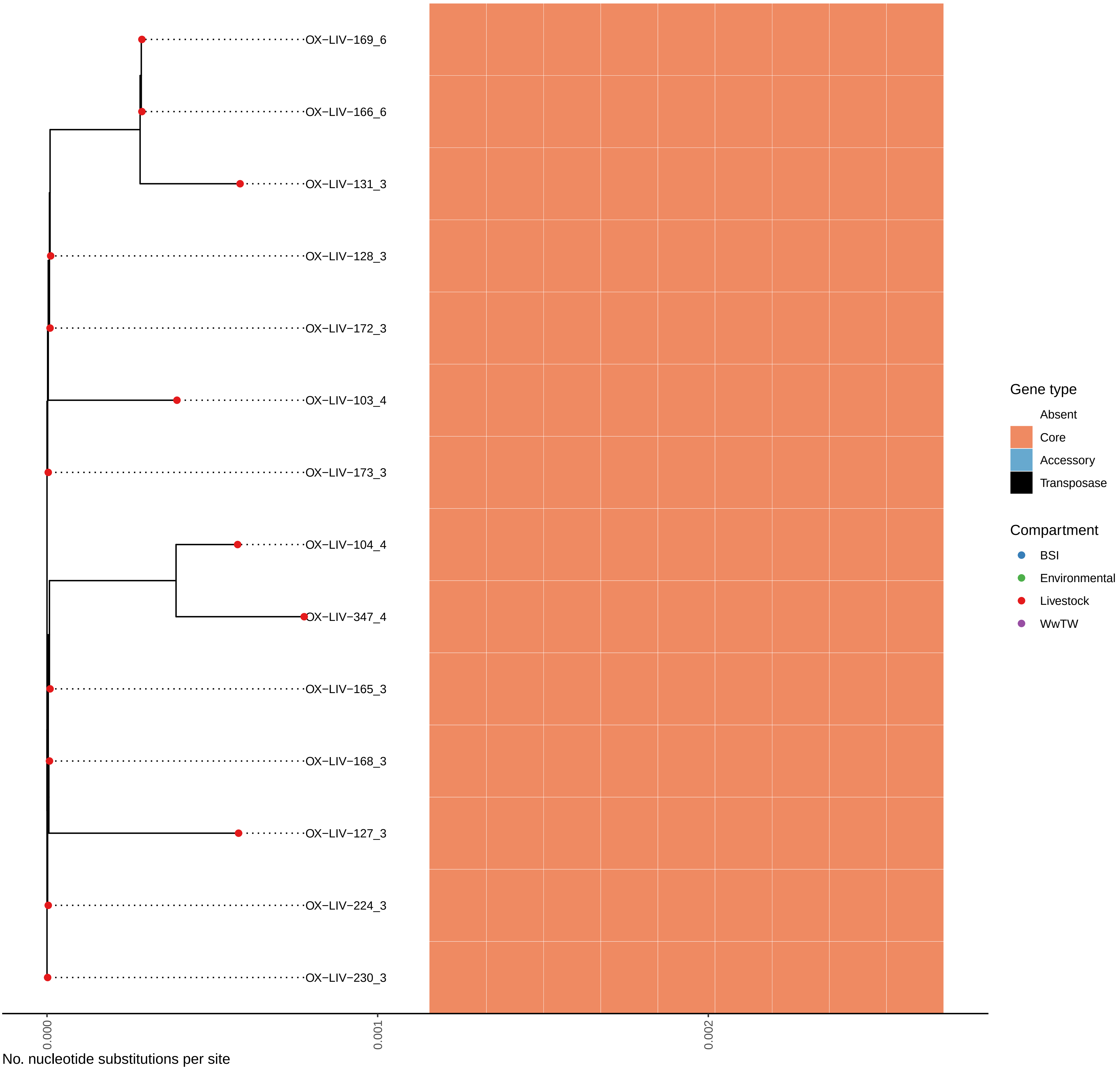

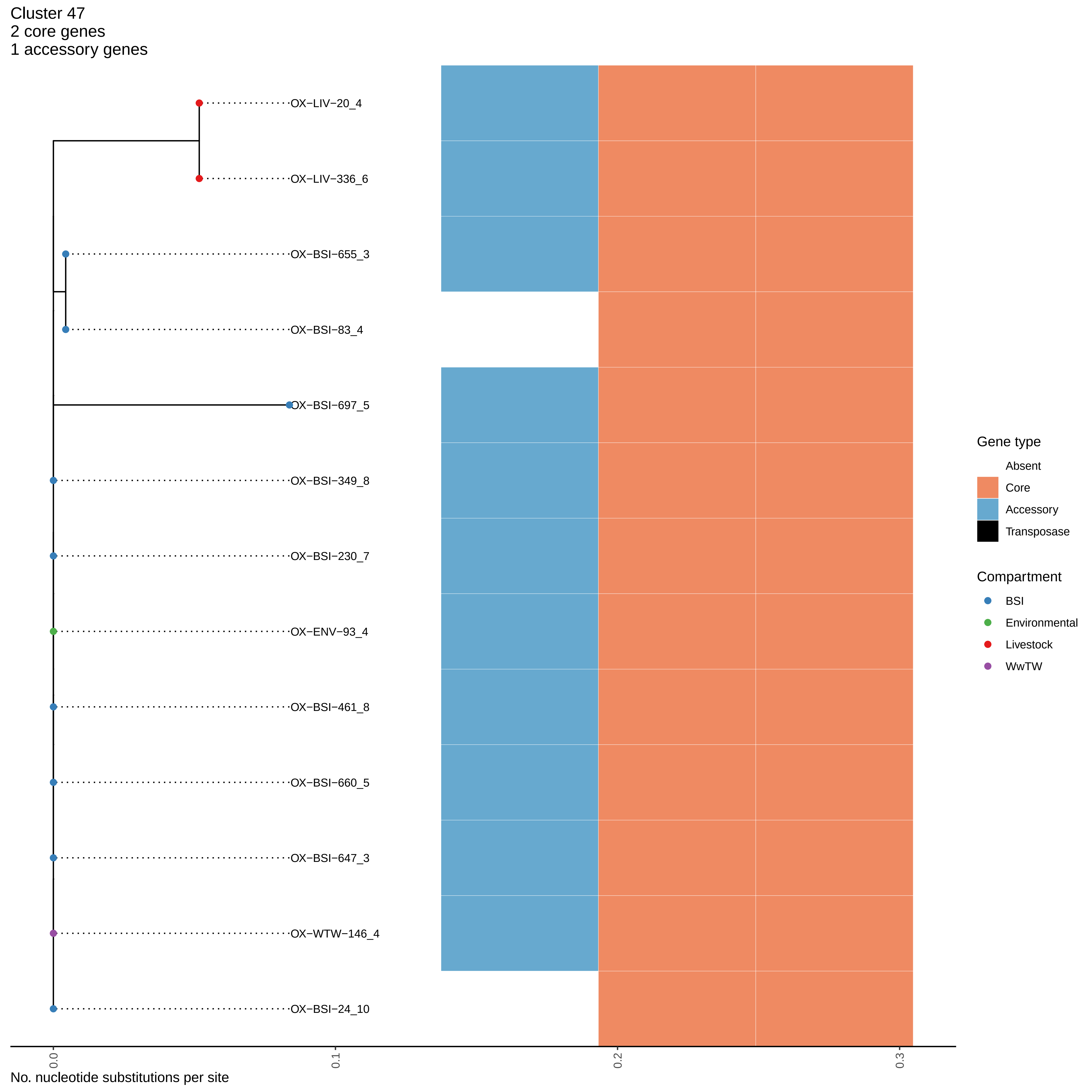

Cluster 47
2 core genes
1 accessory genes
OX−LIV−20_4
OX−LIV−336_6
OX−BSI−655_3
OX−BSI−83_4
OX−BSI−697_5
OX−BSI−349_8
OX−BSI−230_7
OX−ENV−93_4
OX−BSI−461_8
OX−BSI−660_5
OX−BSI−647_3
OX−WTW−146_4
OX−BSI−24_10
0.0
0.1
0.2
0.3
No. nucleotide substitutions per site
Gene type
Absent
Core
Accessory
Transposase
Compartment
BSI
Environmental
Livestock
WwTW

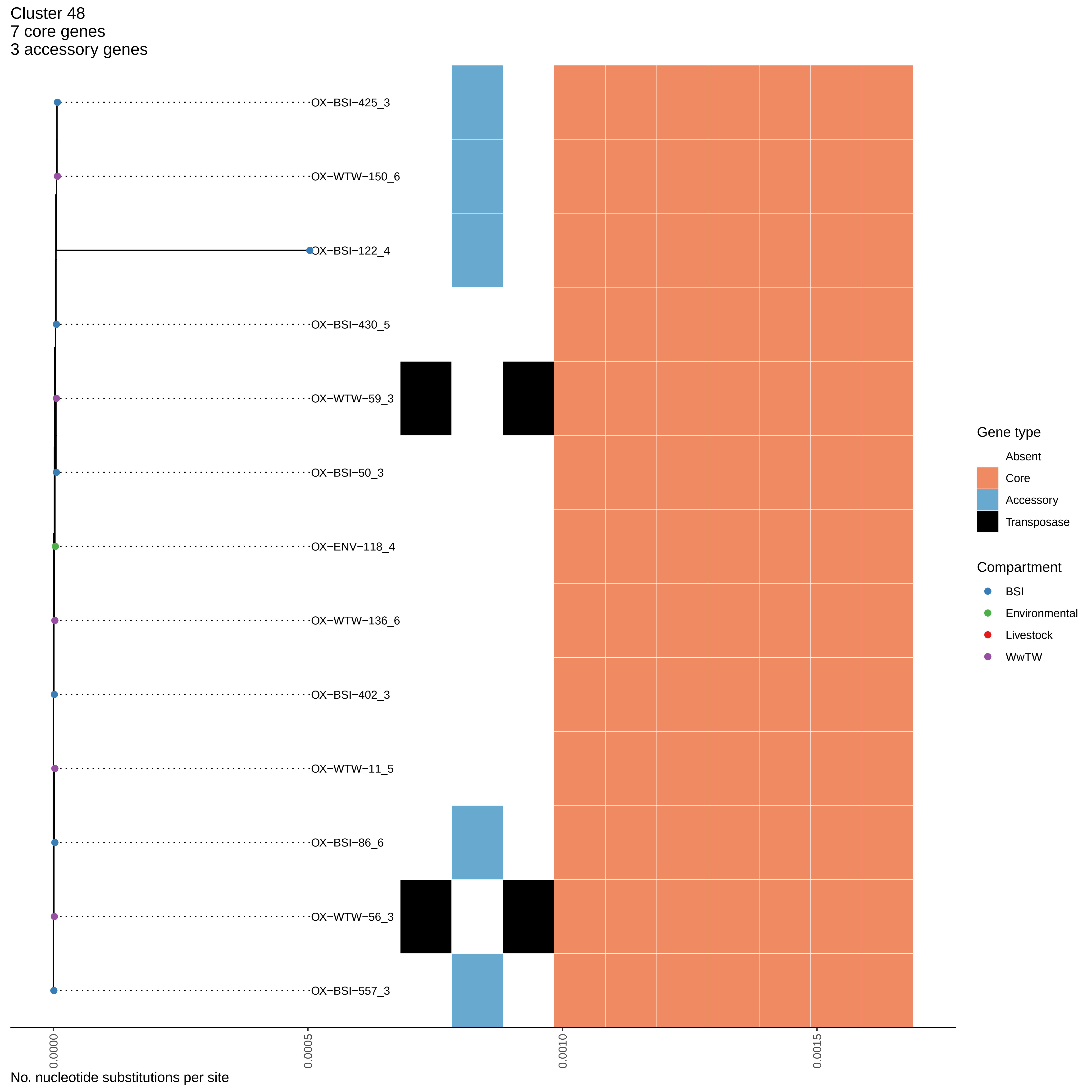
Cluster 48
7 core genes
3 accessory genes
OX−BSI−425_3
OX−WTW−150_6
OX−BSI−122_4
OX−BSI−430_5
OX−WTW−59_3
OX−BSI−50_3
OX−ENV−118_4
OX−WTW−136_6
OX−BSI−402_3
OX−WTW−11_5
OX−BSI−86_6
OX−WTW−56_3
OX−BSI−557_3
0.0000
0.0005
0.0010
0.0015
No. nucleotide substitutions per site
Gene type
Absent
Core
Accessory
Transposase
Compartment
BSI
Environmental
Livestock
WwTW

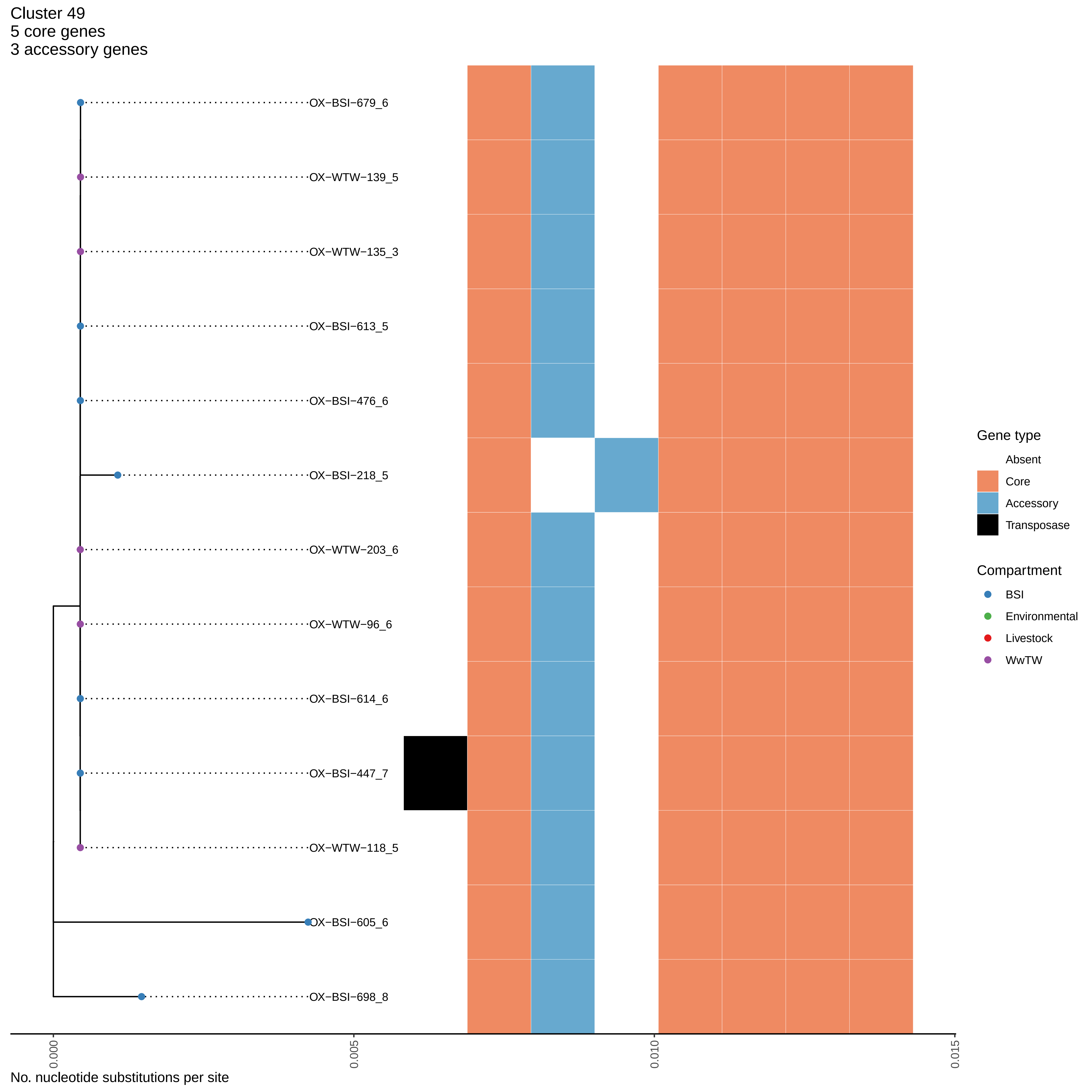
Cluster 49
5 core genes
3 accessory genes
OX−BSI−679_6
OX−WTW−139_5
OX−WTW−135_3
OX−BSI−613_5
OX−BSI−476_6
OX−BSI−218_5
OX−WTW−203_6
OX−WTW−96_6
OX−BSI−614_6
OX−BSI−447_7
OX−WTW−118_5
OX−BSI−605_6
OX−BSI−698_8
Gene type
Absent
Core
Accessory
Transposase
Compartment
BSI
Environmental
Livestock
WwTW
0.000
0.005
0.010
0.015
No. nucleotide substitutions per site

Cluster 50
38 core genes
19 accessory genes

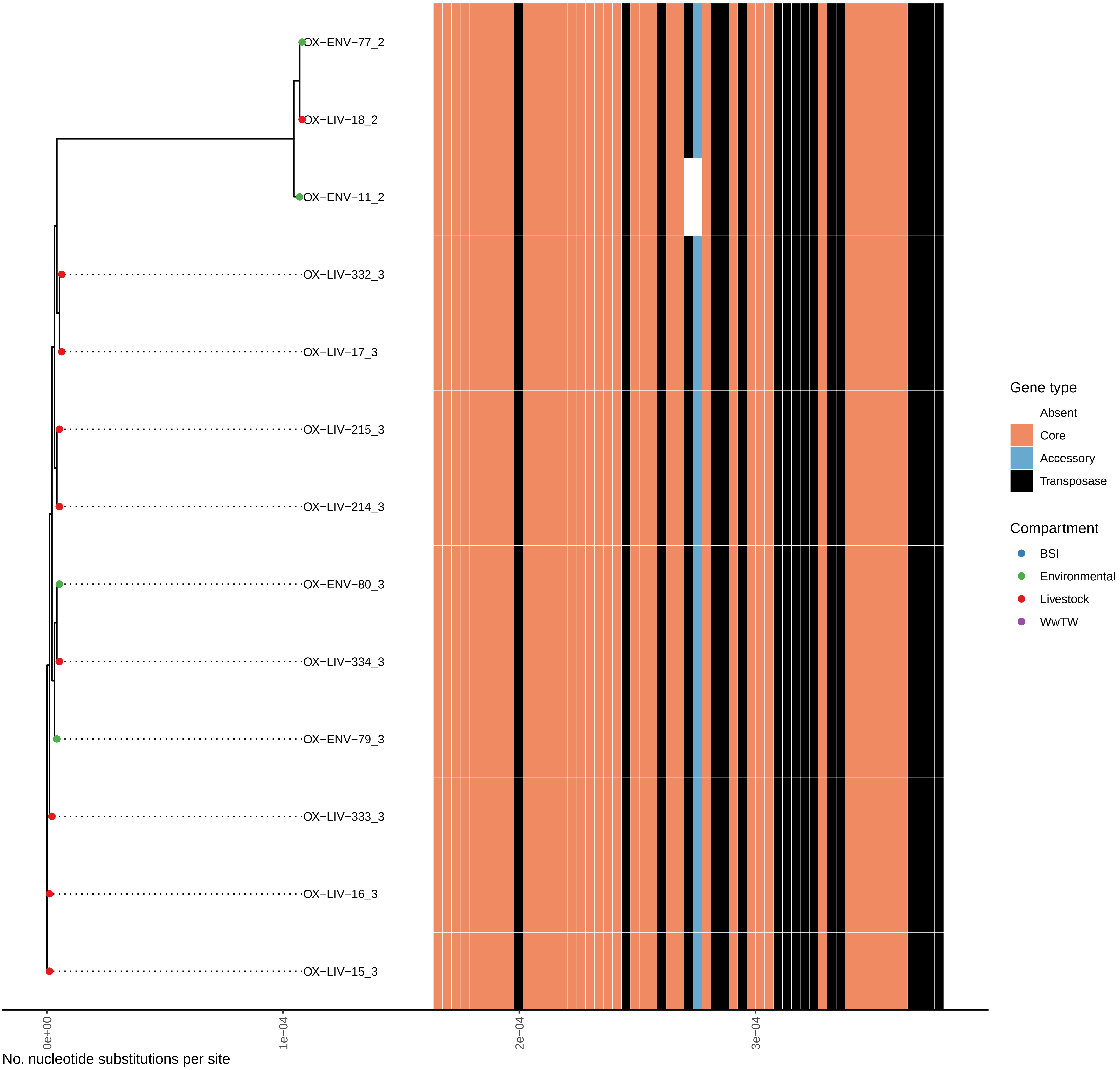

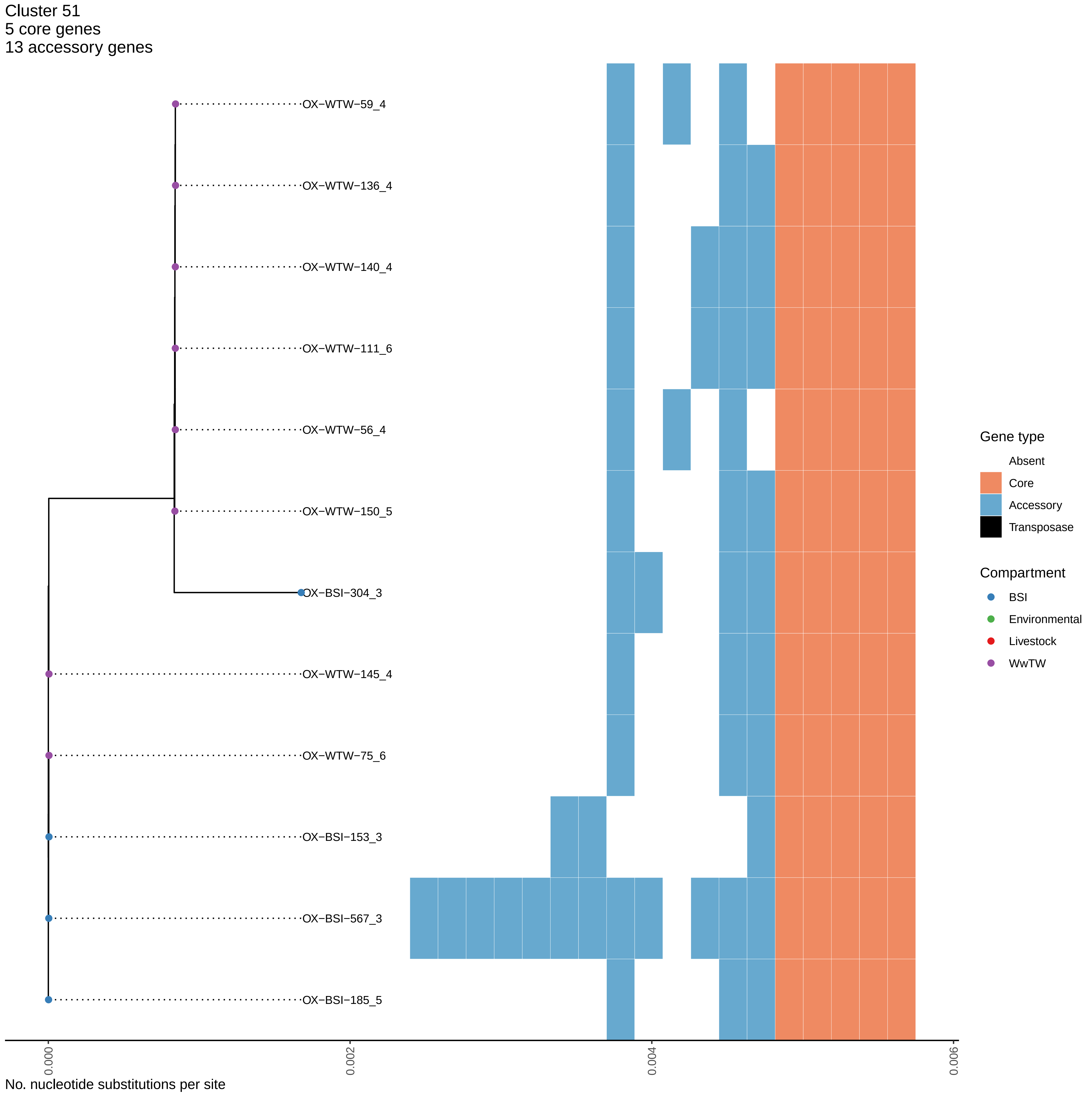

Cluster 51
5 core genes
13 accessory genes
OX−WTW−59_4
OX−WTW−136_4
OX−WTW−140_4
OX−WTW−111_6
OX−WTW−56_4
OX−WTW−150_5
OX−BSI−304_3
OX−WTW−145_4
OX−WTW−75_6
OX−BSI−153_3
OX−BSI−567_3
OX−BSI−185_5
0.000
0.002
0.004
0.006
No. nucleotide substitutions per site
Gene type
Absent
Core
Accessory
Transposase
Compartment
BSI
Environmental
Livestock
WwTW

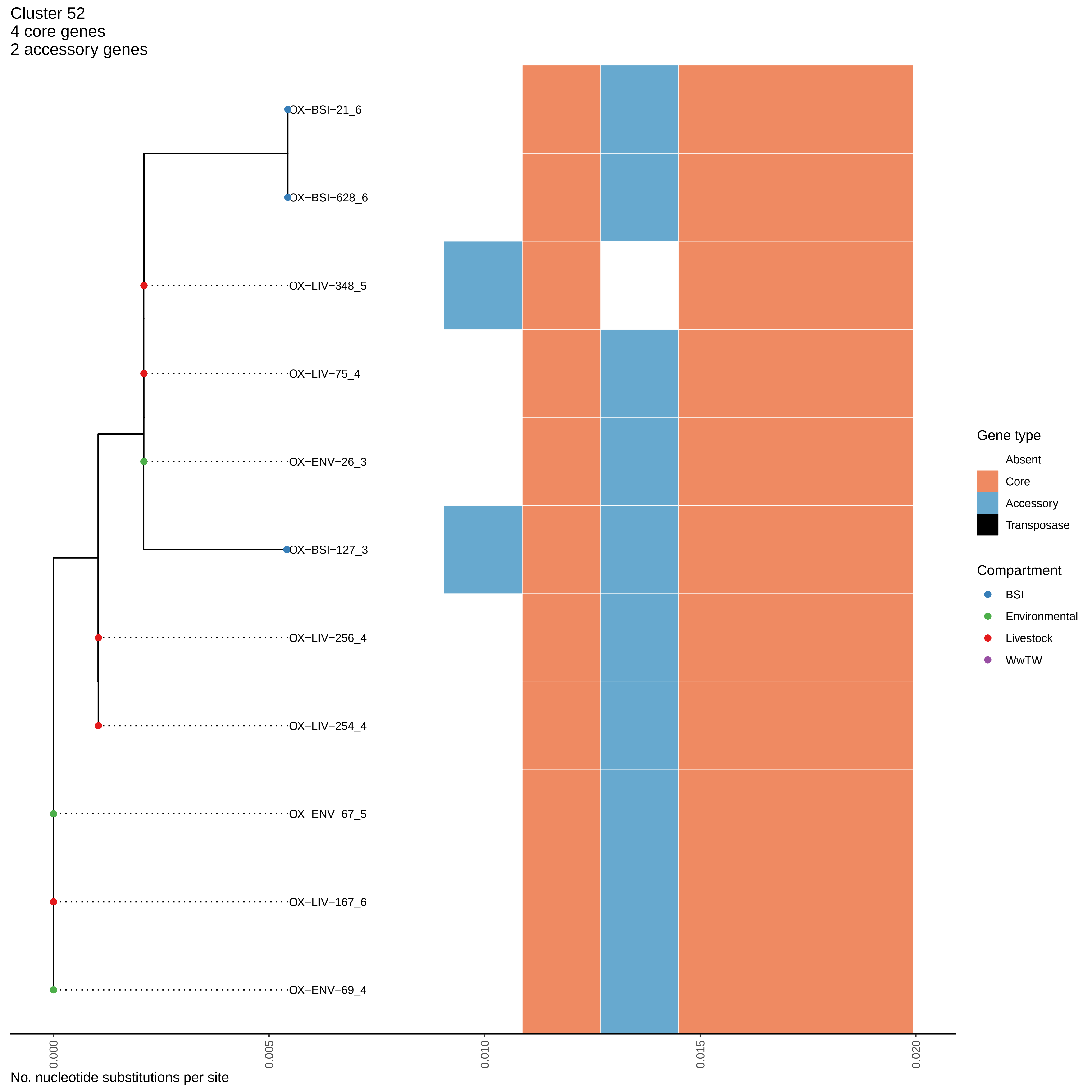
Cluster 52
4 core genes
2 accessory genes
OX−BSI−21_6
OX−BSI−628_6
OX−LIV−348_5
OX−LIV−75_4
OX−ENV−26_3
OX−BSI−127_3
OX−LIV−256_4
OX−LIV−254_4
OX−ENV−67_5
OX−LIV−167_6
OX−ENV−69_4
Gene type
Absent
Core
Accessory
Transposase
Compartment
BSI
Environmental
Livestock
WwTW
0.000
0.005
0.010
0.015
0.020
No. nucleotide substitutions per site

Cluster 53
3 core genes
1 accessory genes

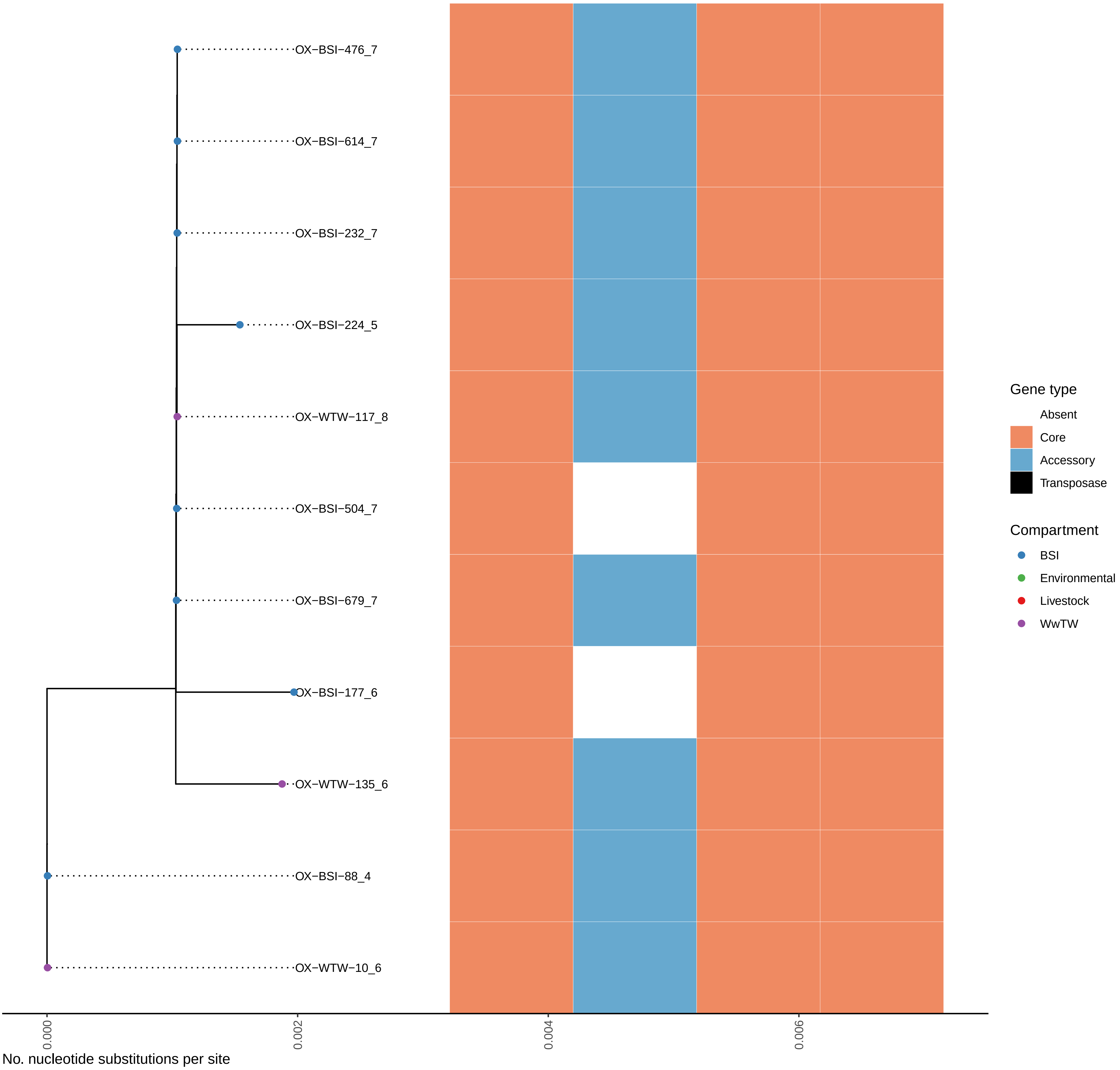

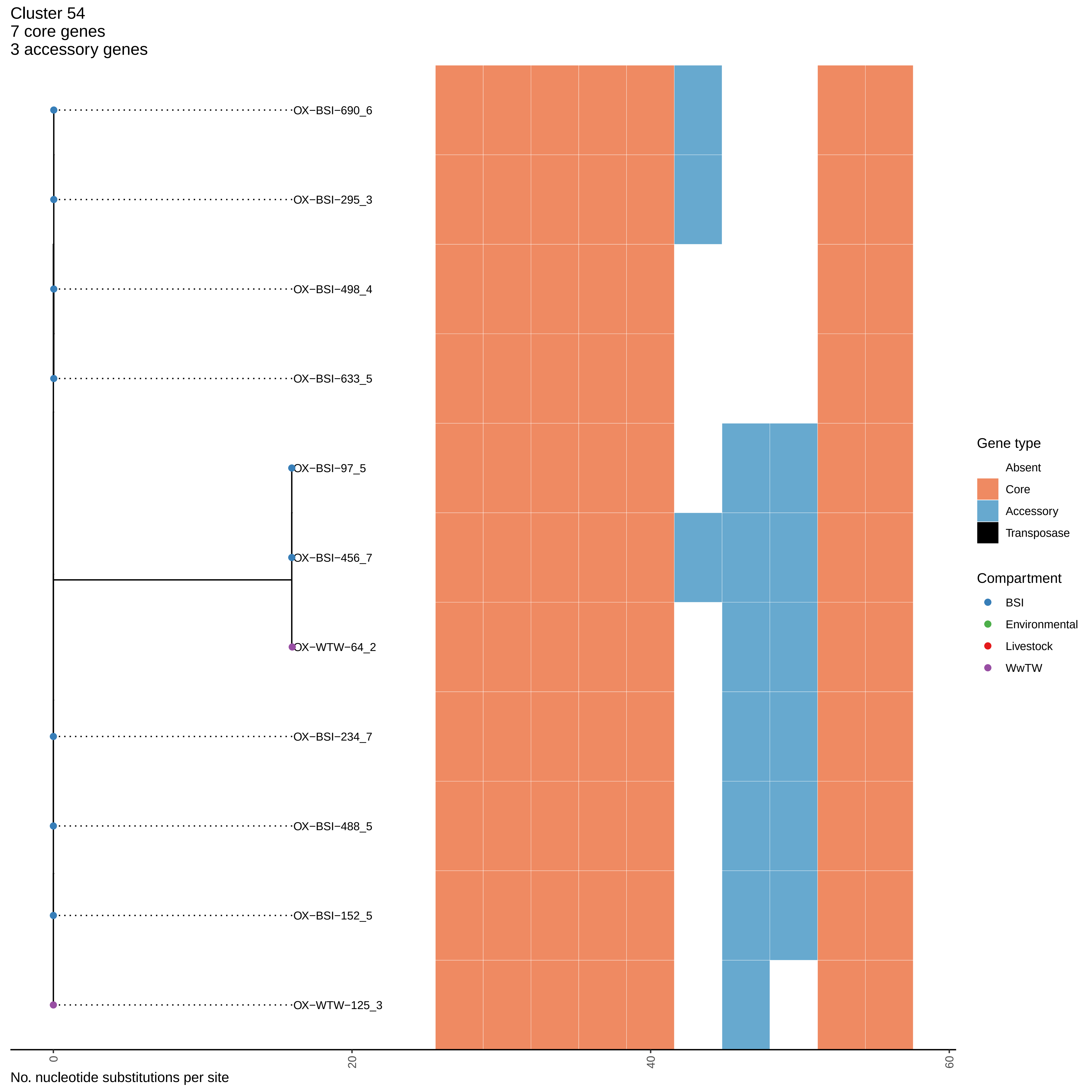
Cluster 54
7 core genes
3 accessory genes
OX−BSI−690_6
OX−BSI−295_3
OX−BSI−498_4
OX−BSI−633_5
OX−BSI−97_5
OX−BSI−456_7
OX−WTW−64_2
OX−BSI−234_7
OX−BSI−488_5
OX−BSI−152_5
OX−WTW−125_3
0
20
40
60
No. nucleotide substitutions per site
Gene type
Absent
Core
Accessory
Transposase
Compartment
BSI
Environmental
Livestock
WwTW

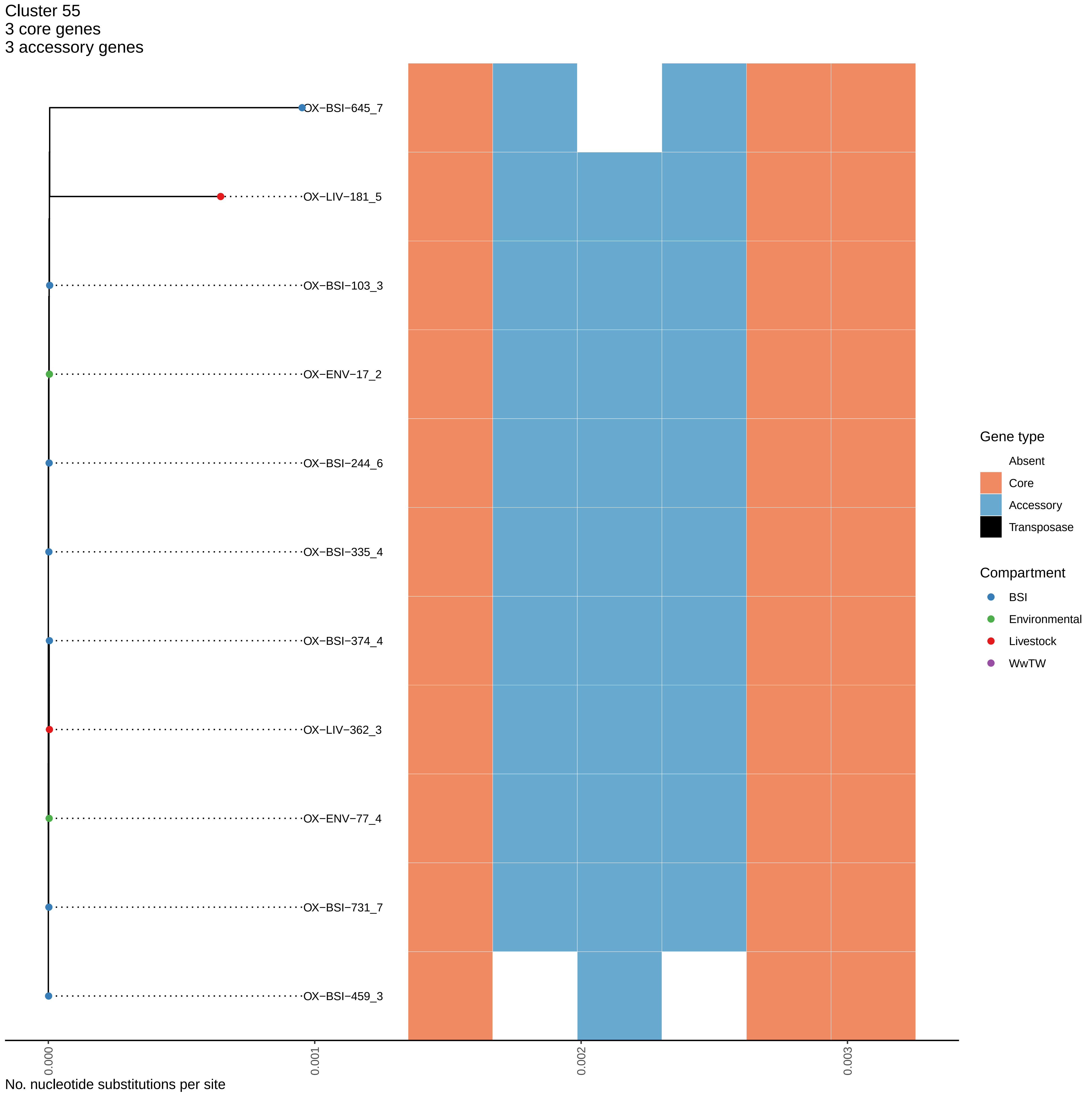
Cluster 55
3 core genes
3 accessory genes
OX−BSI−645_7
OX−LIV−181_5
OX−BSI−103_3
OX−ENV−17_2
OX−BSI−244_6
OX−BSI−335_4
OX−BSI−374_4
OX−LIV−362_3
OX−ENV−77_4
OX−BSI−731_7
OX−BSI−459_3
0.000
0.001
0.002
0.003
No. nucleotide substitutions per site
Gene type
Absent
Core
Accessory
Transposase
Compartment
BSI
Environmental
Livestock
WwTW

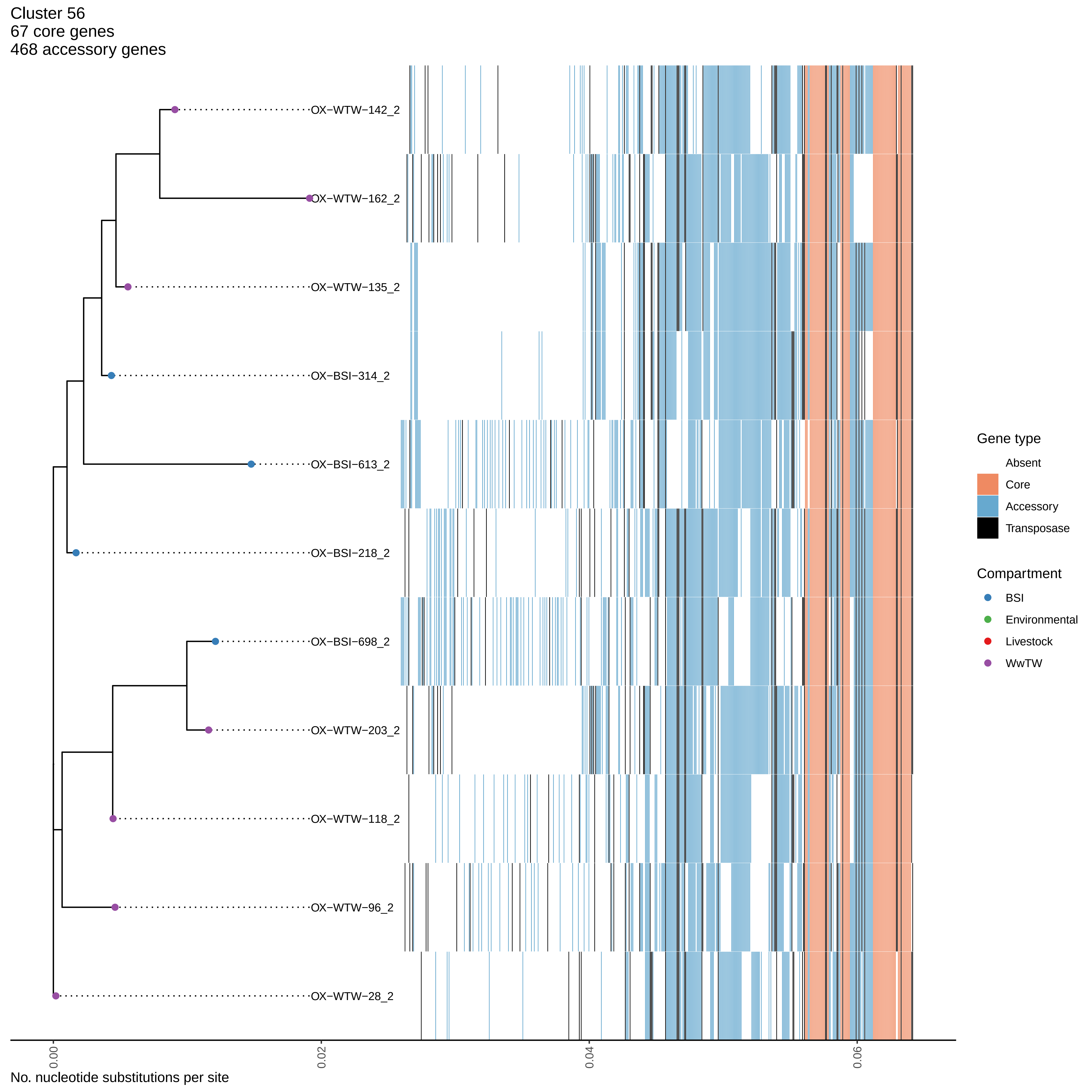

Cluster 56
67 core genes
468 accessory genes
OX−WTW−142_2
OX−WTW−162_2
OX−WTW−135_2
OX−BSI−314_2
OX−BSI−613_2
OX−BSI−218_2
OX−BSI−698_2
OX−WTW−203_2
OX−WTW−118_2
OX−WTW−96_2
OX−WTW−28_2
0.00
0.02
0.04
0.06
No. nucleotide substitutions per site
Gene type
Absent
Core
Accessory
Transposase
Compartment
BSI
Environmental
Livestock
WwTW

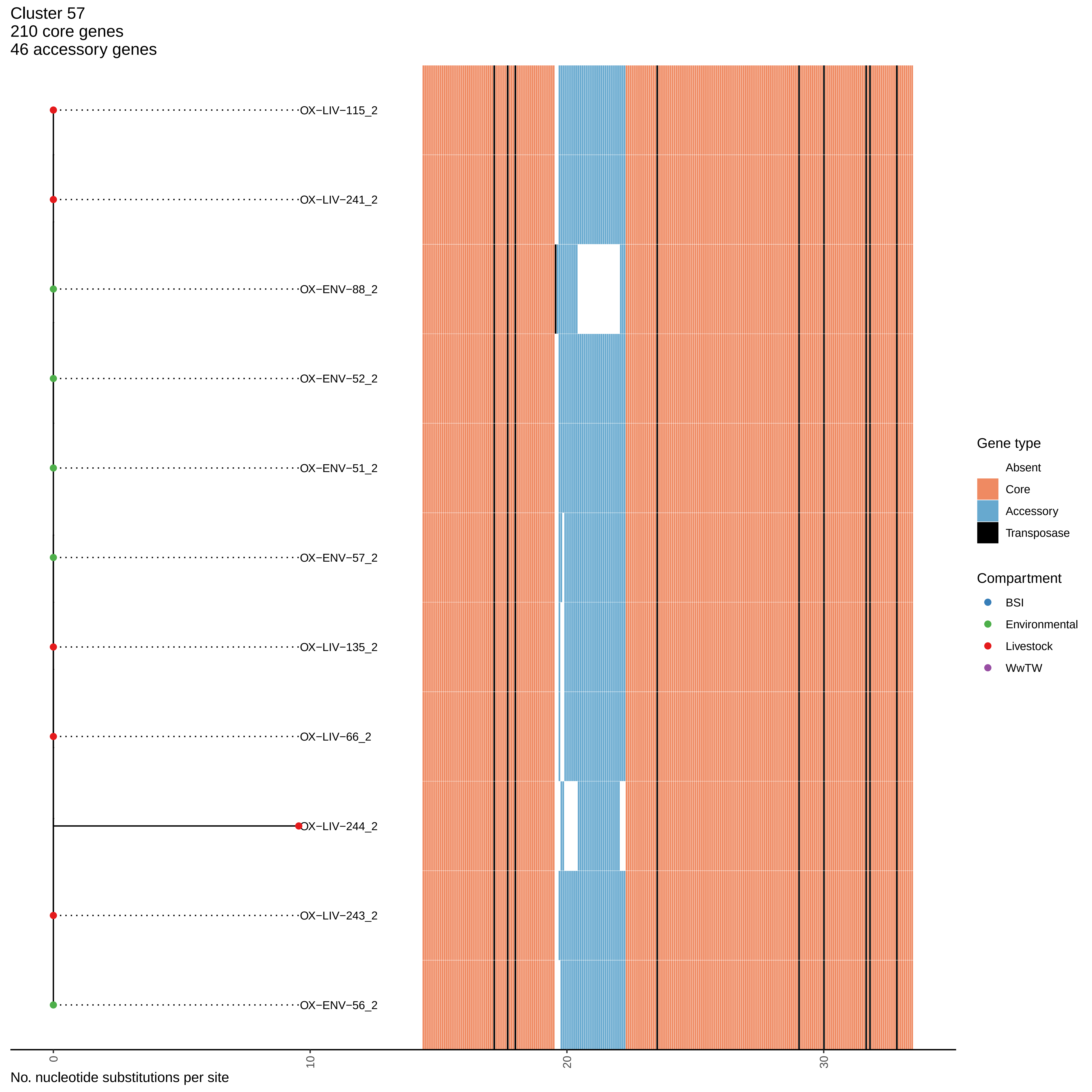

Cluster 57
210 core genes
46 accessory genes
OX−LIV−115_2
OX−LIV−241_2
OX−ENV−88_2
OX−ENV−52_2
OX−ENV−51_2
OX−ENV−57_2
OX−LIV−135_2
OX−LIV−66_2
OX−LIV−244_2
OX−LIV−243_2
OX−ENV−56_2
0
10
20
30
No. nucleotide substitutions per site
Gene type
Absent
Core
Accessory
Transposase
Compartment
BSI
Environmental
Livestock
WwTW

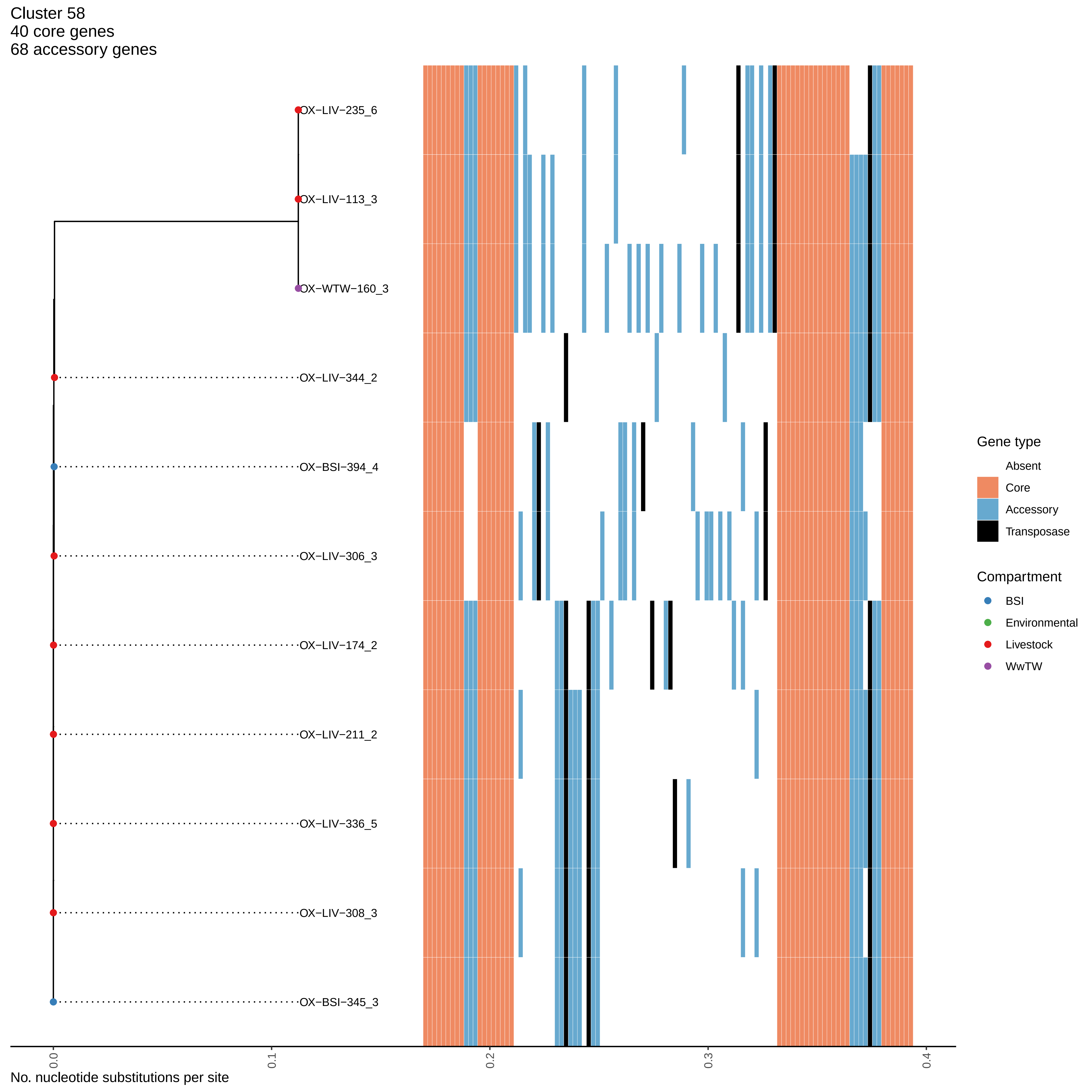

Cluster 58
40 core genes
68 accessory genes
OX−LIV−235_6
OX−LIV−113_3
OX−WTW−160_3
OX−LIV−344_2
OX−BSI−394_4
OX−LIV−306_3
OX−LIV−174_2
OX−LIV−211_2
OX−LIV−336_5
OX−LIV−308_3
OX−BSI−345_3
0.0
0.1
0.2
0.3
0.4
No. nucleotide substitutions per site
Gene type
Absent
Core
Accessory
Transposase
Compartment
BSI
Environmental
Livestock
WwTW

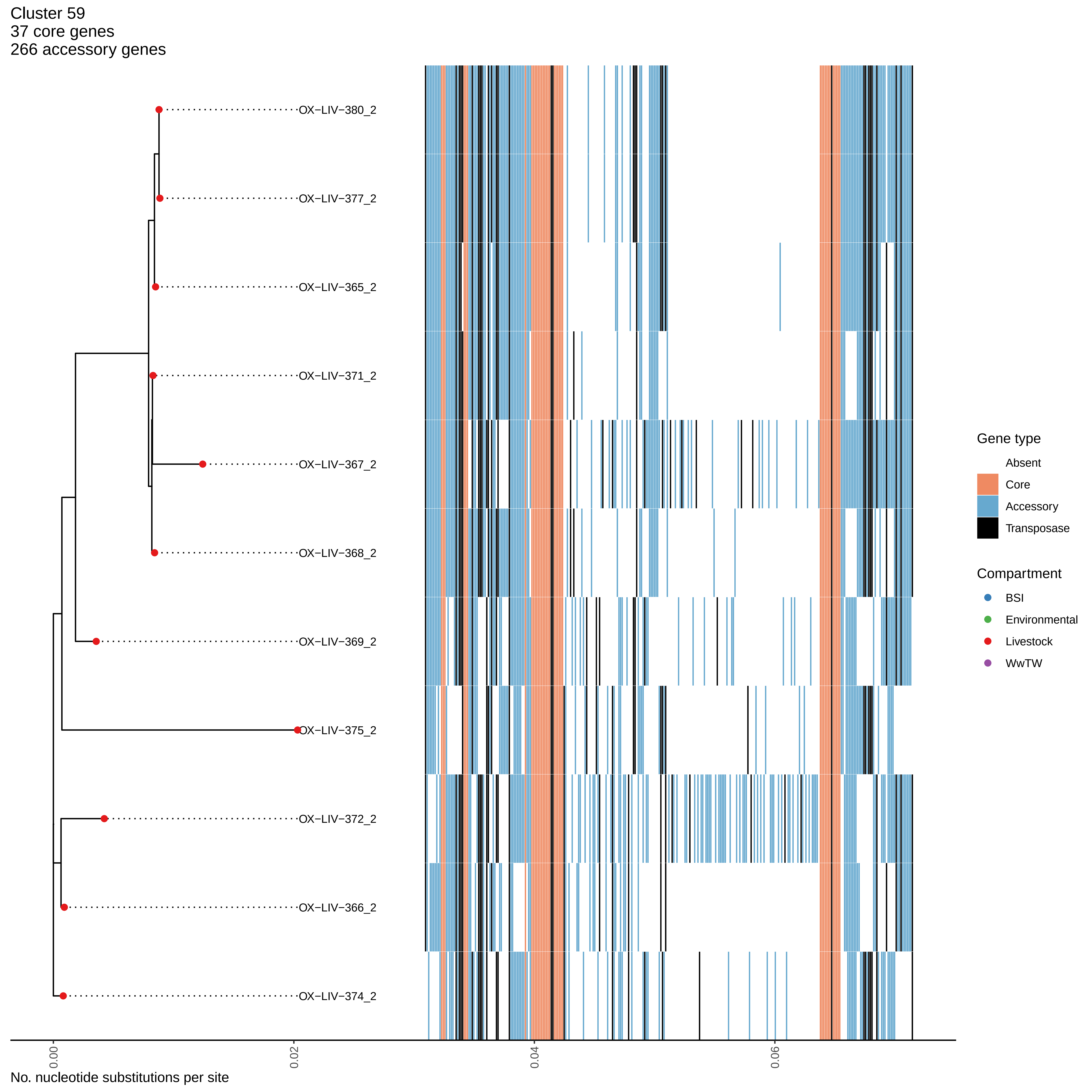
Cluster 59
37 core genes
266 accessory genes
OX−LIV−380_2
OX−LIV−377_2
OX−LIV−365_2
OX−LIV−371_2
OX−LIV−367_2
OX−LIV−368_2
OX−LIV−369_2
OX−LIV−375_2
OX−LIV−372_2
OX−LIV−366_2
OX−LIV−374_2
Gene type
Absent
Core
Accessory
Transposase
Compartment
BSI
Environmental
Livestock
WwTW
0.00
0.02
0.04
0.06
No. nucleotide substitutions per site

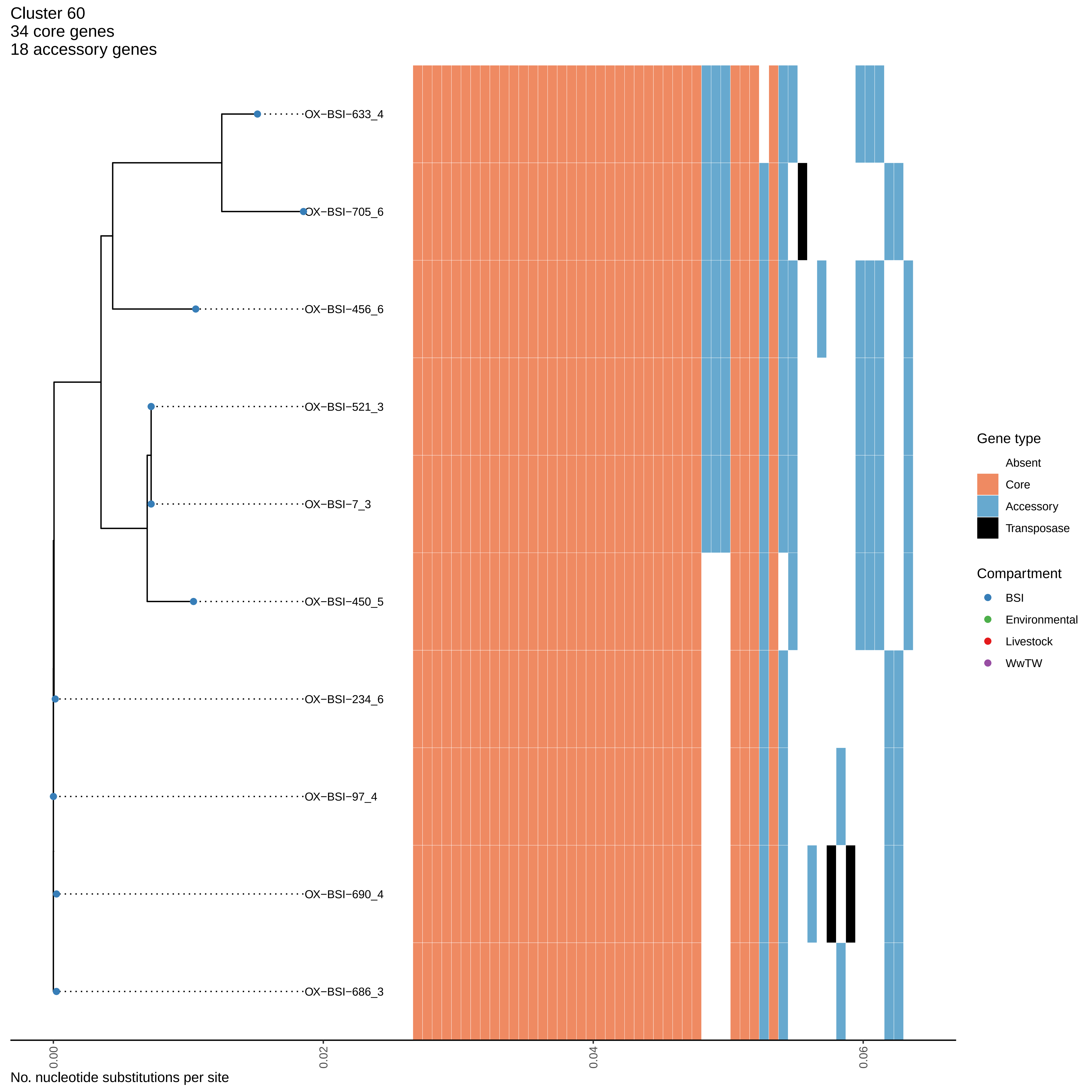

Cluster 60
34 core genes
18 accessory genes
OX−BSI−633_4
OX−BSI−705_6
OX−BSI−456_6
OX−BSI−521_3
OX−BSI−7_3
OX−BSI−450_5
OX−BSI−234_6
OX−BSI−97_4
OX−BSI−690_4
OX−BSI−686_3
0.00
0.02
0.04
0.06
No. nucleotide substitutions per site
Gene type
Absent
Core
Accessory
Transposase
Compartment
BSI
Environmental
Livestock
WwTW

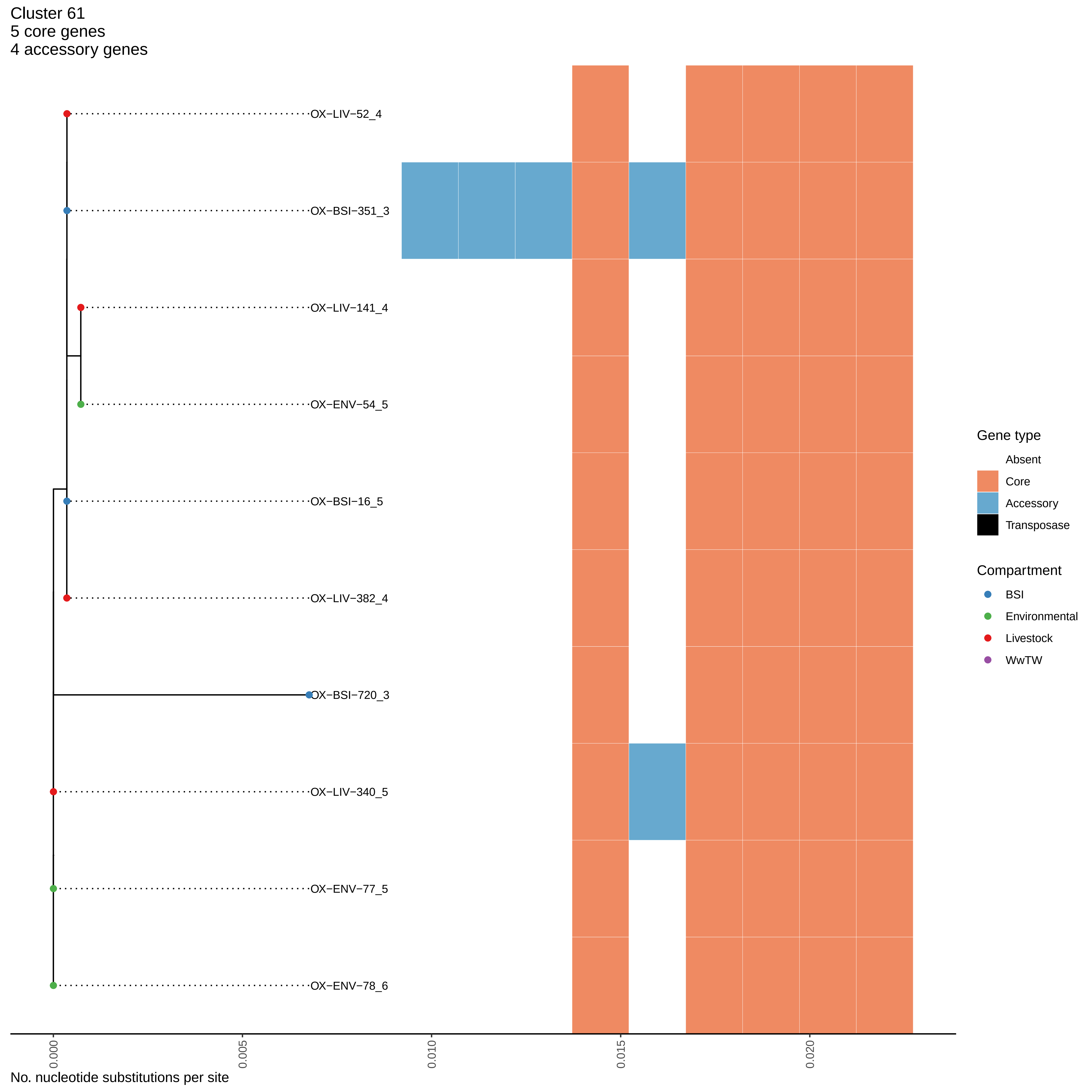
Cluster 61
5 core genes
4 accessory genes
OX−LIV−52_4
OX−BSI−351_3
OX−LIV−141_4
OX−ENV−54_5
OX−BSI−16_5
OX−LIV−382_4
OX−BSI−720_3
OX−LIV−340_5
OX−ENV−77_5
OX−ENV−78_6
0.000
0.005
0.010
0.015
0.020
No. nucleotide substitutions per site
Gene type
Absent
Core
Accessory
Transposase
Compartment
BSI
Environmental
Livestock
WwTW

Cluster 62
4 core genes
0 accessory genes

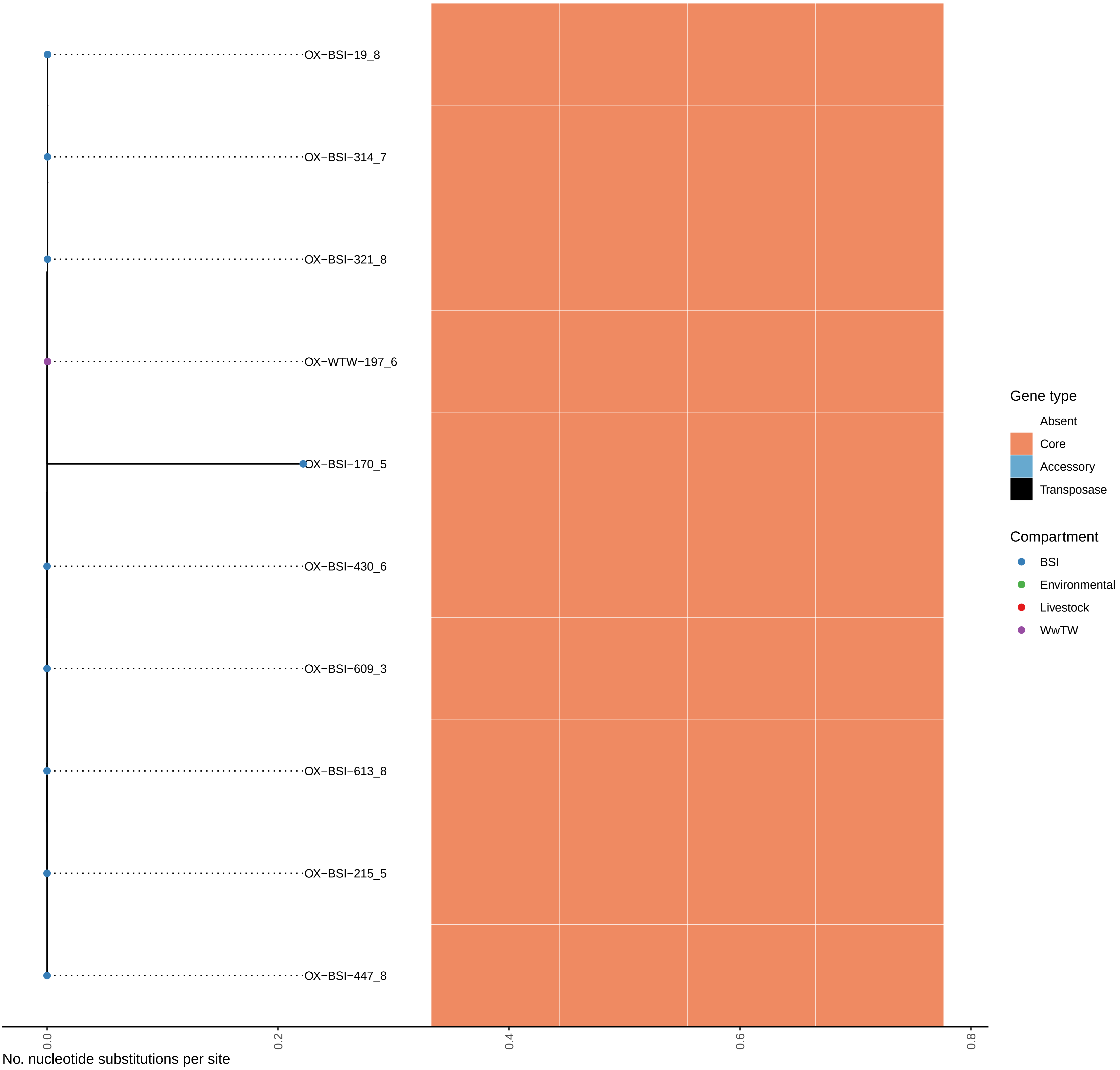

Cluster 63
34 core genes
70 accessory genes

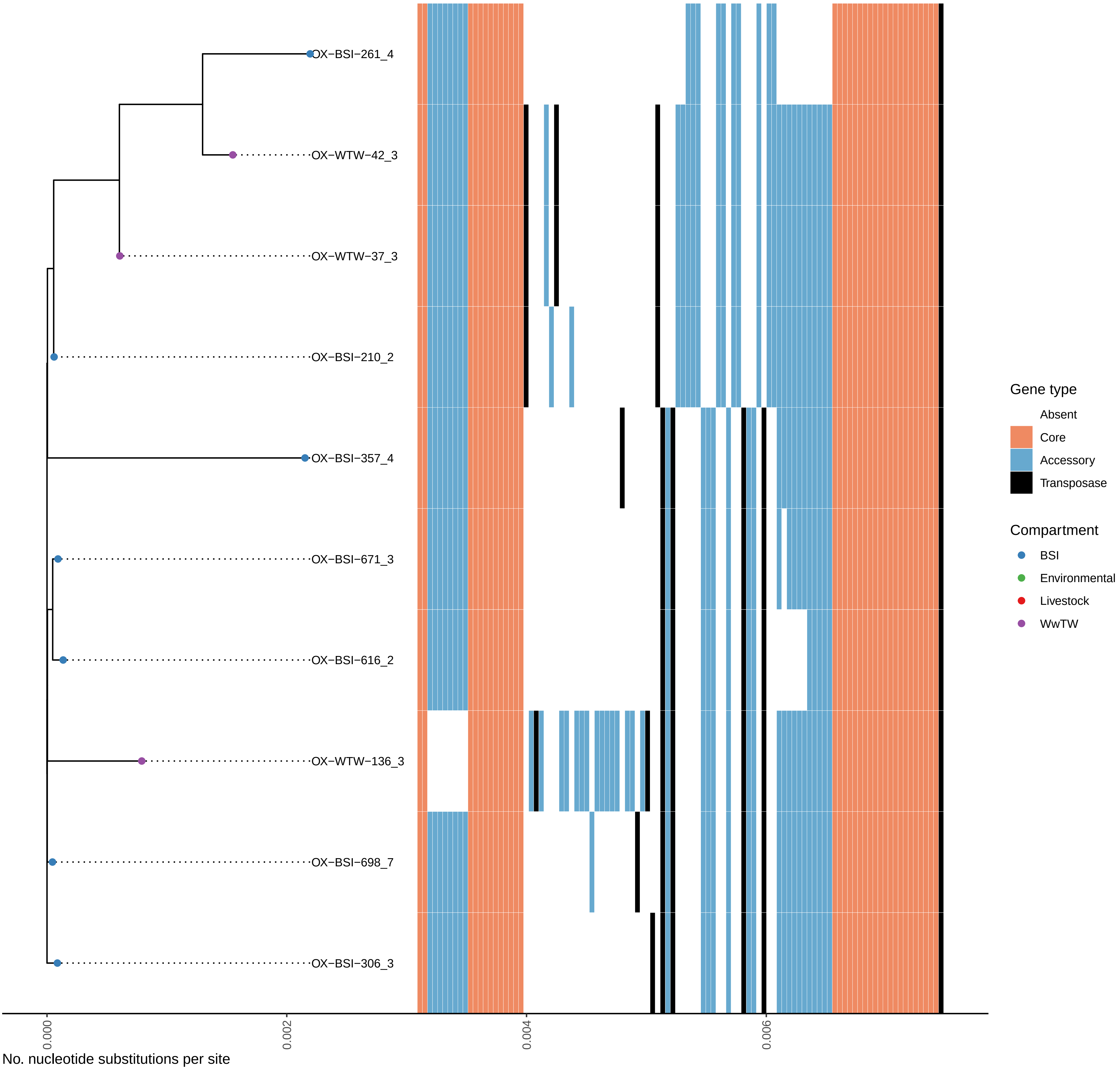

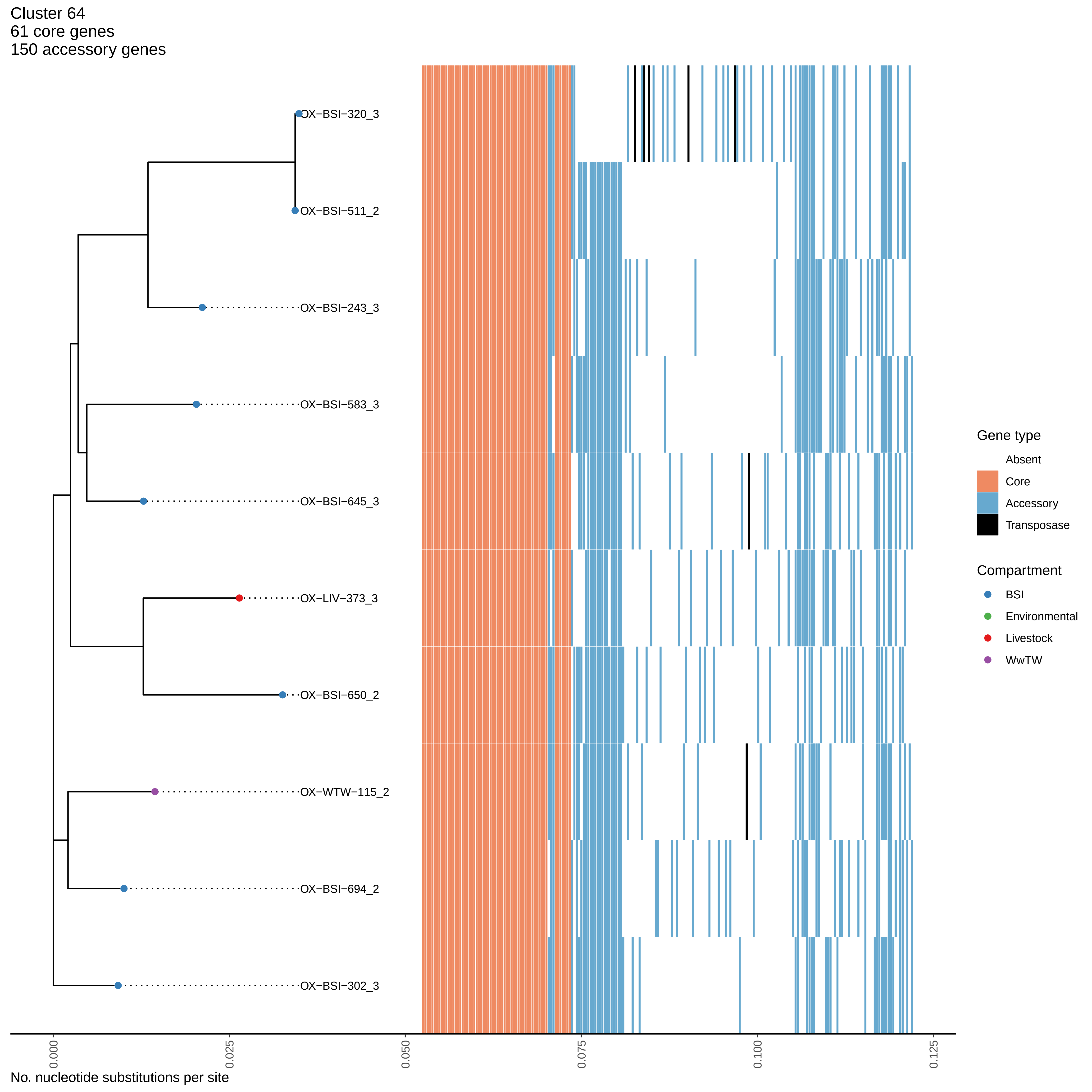

Cluster 64
61 core genes
150 accessory genes
OX−BSI−320_3
OX−BSI−511_2
OX−BSI−243_3
OX−BSI−583_3
OX−BSI−645_3
OX−LIV−373_3
OX−BSI−650_2
OX−WTW−115_2
OX−BSI−694_2
OX−BSI−302_3
0.000
0.025
0.050
0.075
0.100
0.125
No. nucleotide substitutions per site
Gene type
Absent
Core
Accessory
Transposase
Compartment
BSI
Environmental
Livestock
WwTW

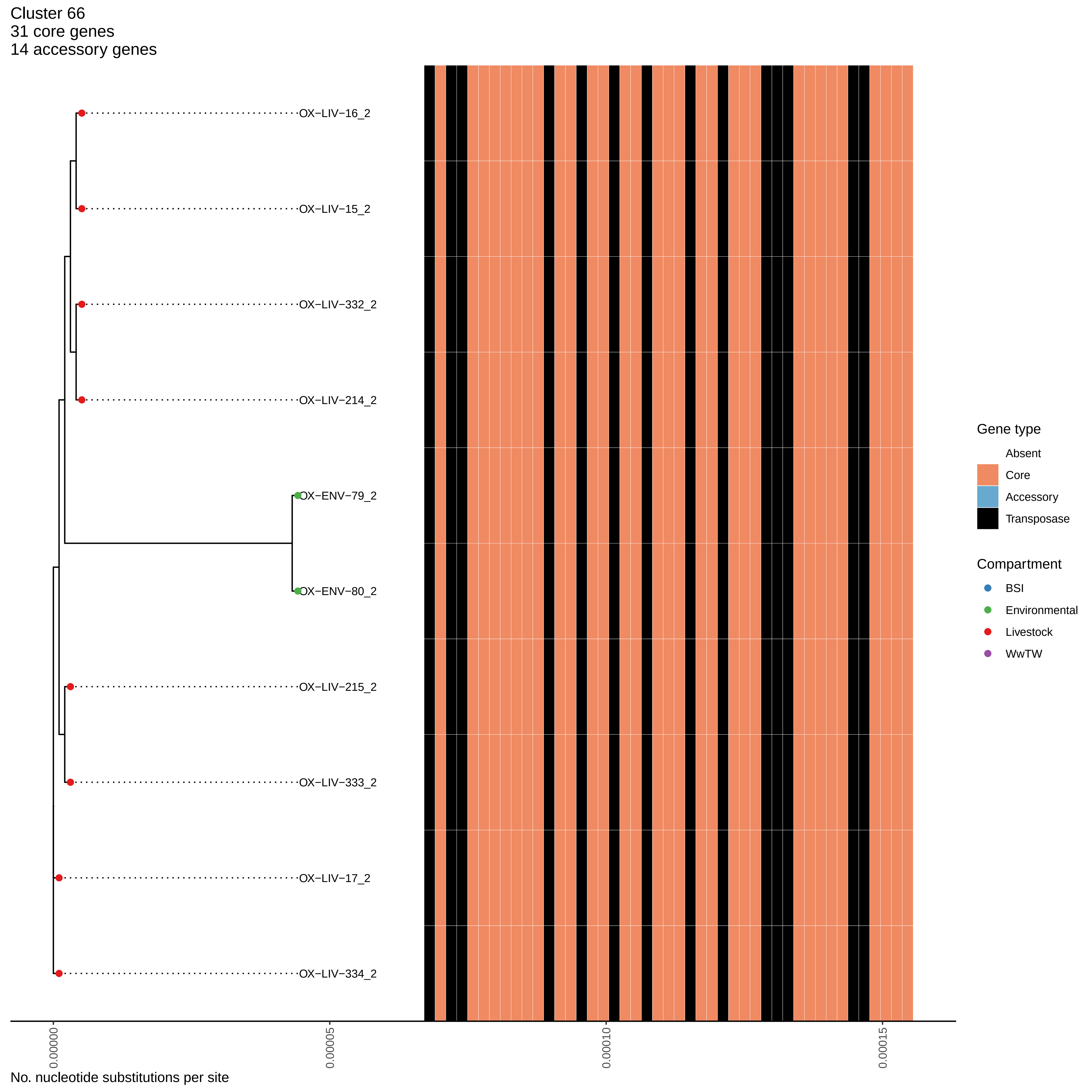

Cluster 66
31 core genes
14 accessory genes
OX−LIV−16_2
OX−LIV−15_2
OX−LIV−332_2
OX−LIV−214_2
OX−ENV−79_2
OX−ENV−80_2
OX−LIV−215_2
OX−LIV−333_2
OX−LIV−17_2
OX−LIV−334_2
Gene type
Absent
Core
Accessory
Transposase
Compartment
BSI
Environmental
Livestock
WwTW
0.00000
0.00005
0.00010
0.00015
No. nucleotide substitutions per site

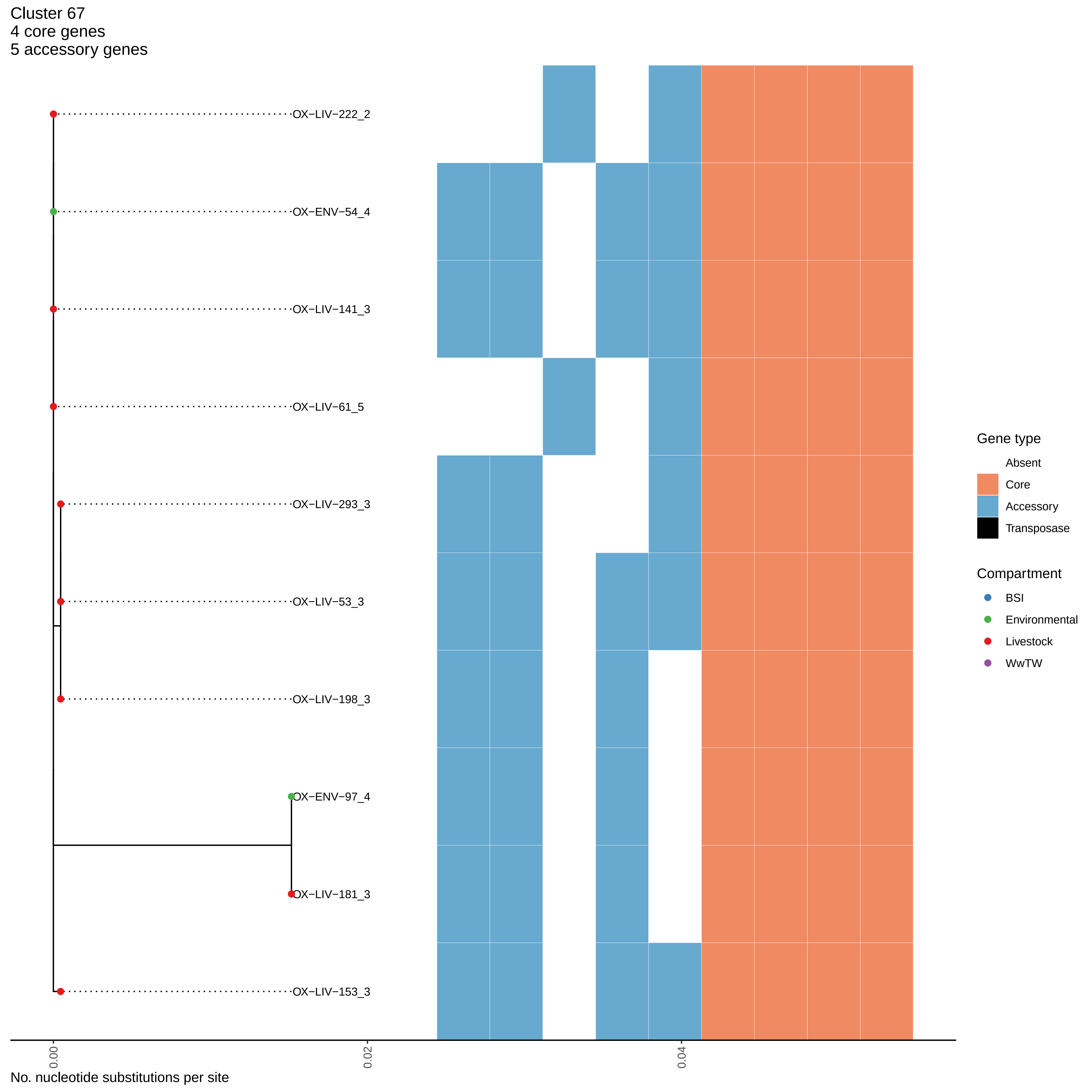

Cluster 67
4 core genes
5 accessory genes
OX−LIV−222_2
OX−ENV−54_4
OX−LIV−141_3
OX−LIV−61_5
OX−LIV−293_3
OX−LIV−53_3
OX−LIV−198_3
OX−ENV−97_4
OX−LIV−181_3
OX−LIV−153_3
0.00
0.02
0.04
No. nucleotide substitutions per site
Gene type
Absent
Core
Accessory
Transposase
Compartment
BSI
Environmental
Livestock
WwTW

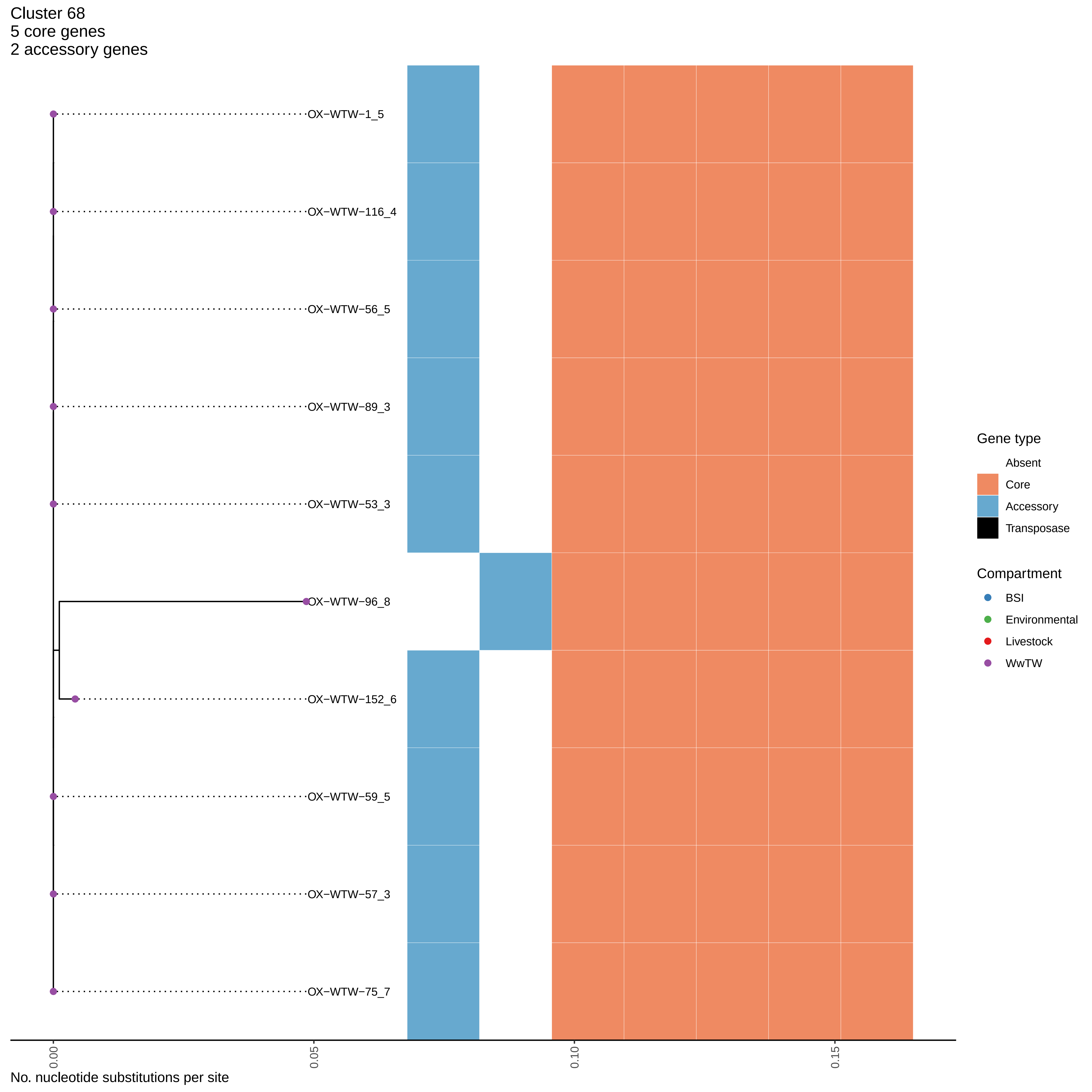
Cluster 68
5 core genes
2 accessory genes
OX−WTW−1_5
OX−WTW−116_4
OX−WTW−56_5
OX−WTW−89_3
OX−WTW−53_3
OX−WTW−96_8
OX−WTW−152_6
OX−WTW−59_5
OX−WTW−57_3
OX−WTW−75_7
0.00
0.05
0.10
0.15
No. nucleotide substitutions per site
Gene type
Absent
Core
Accessory
Transposase
Compartment
BSI
Environmental
Livestock
WwTW

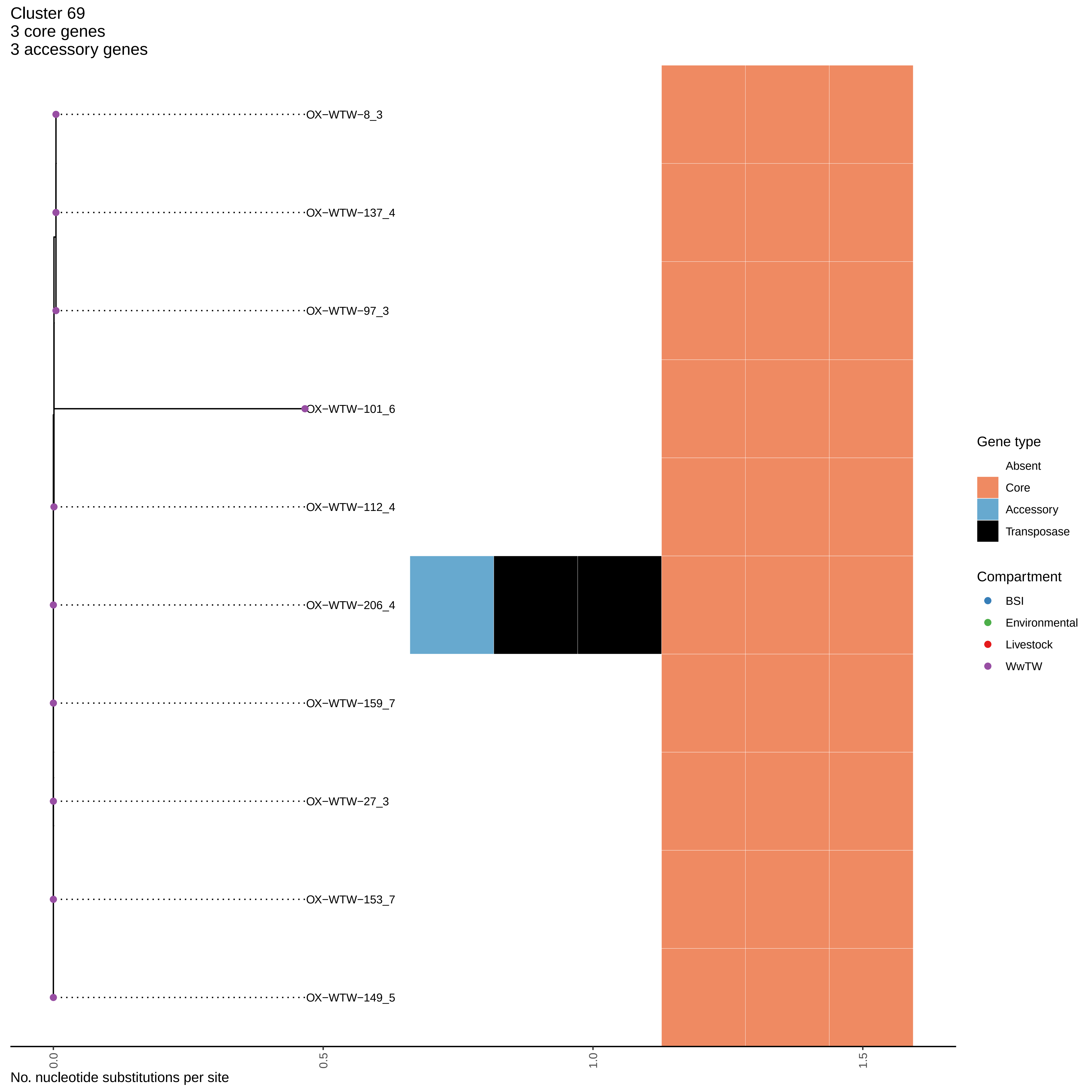
Cluster 69
3 core genes
3 accessory genes
OX−WTW−8_3
OX−WTW−137_4
OX−WTW−97_3
OX−WTW−101_6
OX−WTW−112_4
OX−WTW−206_4
OX−WTW−159_7
OX−WTW−27_3
OX−WTW−153_7
OX−WTW−149_5
0.0
0.5
1.0
1.5
No. nucleotide substitutions per site
Gene type
Absent
Core
Accessory
Transposase
Compartment
BSI
Environmental
Livestock
WwTW
